# Flavodiiron proteins associate pH-dependently with the thylakoid membrane for ferredoxin-1 powered O_2_ photoreduction

**DOI:** 10.1101/2023.05.19.541409

**Authors:** Lauri Nikkanen, Serhii Vakal, Anita Santana-Sánchez, Michal Hubáček, Grzegorz Konert, Yingying Wang, Marko Boehm, Kirstin Gutekunst, Tiina A. Salminen, Yagut Allahverdiyeva

## Abstract

Flavodiiron proteins (FDPs) catalyse light-dependent reduction of oxygen to water in photosynthetic organisms, creating an electron sink on the acceptor side of Photosystem I that protects the photosynthetic apparatus. However, the identity of the electron donor(s) and the molecular mechanisms regulating FDP activity have remained elusive. To elucidate these issues, we employed spectroscopic and gas flux analysis of photosynthetic electron transport, bimolecular fluorescence complementation assays for *in vivo* protein–protein interactions in the model cyanobacterium *Synechocystis* sp. PCC 6803, as well as *in silico* surface charge modelling. We demonstrated that Ferredoxin-1 interacts with Flv1, Flv2, and Flv3, and is the main electron donor to FDP heterooligomers, which are responsible for the photoreduction of oxygen. Moreover, we revealed that association of FDP heterooligomers with thylakoid membranes is promoted by dissipation of the trans-thylakoid proton motive force, providing the first *in vivo* evidence of a self-regulatory feedback mechanism allowing dynamic control of FDP activity and maintenance of photosynthetic redox balance in fluctuating environments. Our findings have direct implications for rationally directing electron flux toward desired reactions in biotechnological applications.

## MAIN TEXT

### Introduction

In oxygenic photosynthetic organisms apart from angiosperms and red and brown algae, class C flavodiiron proteins (FDPs) catalyse light-dependent reduction of O_2_ to H_2_O. This process, referred to as the Mehler-like reaction, creates a strong electron sink on the acceptor side of Photosystem I (PSI) and alleviates excessive reduction of the photosynthetic electron transport chain (PETC), protecting the photosynthetic apparatus from photodamage. It has been proposed that, unlike in the Mehler reaction, no harmful reactive oxygen species (ROS) are produced and O_2_ is reduced directly to H_2_O (Vicente et al. 2002). Indeed, the Mehler-like reaction is essential for the growth of cyanobacteria (Allahverdiyeva et al. 2013), unicellular algae (Chaux et al. 2017; Jokel et al. 2018), and moss (Gerotto et al. 2016) in conditions mimicking natural fluctuations of light intensity. FDPs are also common in non-photosynthetic bacteria, archaea and eukaryotes, and are classified into classes A–H depending on their domain composition (Folgosa et al. 2018). All FDPs share a core modular domain structure containing a metallo *β*-lactamase-like domain with a non-heme diiron catalytic site and a flavodoxin-like domain with a flavin mononucleotide (FMN) binding site. FDPs in photosynthetic organisms typically also harbour a C-terminal flavin-binding domain with predicted NAD(P)H:flavin oxidoreductase activity (Folgosa et al. 2018). Some FDPs, including those in photosynthetic algae, catalyse the reduction of nitric oxide (NO) to nitrous oxide (N_2_O), while yet others can reduce both NO and O_2_ (Folgosa et al. 2018; Burlacot et al. 2020; Blomberg and Ädelroth 2023).

The model cyanobacterium *Synechocystis* sp. PCC 6803 (hereafter *Synechocystis*) has four FDP isoforms: Flv1 (*sll1521*), Flv3 (*sll0550*), Flv2 (*sll0219*), and Flv4 (*sll0217*) (Helman et al. 2003) that likely form various homo- and heterooligomeric conformations. The Mehler-like reaction is catalysed by heterooligomers consisting of Flv1 and Flv3 or Flv2 and Flv4 (Helman et al. 2003; Allahverdiyeva et al. 2013; Mustila et al. 2016; Santana-Sanchez et al. 2019). Although it has been established that FDPs function as a release valve for excessive electrons on the acceptor side of PSI (Helman et al. 2003; Allahverdiyeva et al. 2013; Santana-Sanchez et al. 2019), and that the Mehler-like reaction is dependent on the presence of all four FDPs under air-level CO_2_ (Mustila et al. 2016; Santana-Sanchez et al. 2019), the identity of the electron donor(s) to FDPs has remained uncertain. Based on the presence of the C-terminal NAD(P)H: flavin oxidoreductase-like domain as well as *in vitro* experiments with recombinant proteins, it was suggested that NAD(P)H would function as the main electron donor to Flv1, Flv3, and Flv4 (Vicente et al. 2002; Shimakawa et al. 2015; Brown et al. 2019). However, in those experiments FDPs were present as homooligomers, whose physiological function remains unknown but has been shown to be distinct from O_2_ photoreduction (Mustila et al. 2016). Given the distinct kinetic profiles of O_2_ photoreduction catalysed by Flv1/3 and Flv2/4 heterooligomers under air-level CO_2_ concentrations (Santana-Sanchez et al. 2019), the *in vivo* electron donor to different heterooligomers, and to the unknown reactions catalysed by FDP homooligomers, could differ. Flv1, Flv2, Flv3, and Flv4 homodimers, Flv2/4 heterodimers have been biochemically detected in *Synechocystis,* and formation of tetramers of recombinant Flv3 in *E. coli* has been observed (Allahverdiyeva et al. 2011; Zhang et al. 2012; Mustila et al. 2016). However, Flv1/3 heterooligomers have not been directly observed, and the abilities of FDPs to interact with each other and to form different heterodimers and higher-order oligomers have not been fully explored.

Alternative electron donors such as ferredoxin (Fed) or ferredoxin:NADP^+^ oxidoreductase (FNR) were not tested in the aforementioned *in vitro* studies (Vicente et al. 2002; Brown et al. 2019). There are two FNR isoforms in *Synechocystis*. The larger FNR_L_ form functions as the main photosynthetic carrier of electrons from Fed to NADP^+^, while the smaller FNR_S_ form, deriving from later initiation of transcription of the *petH* gene, is upregulated under nutrient starvation or in heterotrophic conditions (Thomas et al. 2006). FNR_S_ was recently suggested to catalyse NADPH oxidation during cyclic electron transport (CET) upon dark-light transitions, providing reduced Fed for NDH-1 (Miller et al. 2022). However, recent *in vivo* measurements of light-induced NAD(P)H and Fed redox changes in wild-type (WT) *Synechocystis* and Flv1/3-deficient mutant cells support Fed as the main electron donor at least to Flv1/3 heterooligomers (Nikkanen et al. 2020; Sétif et al. 2020). *Synechocystis* has eleven Fed isoforms (Fed1-Fed11) (Artz et al. 2020) but only little is known about their specific functions or the extent of redundancy between them. Fed1–6, as well as Fed10–11 are “plant-type” ferredoxins harbouring a [2Fe-2S] cluster, while Fed7, 8, and 9 are bacterial types with [4Fe-4S], [3Fe-4S] [4Fe- 4S], and [4Fe-4S] [4Fe-4S] iron-sulphur cluster compositions, respectively (Cassier-Chauvat and Chauvat 2014; Artz et al. 2020; Wang et al. 2022). Previously, Fed9 has indeed been shown to interact with Flv3 in a bacterial two-hybrid system (Cassier-Chauvat and Chauvat 2014), and Flv1 and Flv3 with Fed1 in a Fed-affinity chromatographic assay (Hanke et al. 2011). Moreover, the Δ*Fed7* mutant has a lowered expression level of the *flv4-2* operon and decreased photosynthetic efficiency under high irradiance and air-level [CO_2_] (Mustila et al. 2014). Evidence of *in vivo* interaction is however lacking.

To resolve these ambiguities, we developed a bimolecular fluorescence complementation (BiFC) platform to study protein–protein interactions in living cyanobacterial cells, which we then utilised to examine the capability of FDPs to interact *in vivo* with putative electron donors as well as with each other. BiFC takes advantage of reassembly of a fluorescent protein (FP) upon proximity of two co-expressed fusion proteins consisting of a protein of interest and an N- or C-terminal fragment of an FP (Hu et al. 2002; Kerppola 2008). The FP is then excited at the appropriate wavelength, and the resulting fluorescence emission from interactions is typically detected by a confocal microscope. A substantial benefit of BiFC tests in comparison to *in vitro* methods such as co-immunoprecipitation, or methods based on exogenous expression such as two-hybrid systems, is that it can also provide information on the *in vivo* subcellular localisation of the detected protein–protein interactions. Although BiFC has been extensively used to study interactions in plant cells and chloroplasts, as well as in other bacterial species (Walter et al. 2004; Kerppola 2008; Kudla and Bock 2016; Nikkanen et al. 2016), it has not been previously utilised in cyanobacteria. Background fluorescence caused by cyanobacterial phycobilisomes or chlorophyll (Chl) does not, however, interfere with the use of fluorophores such as GFP, YFP or CFP (Yokoo et al. 2015).

To reveal if any of the low abundance Fed isoforms in *Synechocystis* (Fed2-Fed11), NADPH, or FNR are required for the Mehler-like reaction, we performed biophysical characterisation of deletion strains deficient in low abundance Feds and FNR. We then used the BiFC platform to elucidate the protein– protein interactions and subcellular localisations of different FDP isoforms. Our combined approach of biophysical characterisation and BiFC assays allowed us to demonstrate that Fed1, not NADPH or FNR, is likely to be the main electron donor to the light-induced Mehler-like reaction catalysed by Flv1/3 and Flv2/4 heterooligomers. Moreover, the subcellular localisation of the interactions between FDPs and between FDPs and Fed1 provided evidence that the magnitude of the trans-thylakoid proton gradient controls reversible association of different Flv1/3 and Flv2/4 heterooligomers with the thylakoid membrane. *In silico* modelling of FDP heterooligomer surface charges supported pH-dependent electrostatic binding to thylakoid membranes. Based on these findings we propose a novel self-feedback mechanism for precise post-translational regulation of Fed1-driven O_2_ photoreduction activity of FDP heterooligomers.

## Results

### Flv3 interacts with FNR on thylakoids but FNR and NADPH are not main reductants of Flv1/3 heterooligomers in the Mehler-like reaction

First, we investigated the possibility of FDPs interacting directly with, and/or being reduced by FNR. To that end, we monitored real-time NADPH redox changes and light-induced O_2_ exchange in knockout mutant strains ΔFNR_L_ and ΔFNR_S_ (also called FSI and MI6, respectively) (Thomas et al. 2006). Measurement of NADPH fluorescence revealed that light-dependent NADP^+^ reduction was impaired in ΔFNR_L_ (Fig. 1A–B), in agreement with the previous observation that NADPH/NADP^+^ ratio is lowered in the mutant (Korn et al. 2009). No difference to WT was detected in ΔFNR_S_.

**Fig. 1.**
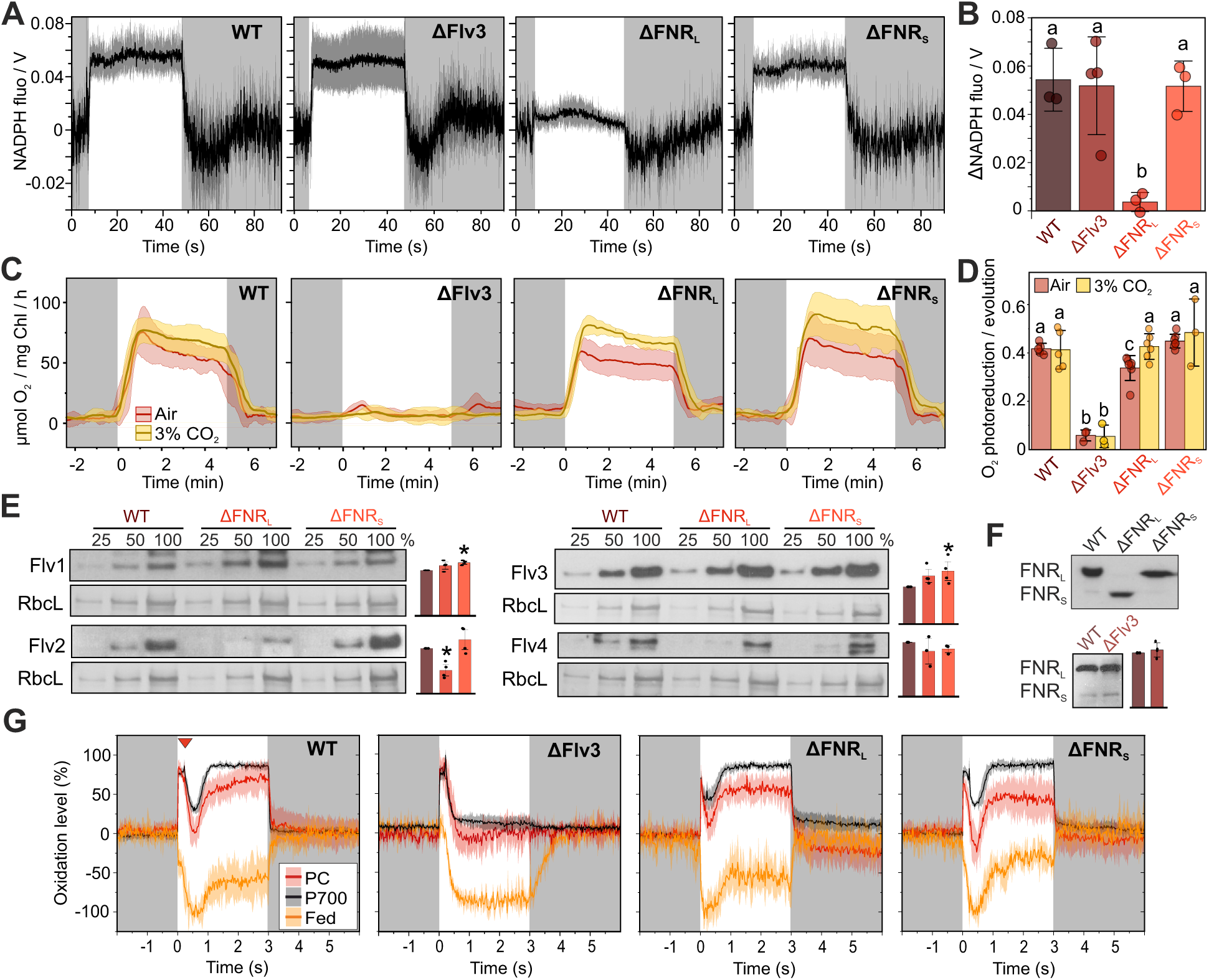
Photosynthetic characterisation of FNR mutants and protein-protein interaction tests between FNR_L_ and FDPs. **(A)** Light-induced NADPH fluorescence changes in WT, ΔFlv3, ΔFNR_L_ (no FNR_L_, increased amount of FNR_S_), and ΔFNR_S_ (no FNR_S_, FNR_L_ present) cells during 40 s illumination at 500 µmol photons m^-2^ s^-1^ and subsequent darkness. Averaged traces from three (WT, ΔFNR_L,_ ΔFNR_S_) or four (ΔFlv3) biological replicates +/- standard deviation (shadowed area) are shown. Dark-adapted NADPH fluorescence levels were set to 0. **(B)** Quantification of light-induced change in the NADPH fluorescence signal in the experiments for (A), calculated as mean fluorescence level between 30 and 40 sec of illumination – mean fluorescence level in the pre-illumination dark period. This timeframe was chosen to coincide with the maximal activity of FDPs. Values are averages from three (WT, ΔFNR_L,_ ΔFNR_S_) or four (ΔFlv3) biological replicates +/- standard deviation with individual data points shown as circles. Statistical significance was tested by one-way ANOVA and a Tukey’s post-hoc test for comparison of means. Means that do not share a grouping letter (a, b) are significantly different (P<0.05). **(C)** O_2_ uptake kinetics measured by MIMS in WT, ΔFlv3, ΔFNR_L_, and ΔFNR_S_ cells during 5 min illumination at 500 µmol photons m^-2^ s^-1^ and subsequent darkness. ^18^O_2_ was added prior to the measurement in equal concentration to ^16^O_2_ in order to distinguish O_2_ uptake from O_2_ evolution. Averaged traces from 5 (WT), 3 (ΔFlv3), and 7 (ΔFNR_L_ and ΔFNR_S_) biological replicates +/- standard deviation (shadowed area) are shown in red from cultures grown under air-level [CO_2_], and in yellow from 5 (WT), 6 (ΔFNR_L_), and 3 (ΔFlv3 and ΔFNR_S_) replicates grown under 3% [CO_2_]. **(D)** The average magnitudes of the maximum O_2_ photoreduction peaks normalised to the maximum O_2_ evolution rate. Averages from 5 (WT), 3 (ΔFlv3), and 7 (ΔFNR_L_ and ΔFNR_S_) biological replicates +/- standard deviation are shown in red from cultures grown under air-level [CO_2_], and in yellow from 5 (WT), 6 (ΔFNR_L_), and 3 (ΔFlv3 and ΔFNR_S_) replicates grown under 3% [CO_2_]. Individual data points shown are as circles. Statistical significance was tested by one-way ANOVA and Tukey’s post-hoc tests for comparison of means. Comparisons were made within the air-level and 3% CO_2_ sets of samples. **(E)** Immunodetection of Flv1, Flv2, Flv3, and Flv4 protein content in WT, ΔFNR_L_, and ΔFNR_S_ cells using specific antibodies. Representative blots and Coomassie brilliant blue-stained band of the large subunit of Rubisco (RbcL, as loading control) from three biological replicates are shown, with bar charts showing quantification of band intensity ± SD, individual data points shown as black dots. * denotes statistically significant difference to WT control according to one-sample Student’s T-tests. Total proteins were extracted from cultures grown under air-level [CO_2_] conditions. **(F)** Immunodetection of FNR protein content in WT, ΔFlv3, ΔFNR_L_, and ΔFNR_S_ cells using specific antibodies. Representative blots from four biological replicates are shown, with the bar chart showing quantification of FNR_L_ band intensity in WT vs ΔFlv3 cells ± SD. Individual data points are shown as black dots. Difference to WT control was found statistically insignificant (P>0.05) by a one-sample Student’s T-test. **(G)** Changes of PC, P700, and Fed in WT, ΔFlv3, ΔFNR_L_ and ΔFNR_S_ cells, as measured with a DUAL-KLAS-NIR spectrophotometer. The traces are normalised to the maximal oxidation values of PC and P700 and maximal reduction of Fed, as determined with the NIRMAX protocol, of which a 3 s illumination with actinic light is shown here, including a multiple turnover pulse after 200 ms (indicated by the red triangle in the WT panel). Averaged traces from 3 (ΔFlv3) or 5 (WT, ΔFNR_L_, and ΔFNR_S_) biological replicates are shown, with standard deviation as the shadowed area.

In WT cells, a strong but transient peak of O_2_ photoreduction during the first minute after the onset of illumination is catalysed mainly by Flv1/3 heterooligomers with some contribution from Flv2/4, while the steadier O_2_ photoreduction thereafter is mostly dependent on Flv2/4 (Santana-Sanchez et al. 2019) (Fig. 1C). If NADPH was the electron donor to FDP heterooligomers, the shortage of light-induced NADPH production in ΔFNR_L_ cells should result in inhibition of the Mehler-like reaction. However, no major impairment of maximal O_2_ photoreduction rate was detected in either ΔFNR_L_ or ΔFNR_S_, when O_2_ fluxes were monitored by time-resolved membrane-inlet mass spectrometry (MIMS) (Fig. 1C–D). A small decrease in maximum O_2_ photoreduction was, however, observed in ΔFNR_L_. This was likely caused by a diminished amount of Flv2 protein in the mutant (Fig. 1E), as lack of Flv2/4 does partially impair the Mehler-like reaction under air level [CO_2_] (Santana-Sanchez et al. 2019). Conversely, slightly elevated protein levels of Flv1 and Flv3 were detected in both FNR mutants, albeit significantly only in ΔFNR_S_ (Fig. 1E). In order to confirm whether the lower Flv2/4 content is the cause for the decreased O_2_ photoreduction in ΔFNR_L_, we repeated the MIMS experiment under elevated (3%) [CO_2_] conditions, where the *flv4-2* operon is not expressed at all (Zhang et al. 2009). Indeed, no difference in the O_2_ photoreduction rate was observed between WT and ΔFNR_L_ cells grown in 3% [CO_2_] (Fig. 1C–D, Fig S1D). Despite being slightly higher under elevated [CO_2_] compared to ambient levels, the light-induced accumulation of NADPH was notably impaired in ΔFNR_L_ cells under both conditions, while no difference to WT was observed in ΔFNR_S_ (Fig. S2A–B). In low-light conditions, where activity of the Mehler-like reaction is low (Allahverdiyeva et al. 2013, Ortega-Martínez et al. 2024), accumulation of NADPH was also diminished in ΔFNR_L_ compared to WT (Fig. S2C–D).

An elevated level of dark respiration was detected in air-grown ΔFNR_L_ cells (Fig. S1C), possibly due to the increased amount of FNR_S_ (Fig. 1F) (Thomas et al. 2006). No significant differences in maximal gross O_2_ evolution (Fig. S1B) or steady state CO_2_ fixation rate (Fig. S1E) were detected between the FNR mutants and WT.

Deficiency of Flv1/3 also results in a distinct inability to re-oxidise Fed and P700 during the first seconds at the onset of strong illumination (Nikkanen et al. 2020; Sétif et al. 2020; Theune et al. 2021). We therefore used the DUAL-KLAS-NIR spectrometer to probe the redox kinetics of the PSI electron donor plastocyanin (PC), P700, and Fed upon onset of strong illumination. Whereas ΔFlv3 cells are unable to re-oxidise Fed, PC, and P700 following initial reduction, both ΔFNR_L_ and ΔFNR_S_ showed similar re-oxidation kinetics of the electron carriers to WT in the seconds-timescale (Fig. 1G), suggesting robust FDP activity. However, full reduction of the Fed pool was reached already after c.a. 100 ms in the ΔFNR_L_ mutant, while in WT and ΔFNR_S_ full reduction took c.a. 400 ms and in ΔFlv3 as long as 600 ms of strong illumination (Fig. S3A). This means that electrons do initially pile up at Fed when FNR_L_ is missing, but once FDPs are activated at roughly 500 ms (Nikkanen et al. 2020) that congestion is effectively alleviated. Moreover, oxidation of P700 under far-red light was delayed in ΔFNR_L_ (Fig. S3B), suggesting increased CET in the mutant.

We also investigated potential protein–protein interactions between FNR_L_ and FDPs using the BiFC system. When Flv3 fused to an N-terminal Venus (an enhanced YFP variant) fragment (Flv3-VN) was co-expressed in WT *Synechocystis* cells with FNR_L_ fused to a C-terminal Venus fragment (FNR_L_-VC), strong fluorescence emission at 519–570 nm was detected upon excitation at 488 nm, indicating successful reassembly of the Venus fragments and interaction between Flv3 and FNR_L_ (Fig. 2A). In order to assess the subcellular localisation of the interaction, we compared the intensity of Chl *a* fluorescence, indicative of the location of the thylakoid membranes, with the fluorescence from re-assembled Venus proteins, over cross-sections of individual cells. As thylakoid membranes are mostly located along the edges of the cell in *Synechocystis*, the Chl regression line forms an M-shape over cell cross-sections. We also plotted the co-localised Chl and Venus fluorescence intensities against each other in a cytofluorograph and calculated the Pearson correlation coefficient (PCC) of Chl and Venus fluorescence co-localisation. Venus fluorescence from FNR_L_-Flv3 interactions along cell cross sections closely matched that of Chl and showed an average PCC of 0.66±0.17 (Fig. 2A). These analyses indicated that the FNR_L_-Flv3 interactions localised consistently to the thylakoid membrane.

**Fig. 2.**
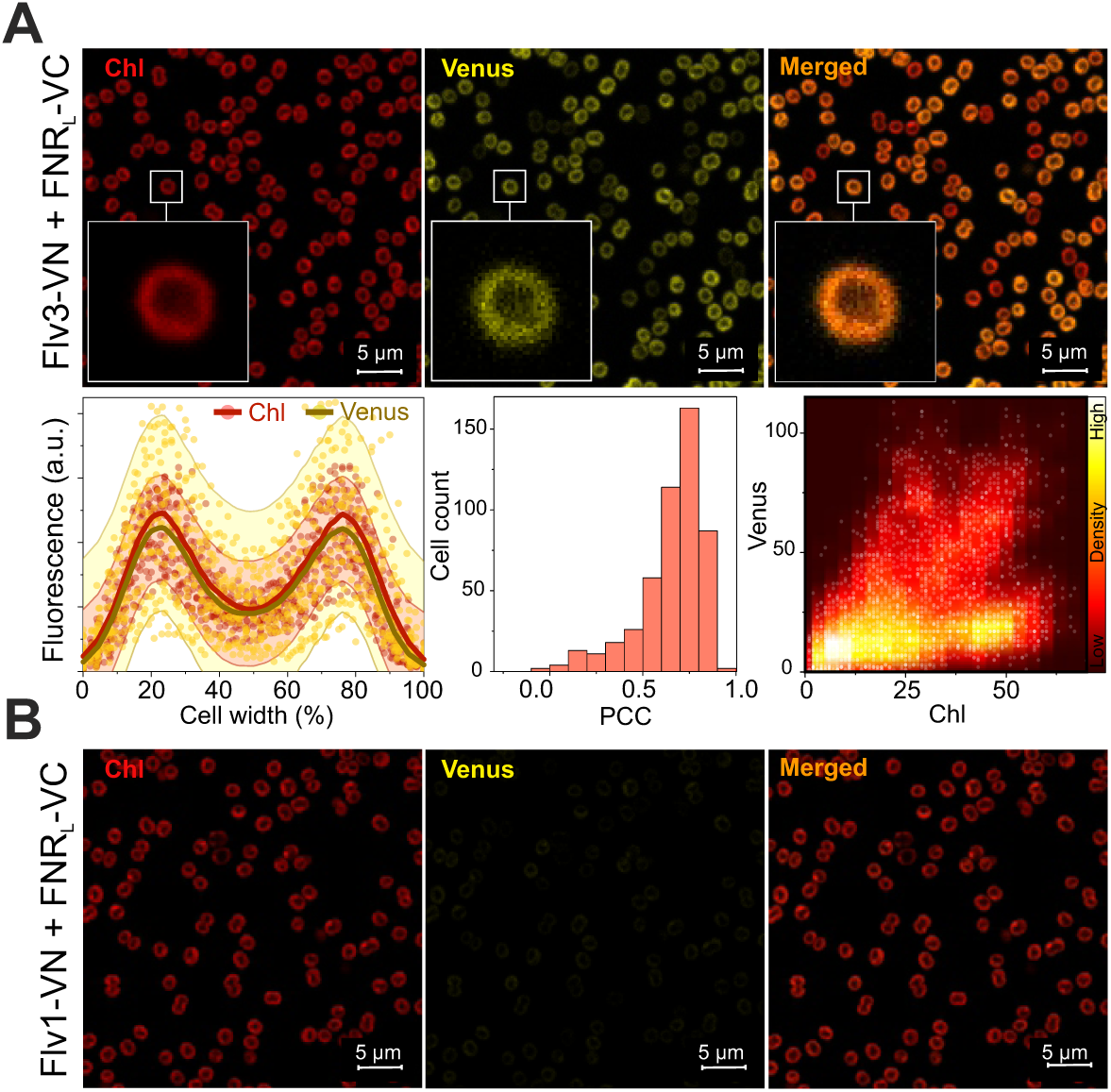
Protein-protein interaction tests between FNR_L_ and FDPs. **(A)** BiFC tests between Flv3 and FNR_L_ and **(B)** Flv1 and FNR_L_. Representative confocal micrographs from three independent experiments are presented, showing chlorophyll a (Chl) autofluorescence emanating from the thylakoid membranes in the left panel, fluorescence from re-assembled Venus I152L fluorescent proteins in the middle, and overlaid Chl and Venus fluorescence in the right panel. The inlets show zoomed-in views of a single cell. For quantification of the subcellular localisations of the interacting fusion protein pairs, the co-localisation of Chl (red) and Venus (yellow) fluorescence is also presented with fluorescence intensities over 5-pixel-wide cross-sections of 30 cells (with width normalised to 100%) shown as a scatterplot. Red and gold lines show moving regression curves produced by locally weighted scatterplot smoothing (LOWESS) with 100 points of window. 95% confidence intervals are shown as the pale colour shadows. The histogram shows the distribution of the Pearson Correlation Coefficient (PCC) for Chl and Venus fluorescence co-localisation in all cells from three micrographs from individual replicates. A PCC of 1 indicates perfect co-localisation and a PCC of 0 no co-localisation. The cytofluorogram shows co-localised Chl and Venus fluorescence intensities as a scatter plot.

No interaction was detected between Flv1-VN and FNR_L_-VC (Fig. 2B), or between Flv2-VC and FNR_L_-VN or FNR_S_-VN (Fig. S4). All FDP+FNR BiFC strains showed strong expression of both fusion proteins (Fig. S5).

### Deficiency of Fed2–11 does not impair the Mehler-like reaction

We proceeded to investigate how the lack of single or multiple Fed isoforms affects the Mehler-like reaction. To this end, we monitored light-induced reduction of O_2_ in *Synechocystis* mutant strains deficient in Fed3, Fed4, Fed6, Fed 7, 8, and 9, Fed9, Fed10, or Fed11. Fully segregated mutants of Fed1, Fed2 or Fed5 cannot be obtained (Wang et al. 2022), but partially segregated deletion strains of Fed2 and Fed5 were nonetheless examined. We quantified the activity of the Mehler-like reaction as a ratio between maximum O_2_ photoreduction and maximum gross O_2_ evolution. In none of the Fed2-11 mutants was major impairment of the Mehler-like reaction observed (Fig. 3, Fig. S6B). Slight but statistically non-significant decrease was observed in cells deficient in bacterial-type ferredoxins (ΔFed789 and (ΔFed9), while there was a non-significant increase in ΔFed4 (Fig. 3B).

**Fig. 3.**
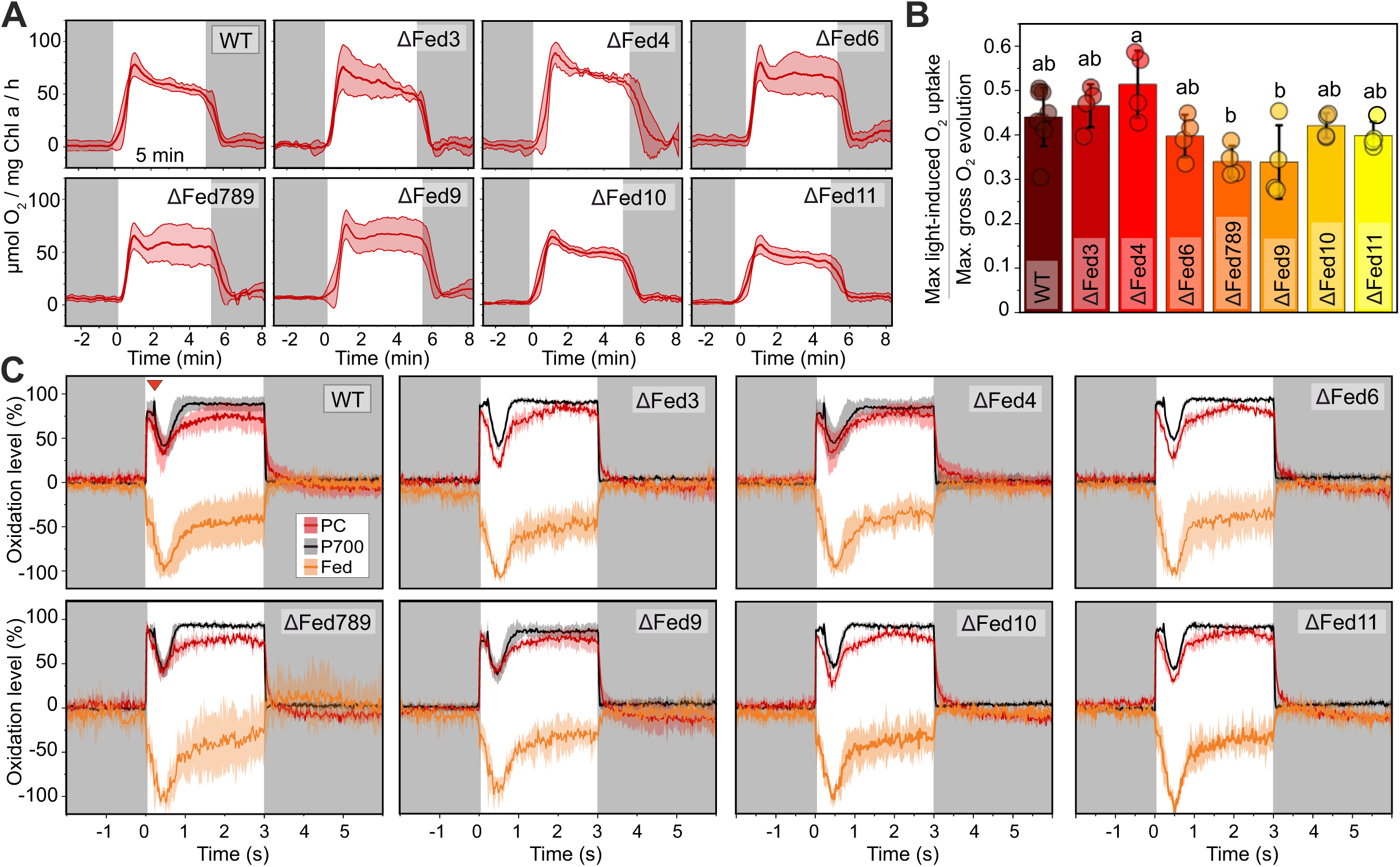
Effect of low abundance Ferredoxins on the Mehler-like reaction. **(A)** O_2_ photoreduction in cells deficient in low abundance Fed isoforms. O_2_ gas exchange was monitored by MIMS before, during, and after 5 min of illumination with strong actinic light (500 µmol photons m^-2^ s^-1^). Averaged traces from 8 (WT) or 4 (ΔFed3, ΔFed6, ΔFed789, ΔFed10, ΔFed11, ΔFed4, and ΔFed9) biological replicates with standard deviation are shown. **(B)** Averages with standard deviation of peak O_2_ photoreduction rate at c.a. 30 s of illumination normalised to the maximum gross rate of O_2_ evolution. Data are averages from 4 or 8 biological replicates as in (A) and individual data points are shown as circles. Statistical significance of differences is indicated by lower case letters according to one way ANOVA and Tukey’s post hoc test for comparison of means, with P<0.05 considered significant. **(C)** Redox changes of PC, P700, and Fed in WT and Fed mutant cells. PC, P700, and Fed redox changes as measured with a DUAL-KLAS-NIR spectrophotometer. The traces are normalised to the maximal oxidation values of PC and P700 and maximal reduction of Fed, as determined with the NIRMAX protocol, of which a 3 s illumination with actinic light is shown here (see Figure S5 for the full experiment), including a multiple turnover pulse after 200 ms (indicated by the red triangle in the WT panel). Averaged traces from 6 (WT), 5 (ΔFed4), 4 (ΔFed9), or 3 (ΔFed3, ΔFed6, ΔFed789, ΔFed10, and ΔFed11) biological replicates are shown, with standard deviation as the shadowed area.

We next used a Dual-KLAS-NIR spectrophotometer to monitor the redox changes of Fed, P700, and PC upon illumination in ΔFed mutants. All ΔFed mutants showed WT-like redox kinetics of Fed, PC, and P700 (Fig. 3C, Fig. S6C, Fig. S7). Low-abundance ferredoxins are thus unlikely to be needed as main electron donors for the Mehler-like reaction, or as final electron acceptors in the photosynthetic electron transport chain.

### Fed1 interacts with Flv1, Flv2, and Flv3, but not with Flv4

Fed1, the most abundant Fed isoform and the one mainly involved in photosynthetic electron transfer, is essential for survival of *Synechocystis* and a knockout mutant strain of it cannot be obtained (Poncelet et al. 1998; Cassier-Chauvat and Chauvat 2014; Gutekunst et al. 2014). We therefore proceeded to investigate potential interactions between Fed1 and FDPs using the BiFC system. When Fed1 fused to a C-terminal Venus fragment (Fed1-VC) was co-expressed in WT *Synechocystis* cells with Flv1 fused to an N-terminal Venus fragment (Flv1-VN), strong Venus fluorescence was detected, suggesting a protein-protein interaction between Fed1 and Flv1 (Fig. 4A). Fed1-Flv1 interactions co-localised with Chl autofluorescence, indicating that they occurred on the thylakoid membrane. In contrast, co-expression of Fed1-VN with Flv3-VC resulted in a Venus fluorescence signal that showed little co-localisation with Chl fluorescence, indicating a cytosolic localisation for Fed1-Flv3 interactions (Fig. 4B). A similar result was also obtained with an opposite orientation of the Venus fragment fusions (Flv3-VN+Fed1-VC, Fig. S8). Interestingly, a clear, largely cytosolic interaction was detected also between Fed1-VN and Flv2-VC (Fig. 4C), but Fed1-VC failed to interact with Flv4-VN (Fig. 4D), suggesting that Flv2/4 heterooligomers may also receive reducing power from Fed1, but that the interaction occurs strictly through Flv2.

**Fig. 4.**
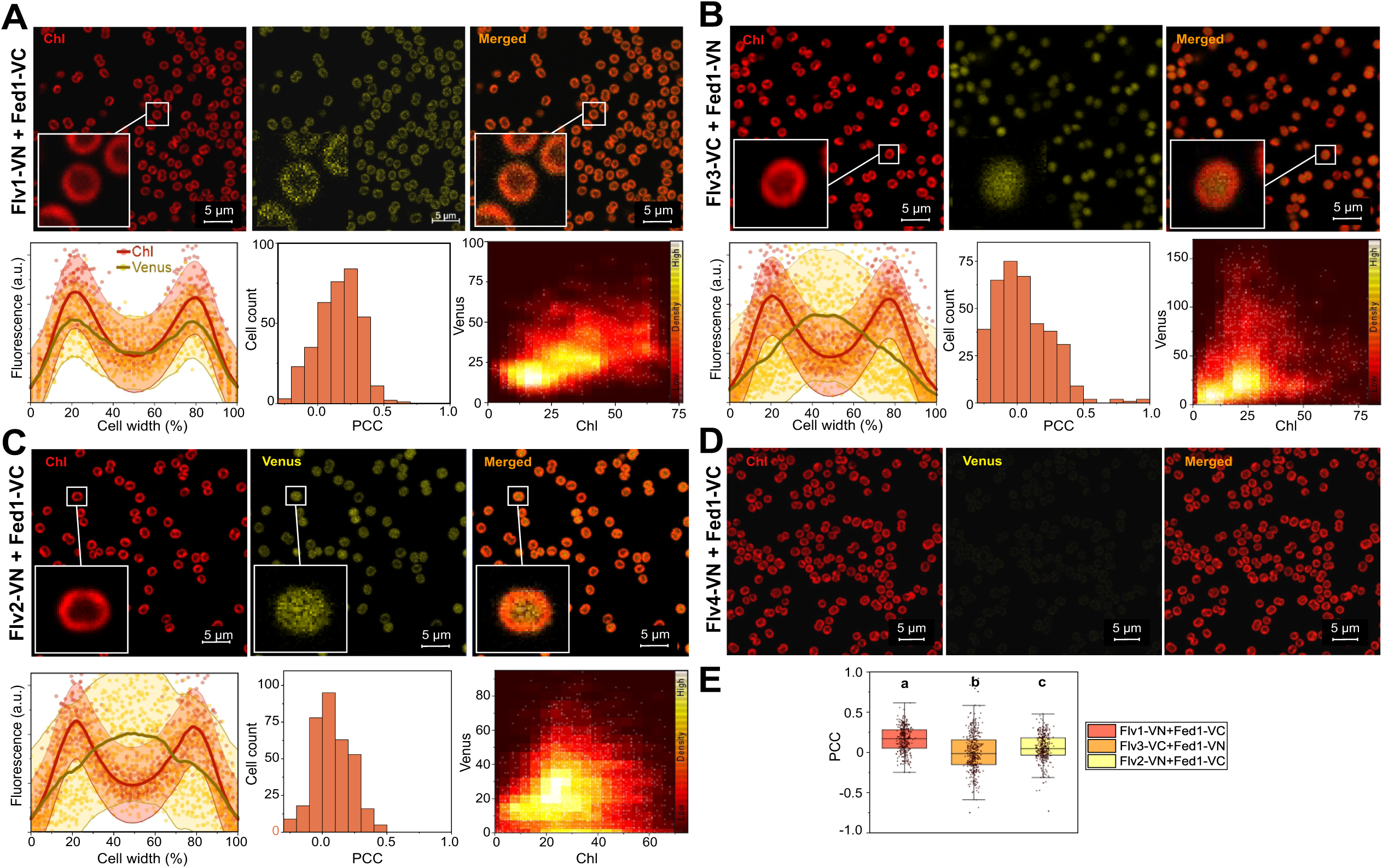
Interactions between Fed1 and FDPs in BiFC tests. Representative confocal micrographs from 3 independent experiments are presented, showing chlorophyll a (Chl) autofluorescence emanating from the thylakoid membranes in the left panel, fluorescence from re-assembled Venus I152L fluorescent proteins in the middle, and overlaid Chl and Venus fluorescence on the right panel. **(A)** Flv1-VN + Fed1-VC, **(B)** Flv3-VC + Fed1-VN, **(C)** Flv2-VN + Fed1-VC, **(D)** Flv4-VN + Fed1-VC. LOWESS smoothing was done with 120 points of window with 95% confidence intervals shown as the pale colour shadows. See the legend for Fig. 2 for other details on quantification of co-localisation. **(E)** Comparison of PCC values for Chl and Venus fluorescence co-localisation between Flv1-VN+Fed1-VC, Flv3-VC+Fed1-VN, and Flv2-VN+Fed1-VC cultures. Box plots with median ± SD and individual data points as dots are shown. Statistical significance is indicated by lower case letters according to one-way ANOVA and Tukey’s post-hoc tests for comparisons of means (P<0.05).

Motivated by the previously reported two-hybrid interaction between Fed9 and Flv3 (Cassier-Chauvat and Chauvat 2014) and the slightly but non-significantly decreased O_2_ photoreduction rate in the ΔFed9 mutant (Fig. 3B), we investigated potential *in vivo* interactions between Fed9 and Flv1 or Flv3. A thylakoid-localised interaction was indeed detected between Flv1-VN and Fed9-VC (Fig. S9A), and a mixed-localisation interaction between Flv3-VN and Fed9-VC (Fig. S9B). Therefore, we cannot exclude a minor or redundant role for Fed9 as an interactor with Flv1/Flv3 heterooligomers or Flv1 or Flv3 homooligomers.

### Flv1/3 and Flv2/4 interactions exhibit *pmf* dependent association with the thylakoid membrane

Interactions between the FDP heterooligomer pairs, Flv1-VN and Flv3-VC (Fig. 5A), and Flv4-VN and Flv2-VC (Fig. 5B) were localised predominantly in the cytosol. In smaller subpopulations of cells, however, the interactions were localised at the thylakoids (Fig. 5A–B, Fig. S10). This may indicate that the associations of Flv1/3 and Flv2/4 heterooligomers with the thylakoid membrane are dependent on the physiological state of the cell. We have previously hypothesised that such reversible membrane association, which would control the participation of FDP heterooligomers in O_2_ photoreduction, may be mediated by changes in cytosolic pH or proton motive force (*pmf*) which consists of transmembrane delta pH and electric potential component Δψ (Nikkanen et al. 2021b), or by changes in Mg^2+^ and Ca^2+^ cation concentrations (Zhang et al. 2012). To investigate these hypotheses, we determined the localisation of interactions between FDP isoforms when suspended in a medium containing either a low or high Mg^2+^ and Ca^2+^ concentration, as well as in the presence of the proton gradient uncoupler carbonyl cyanide m-chlorophenylhydrazone (CCCP). Measurement of the 500-480 nm electrochromic shift (ECS) signal (Viola et al. 2019) confirmed that 30 µM CCCP is sufficient to diminish the *pmf* by elevating the proton conductivity of the thylakoid membrane while only moderately affecting photosynthetic proton flux (Fig. S11). The initial generation of the *pmf* at dark-to-light transitions was decreased in low compared to high cation medium due to lower proton flux (Fig. S12).

**Fig. 5.**
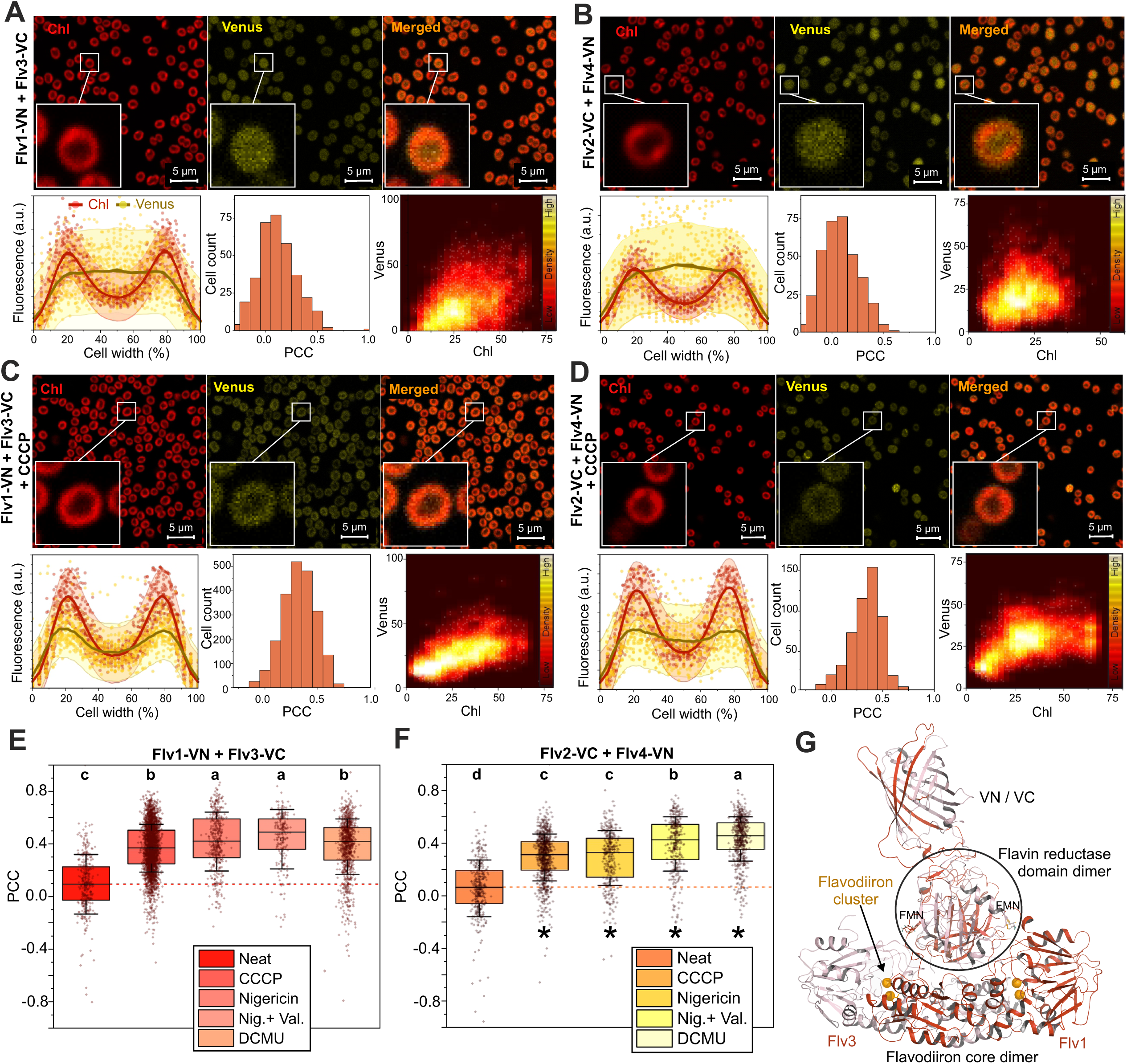
Subcellular localisations of Flv1/3 and Flv2/4 interactions in BiFC tests. **(A)** BiFC tests between Flv1-VN and Flv3-VC. **(B)** BiFC tests between Flv2-VC and Flv4-VN. **(C)** BiFC tests between Flv1-VN and Flv3-VC with 30 µM CCCP added in the medium. **(D)** BiFC tests between Flv2-VC and Flv4-VN with 30 µM CCCP added in the medium. Upper panels show representative confocal micrographs from the chlorophyll and Venus fluorescence channels as well as a merged image. See the legend for Fig. 1 for details on quantification of co-localisation. Fluorescence scatterplots of cell cross sections include data from 48, 50, 39, and 44 cells in A, B, C, and D, respectively. LOWESS smoothing was done with 120 points of window with 95% confidence intervals shown as the pale colour shadows. **(E-F)** Comparison of PCC values for Chl and Venus fluorescence co-localisation between un-treated (neat) Flv1-VN+Flv3-VC (E) and Flv2-VC+Flv4-VN (F) cultures and cultures treated with 30 µM CCCP, 20 mM nigericin, 20 mM nigericin (Nig.) and 40 µM valinomycin (Val.), or 20 µM DCMU. Box plots with median ± SD and individual data points as dots are shown. Statistical significance is indicated by lower case letters according to one-way ANOVA and Tukey’s post-hoc tests for comparisons of means (P<0.05). The dotted lines indicate the medians of the neat experiments. **(G)** Predicted structure of an Flv1-VN + Flv3-VC interaction complex.

Dissipation of the *pmf* due to addition of CCCP in the medium resulted in increased thylakoid localisation of both Flv1/3 (Fig. 5C) and Flv2/4 interactions (Fig. 5D). The average PCC of Flv1/3 and Flv2/4 interactions with Chl increased from 0.09±0.01(SEM) to 0.37±0.01 and from 0.06±0.01 to 0.33±0.01, respectively, upon addition of CCCP in the medium. Similar results were obtained by supplementing cultures with another uncoupler, nigericin, which, by acting as a H^+^/K^+^ exchanger, specifically dissipates the proton gradient and delays alkalization of the cytosol, as confirmed by an acridine orange fluorescence measurement (Fig. S13). In the presence of 20 µM nigericin the average PCC of Flv1/3 interactions with Chl increased to 0.41±0.01 (Fig. 5E, Fig. S14A) and that of Flv2/4 interactions to 0.28±0.01 (Fig. 5F, Fig. S14B). Interestingly, by supplementing nigericin with 40 µM of potassium-specific ionophore valinomycin further increased the PCC between Flv2/4 interactions and Chl fluorescence to 0.39±0.01 (Fig. 5F, Fig. S14D). Addition of DCMU to block electron transport from PSII, and thus partly *pmf* generation (Fig. S15), also significantly increased the PCC of both Flv1/3 and Flv2/4 interaction signals with Chl fluorescence (Fig. 5E–F), although results were visually more ambiguous than with the uncouplers (Fig. S15). We also investigated the subcellular localization of FDPs by immunoblotting. Addition of CCCP significantly decreased the amount of Flv1 in the soluble protein fraction and non-significantly increased it in the membrane fraction (Fig. S16A). CCCP also non-significantly decreased the amount of Flv2 in the soluble fraction, while Flv4 was exclusively detected in the membrane fraction (Fig. S16A). Some contamination of the membrane fractions by soluble proteins, as indicated by the presence of the large Rubisco subunit (Fig. S16B), resulted in variation between replicates and decreased the significance of observed effects.

Computational modelling of Flv1-VN+Flv3-VC interaction complexes predicted that the Venus fragment fusions should not interfere with formation of functional FDP hetero-oligomers (Fig. 5G). These results suggest that a low *pmf* induces association of FDP heterooligomers with the thylakoid membrane, likely enabling rapid activation of the Mehler–like reaction e.g. during dark-to-light transitions, while an increase in *pmf* and concomitant alkalisation of the cytosol would cause them to disassociate from the membrane. Indeed, addition of 15 µM CCCP, which results in dissipation of ∼30% of the *pmf* (Fig. S11), increased the peak O_2_ photoreduction rate relative to maximum O_2_ evolution from 39% to 69%, and 30 µM CCCP to 65% (Fig. S17A). No O_2_ photoreduction was detected in the presence of 30µM CCCP in the ΔFlv3 mutant (Fig. S17B), indicating that the CCCP-induced relative increase in the O_2_ photoreduction rate is dependent on FDPs. However, the fact that CCCP also has a significant inhibitory effect of photosynthesis makes interpretation of the MIMS results complex. Therefore, we also measured the O_2_ photoreduction rate in the presence of 20 µM nigericin, which only has a minor effect on O_2_ evolution (Fig. S18B). A statistically significant increase in the maximal O_2_ photoreduction rate (normalized to maximal O_2_ gross evolution rate) was indeed detected upon addition of nigericin to WT cells (Fig. 6), while no nigericin-induced O_2_ photoreduction was detected in the ΔFlv3 mutant (Fig. S18C).

**Fig. 6.**
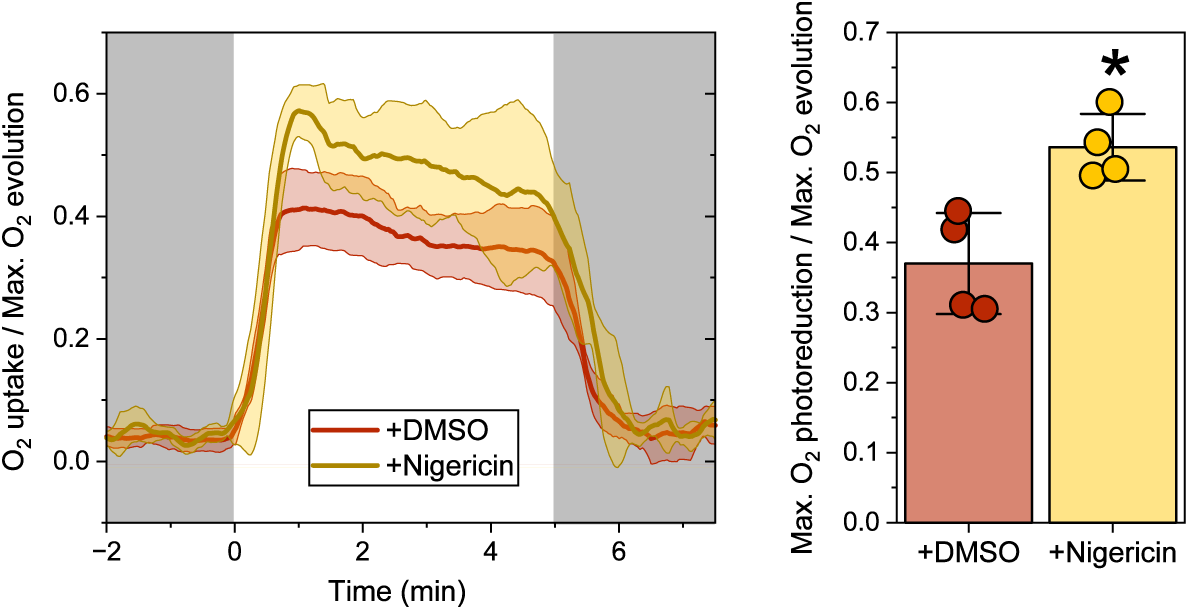
Effect of nigericin on the O_2_ photoreduction rate. O_2_ exchange in WT cells grown under 3% [CO_2_] at 50 µmol photons m^-2^ s^-1^ and at pH 7.5 was measured by MIMS during dark and 5 min of illumination at 500 µmol photons m^-2^ s^-1^. Cells were measured either in the presence of 4 µl DMSO solvent as control or with 20 µM nigericin (in 4 µl DMSO). The traces are shown as O_2_ uptake rate / maximal O_2_ gross evolution rate, and are averaged from four biological replicates ± SD. The right-hand panel shows the mean ratios between maximal light-induced O_2_ uptake and maximal gross O_2_ evolution with and without 20 µM nigericin ± SD with individual datapoints shown as circles. Statistical significance of the difference was tested by a two-sample t-test and is indicated by * (p=0.0113).

A high O_2_ photoreduction rate (up to 80-90% of gross O_2_ evolution) is also maintained in the mutant strain lacking the NdhD1 and NdhD2 subunits of the NDH-1 complex (ΔD1D2) (Nikkanen et al. 2020). Acridine yellow and -orange fluorescence measurements showed that alkalisation of the cytosol is delayed in the ΔD1D2 strain (Fig. S19), suggesting that maintaining a more acidic cytosolic pH for longer allows FDP heterooligomers to remain at the thylakoid membrane and to sustain a high rate of O_2_ photoreduction. Inhibition of respiratory terminal oxidases by potassium cyanide (KCN) also delayed cytosolic alkalisation (Fig. S19) and increased light-dependent O_2_ uptake (Fig. S20A). This increase likely derived from photorespiration rather than the Mehler-like reaction, as it was also detected in the ΔFlv3 D1D2 mutant and was coupled to CO_2_ production (Fig. S20B). At steady state, the cytosolic pH in KCN-treated cells was slightly higher (Fig. S19A), and *pmf* significantly higher than in untreated cells (Fig. S20C). In line with the higher *pmf*, addition of KCN decreased the association of Flv2/4 interactions with the thylakoid membrane (Fig. S21C–D), while no significant effect was observed for Flv1/3 interactions (Fig. S22C–D).

We also tested the effect of altered cellular thiol redox state on the localisation of Flv1/3 or Flv2/4 interactions by supplementing the cultures with 2mM DTT or 100 µM CuCl_2_ to create reducing or oxidizing conditions, respectively. DTT had no effect on the subcellular localization of the FDP hetero-oligomer interactions, while CuCl_2_ very slightly decreased the thylakoid association of Flv1/Flv3 and Flv2/Flv4 interactions (Fig. S22 and Fig. S21). We also confirmed that 2 mM DTT did not significantly affect *pmf* generation, O_2_ evolution, or light-induced O_2_ uptake. DTT did, however, increase the level of dark-respiration, while addition of CuCl_2_ resulted in partial inhibition of O_2_ evolution and slightly elevated *pmf* (Fig. S20). No difference in the localisation of Flv1/3 or Flv2/4 interactions was detected between low or high cation concentration in the medium (Fig. S23).

### Subcellular localization of Flv1 an Flv3 self-interactions

Flv1 interacted with itself, and similarly to Flv1-Fed1 interactions, primarily on the thylakoids (Fig. S24A), suggesting either homooligomerisation or involvement in higher-order heterooligomers. Flv3 self-interactions showed more variation in their subcellular localisation, with two distinct populations of cells. Most cells exhibited relatively weak interaction signals localising on thylakoids while stronger signals localising in the cytosol were detected from a subpopulation of cells (Fig. S24B). Both a high cation concentration in the medium and addition of CCCP slightly decreased the amount of cytosolic Flv3 self-interactions (Fig. S25).

No interaction was detected between Flv3-VN and Flv2-VC (Fig. S24C) despite strong expression of both fusion proteins (Fig. S5), supporting the notion that Flv2 and Flv3 are unable to form heterooligomers *in vivo* (Mustila et al. 2016), as well as providing a negative control for the observed interactions.

### Surface charge modelling supports pH-dependent association of FDP heterooligomers with the thylakoid membrane

Lastly, we examined the molecular mechanism underlying the ΔpH-dependent association of FDP heterooligomers with the thylakoid membrane. At low *pmf*, low pH on the cytosolic side of the thylakoid membrane may cause protonation of acidic residues in proteins, giving them a more positive surface charge and promoting electrostatic interactions with negatively charged membrane lipids, or possibly with another protein or a protein complex in the membrane (Johnson and Cornell 1999; Mulgrew-Nesbitt et al. 2006). To test this hypothesis, we modelled the electrostatic surface potentials of FDP monomers and oligomers at different physiologically relevant pH values. In cyanobacteria, cytosolic pH ranges between 7 in dark or low light and 8.5 under strong illumination, also depending on extracellular pH (Coleman and Colman 1981; Mangan et al. 2016).

Our analysis revealed that at pH 7.0, Flv1 and Flv3 core monomers have net surface charges of −3.0 ± 0.2e and −11.5 ± 0.6e, respectively. In contrast, at a more alkaline pH of 8.5, the net surface charges decrease to −9.3 ± 0.6e for Flv1 and −18.5 ± 1.1e for Flv3 (Fig. S26A, C). Similarly, Flv1 core homodimers have a small negative net charge of −3.1 ± 0.8e, with positively charged patches at neutral pH. Those positive patches remain even at alkaline pH, when the Flv1 core homodimer has a negative net charge of −18.9 ± 1.2e, allowing interactions with the membrane, albeit with lower affinity (Fig. 7A). The Flv3 core homodimer carries a negative net charge of −12.4 ± 1.9e at pH 7.0, which decreases as low as −29.8 ± 0.8e at pH 8.5 (Fig. 7B). Flv1/3 core heterooligomers have a net charge of −11.4 ± 1.6e, with positively charged patches that likely permit interaction with negatively charged membrane lipids. At pH 8.5, the Flv1/3 heterooligomer becomes highly negatively charged (−25.9 ± 1.5e), making membrane interactions unfavourable due to electrostatic repulsion (Fig. 7C).

**Fig. 7.**
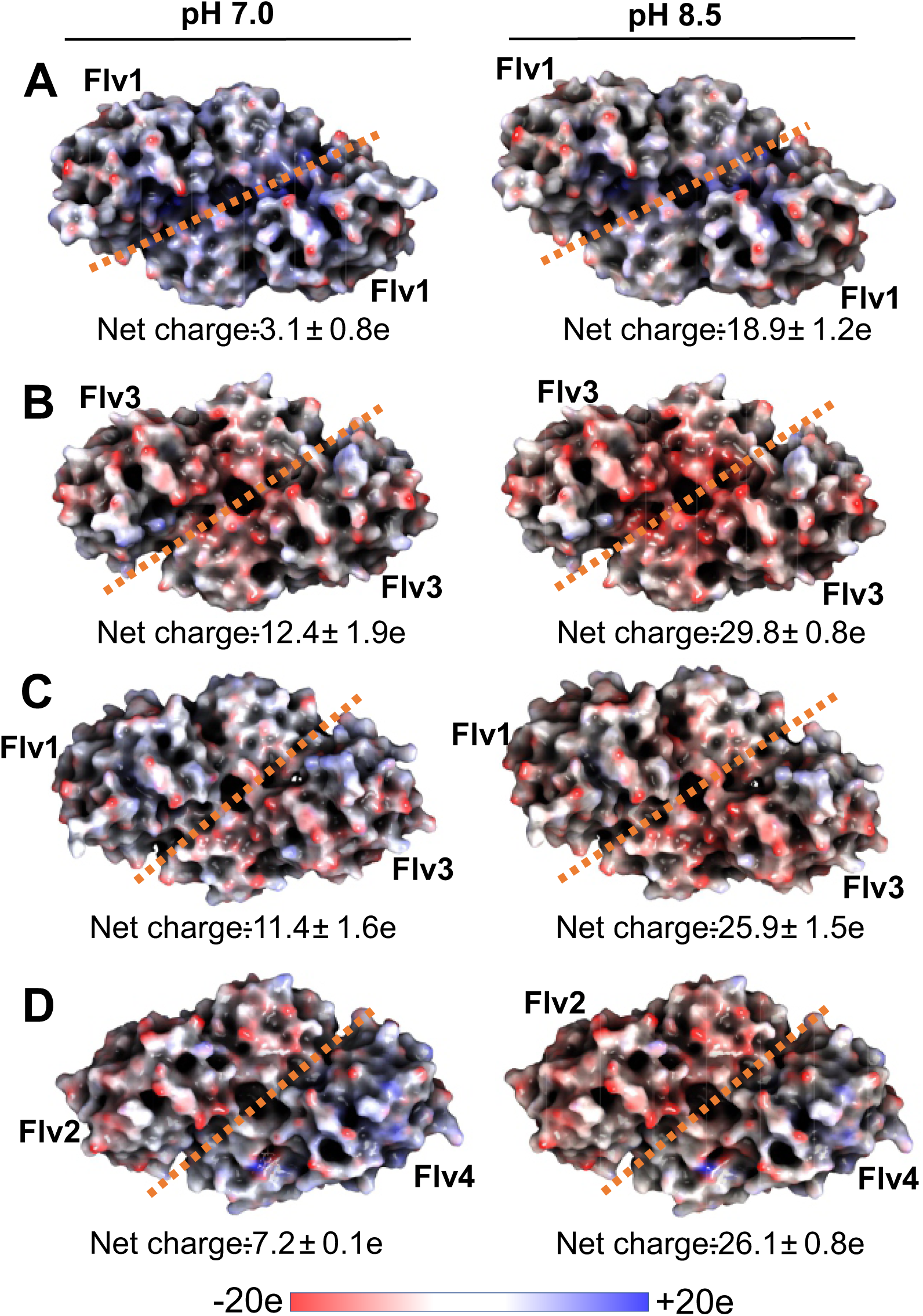
*In silico* analysis of the electrostatic surface of FDP oligomers at different pH. **(A)** Flv1/Flv1 homodimer, **(B)** Flv3/Flv3 homodimer, **(C)** Flv1/Flv3 heterodimer, and **(D)** Flv2/Flv4 heterodimer. Surface charges are color-coded with red representing negative charge, blue positive charge, and white neutral regions. The left and right panels show charges at pH 7.0 and pH 8.5, respectively. Since the relative orientation of the C-terminal Flavin reductase domain could not be modelled reliably, only the β- lactamase-like and Flavodoxin-like domains are shown. The orange line indicates the symmetry axis between monomers.

Monomeric Flv2 and Flv4, in turn, carry contrasting net charges. While Flv2 has a highly negative net charge at both pH 7.0 and 8.5 (−16.2 ± 0.4e and −24.2 ± 0.8e), Flv4 monomers have a positive surface charge (+5.8 ± 0.1e) at pH 7.0 and a slightly negative charge (−2.2 ± 0.2e) with large positive patches at pH 8.5 (Fig. S26B, D). The Flv2/Flv4 heterooligomer has a mostly negative charge of −7.2 ± 0.1e at pH 7.0. At pH 8.5, interaction with a negatively charged membrane surface becomes compromised due to a highly negative net charge of −26.1 ± 0.8e (Fig. 7D).

## Discussion

Light-dependent reduction of O_2_ to H_2_O by FDPs, known as the Mehler-like reaction, constitutes a vital, strictly regulated outlet for excessive electrons in the PETC during fluctuations in environmental conditions (Allahverdiyeva et al. 2013; Alboresi et al. 2019; Nikkanen et al. 2021b). Photosynthetic microorganisms are increasingly employed as green cell factories for sustainable bioproduction of desired compounds, and a thorough understanding of the regulatory mechanisms of photosynthesis is crucial for optimising the light-energy-to-product conversion efficiency in those platforms (Nikkanen et al. 2021a; Hubáček et al. 2024). In recent years, it has become evident that FDPs can take several homo- and heterooligomeric conformations, but heterooligomers consisting of Flv1 and Flv3 as well as Flv2 and Flv4 are required for the Mehler-like reaction under air-level [CO_2_] in *Synechocystis* (Mustila et al. 2016; Santana-Sanchez et al. 2019). Several key aspects of this essential regulatory mechanism of photosynthesis, however, have remained unclear. A) What is the electron donor to FDPs, and does the donor differ between different homo- and heterooligomeric conformations? B) What kind of oligomers can FDPs form and how do they localise within the cell? C) How is the activity of the Mehler-like reaction controlled? An experimental approach combining a BiFC system to visualise protein–protein interactions, biophysical characterisation of photosynthetic electron transport in *Synechocystis* mutant strains, and *in silico* surface charge modelling allowed us to address these questions in the present study. First, we identified *in vivo* interactions between Fed1 and Flv1, Flv2, and Flv3, and confirmed Fed1 as the primary electron donor to both Flv1/3 and Flv2/4 heterooligomers. We also showed that the subcellular localisation differs between different FDP/FDP and FDP/Fed1 interactions, and finally, that the association of Flv1/3 and Flv2/4 interactions with the thylakoid membrane depends on the trans-thylakoid proton gradient (*pmf*), suggesting a feedback mechanism to dynamically control the activity of the Mehler-like reaction.

Previous studies have shown NAD(P)H to be able to donate electrons to recombinant Flv1 or Flv3 (Vicente et al. 2002; Brown et al. 2019) as well as to Flv4 (Shimakawa et al. 2015) *in vitro*. However, the measured rates are far too low to account for the observed *in vivo* rates of O_2_ photoreduction. The maximal *in vitro* rates of NAD(P)H-dependent O_2_ reduction by Flv1 or Flv3 as measured by (Brown et al. 2019) were roughly 30 µmol O_2_ / mg Flv1 / min. Maximal *in vivo* O_2_ photoreduction rates under strong illumination in WT *Synechocystis* grown in air-level [CO_2_], on the other hand, are around 100 µmol O_2_ / mg Chl / hr (Santana-Sanchez et al. 2019). In high [CO_2_] conditions, a similar rate is maintained also at steady state even though Flv2 and Flv4 are not expressed (Santana-Sanchez et al. 2019) and Flv1 and Flv3 are less abundant than in air-level [CO_2_] (Zhang et al. 2009). Based on recent absolute quantification of Flv1 proteins (1100–1400 copies per cell) and chlorophyll molecules per cell (∼1.54E+07 copies per cell) (Jackson et al. 2023), we can convert that *in vivo* rate to ∼270 µmol O_2_ / mg Flv1 / min. This is nine-fold higher than the NAD(P)H-dependent *in vitro* rates reported by Brown et al. for Flv1 (Brown et al. 2019). The *in vitro* rate for NADH-dependent O_2_ reduction by Flv4 was several orders of magnitude lower still, at ∼0,29 µmol O_2_ / mg Flv4 / min (Shimakawa et al. 2015).

Moreover, Flv1 and Flv3 monomers or homooligomers cannot catalyse the Mehler-like reaction *in vivo*, (Mustila et al. 2016), and in this study we demonstrated that severe impairment of light-induced NADP^+^ reduction in the ΔFNR_L_ mutant does not significantly decrease the O_2_ photoreduction activity of FDPs or the ability to rapidly re-oxidise Fed or P700 under strong illumination (Fig. 1). The slight decrease in O_2_ photoreduction rate observed in ΔFNR_L_ under air-level [CO_2_] was due to diminished Flv2 content, because under elevated [CO_2_], where the *flv4-2* operon is not expressed (Zhang et al. 2009), the difference in O_2_ photoreduction rate between WT and ΔFNR_L_ was abolished (Fig. 1). We hypothesise that the downregulation of Flv2 in the ΔFNR_L_ mutant in air-level [CO_2_] could be due to a disturbed redox state of the NADP^+^ / NADPH pool in the absence of FNR_L_ causing physiological conditions that result in the same downregulatory signal for expression of the *flv4-2* operon that happens in high [CO_2_] or highly alkaline (Santana-Sanchez et al. 2019) conditions. The slight increase in the abundances of Flv1 and Flv3 in ΔFNR_S_ and (insignificantly) in ΔFNR_L_ (Fig. 1E), in turn, could arise from the lack of FNR as an electron acceptor from Fed, which would cause more reduction pressure on the Fed pool, possibly triggering an up-regulatory signal to increase the transcription of *flv1* and *flv3* to relieve it. Moreover, as FNR_S_ likely feeds electrons to NDH-1 catalysing CET (Miller et al. 2022), and functional redundancy exists between Flv1/Flv3 and NDH-1 (Nikkanen et al. 2020; Storti et al. 2020), the upregulation of Flv1 and Flv3 may constitute a compensation for impaired CET in ΔFNR_S_.

Although NADPH is also produced in the dark in the oxidative pentose phosphate pathway, and FDPs have also been shown to be able to use NADH to catalyse O_2_ reduction *in vitro* (Shimakawa et al. 2015; Brown et al. 2019), it is highly unlikely that the light-independent NADPH or NADH production would be responsible for powering the strongly light-dependent Mehler-like reaction in the ΔFNR_L_ mutant. We detected increased dark-respiration in ΔFNR_L_ (Fig. S1C), possibly due to an elevated amount of the FNR_S_ isoform that is suggested to feed electrons from NADPH to NDH-1 via Fed in a respiratory/cyclic pathway (Miller et al. 2022). This would rather cause the NADP^+^/NADPH pool to be more oxidized in the dark than in WT, making it even less likely that FDPs would have sufficient electron donors if NADPH was in fact the main donor. Together with previous reports on Fed and NADPH redox kinetics in FDP mutants (Nikkanen et al. 2020; Sétif et al. 2020) these results indicate that neither NADPH nor FNR can be the main electron donor to the Mehler-like reaction catalysed by Flv1/3 and Flv2/4 heterooligomers. This poses the question about the physiological role of the C-terminal domain with putative NAD(P)H:flavin oxidoreductase activity. It has been suggested that the domain could participate in Fed binding, or be involved in taking electrons from a reduced flavodoxin domain and using them to reduce NADP^+^ (Sétif et al. 2020). This could protect the iron-sulfur center of the enzyme by relieving its over-reduction, for example in low [O_2_] conditions (Sétif et al. 2020). We suggest that the function of the C-terminal domain may also be related to the unknown physiological function of FDP homo-oligomers.

We observed a protein–protein interaction between FNR_L_ and Flv3 that strictly localised to the thylakoid membrane (Fig. 2). The MIMS results suggest that the interaction does not occur due to FNR_L_ functioning as an electron donor to Flv1/3 heterooligomers in the Mehler-like reaction. Instead, considering no interaction was observed between FNR_L_ and Flv1, FNR_L_ may interact with Flv3 homooligomers to perform an unknown function on thylakoids, or possibly associated with the phycobilisomes, to which FNR binds in cyanobacteria (Liu et al. 2019). While Flv3 homooligomers do not catalyse the Mehler-like reaction, Flv3 is 4 to 6-fold more abundant than Flv1 in cells (Jackson et al. 2023), and overexpression of Flv3 in ΔFlv1 mutant background does partially rescue the growth phenotype of ΔFlv1 in fluctuating light (Mustila et al. 2016), suggesting some non-Mehler-like reaction-related photoprotective function for Flv3 homooligomers. Considering the *in vitro* electron transfer from NAD(P)H to recombinant Flv3 and the presence of the NAD(P)H: flavin oxidoreductase-like domain in cyanobacterial FDPs, it is possible that NAD(P)H could donate electrons to Flv3 homooligomers, possibly facilitated by the homooligomers interacting directly with FNR_L_. We cannot fully exclude the possibility of a false positive BiFC result between Flv3 and FNR_L_ due to the proximity of the overexpressed proteins at the thylakoid membrane. However, the strong Venus fluorescence signal and the near absence of a signal between Flv1 and FNR_L_, despite similar proximity and comparable abundance of fusion proteins (Fig. S5), suggest that this is unlikely. Negative results from BiFC, should nonetheless, also be interpreted with caution and cannot be taken as definitive proof of lack of *in vivo* interaction. Unfavourable structural orientation of the Venus fragments in the interaction complex, steric hindrance, or improper folding of the fusion proteins may prevent reassembly of a functional fluorophore (Kudla and Bock 2016). In the current study, to minimize the topological constrains and steric hindrance, we introduced a flexible linker sequence between the proteins of interest and the Venus fragments. Computational modelling (Fig. 5G) supported the reliability of the BiFC tests, predicting that FDP-VN/C monomers from two different oligomers would be unlikely to interact with each other and thus contribute to the total fluorescence signal, since the interacting motifs of the FDPs are blocked within the interaction complex. Any interactions with endogenous FDPs will also be negligible, due to the vastly higher abundance of the expressed fusion proteins. This is especially true of Flv2 and Flv4, which are not expressed under high [CO_2_], as in the current study.

Recent *in vivo* studies have suggested Fed as a likely electron donor to Flv1/3 (Nikkanen et al. 2020; Sétif et al. 2020), and a recombinant class A FDP from the protozoan *Entamoeba histolytica* has been shown to be able to oxidise Fed *in vitro* (Cabeza et al. 2015). Out of the low-abundance Fed isoforms (Fed2-Fed11), Fed2 has been shown to be essential for the photoautotrophic growth of *Synechocystis* (Cassier-Chauvat and Chauvat 2014). However, Fed2 functions in low-iron response instead of photosynthetic electron transfer (Schorsch et al. 2018), and is unlikely to be a primary electron donor to FDPs. Fed7, 8, and 9, in turn, have been proposed to function in a stress response pathway in cooperation with the thioredoxin system (Marteyn et al. 2009; Mustila et al. 2014), and Fed9 was more recently shown to be important for photomixotrophic growth of *Synechocystis*, likely receiving electrons from pyruvate:ferredoxin oxidoreductase (PFOR) (Wang et al. 2022). Bacterial-type low-abundance ferredoxins are thus unlikely to constitute the main source of electrons for the light-induced Mehler-like reaction, despite the two-hybrid interaction reported between Fed9 and Flv3 (Cassier-Chauvat and Chauvat 2014). Accordingly, we detected no statistically significant impairment of the Mehler-like reaction in any of the mutant strains lacking Fed2-Fed11 (Fig. 3, Fig. S6, Fig. S7). We showed here that Fed1 interacts *in vivo* with Flv1, Flv3, as well as Flv2 (Fig. 4). Although we cannot exclude minor or redundant roles for low-abundance Feds such as Fed9, and direct *in vitro* evidence of Fed-dependent O_2_ reduction by cyanobacterial FDP hetero-oligomers is still lacking, these results pinpoint Fed1 as the main electron donor to FDP heterooligomers.

Flv1/Fed1 and Flv3/Fed1 interactions showed contrasting subcellular localisations, as Flv1 interacted with Fed1 at the thylakoids and Flv3 mostly in the cytosol (Fig. 4). Due to strong expression of the fusion proteins (Fig. S5), the size of the Flv3 pool in Fed1-VN+Flv3-VC cells substantially exceeds that of Flv1, likely causing a large proportion of Flv3-VC proteins to form homooligomers. The opposite is true in Fed1-VC+Flv1-VN cells. This would suggest that Flv1 homooligomers localise to the thylakoids and Flv3 homooligomers to the cytosol. However, in a majority of cells Flv3 self-interactions (Fig. S24B) as well as Flv3-FNR_L_ interactions (Fig. 2A) localised to the thylakoids, and the thylakoidal Flv3 self-interaction signals intensified in high concentrations of Ca^2+^ and Mg^2+^ cations (Fig. S25). These results suggest that formation of Flv3 homooligomers and/or their association with the thylakoids and FNR may be controlled by the physiological state of the cell, while Flv1 homooligomers consistently associate with the thylakoid membrane. These observations were supported by modelling of surface charges, showing that Flv1 homooligomers carry a low net negative charge with large positively charged patches that likely allow interaction with the thylakoid membrane, while the net surface charge of Flv3 homooligomers was highly negative, making their affinity to negatively charged surface of the thylakoid membrane (Barber 1982) low (Fig. 7). It is clear, however, that further studies are required to elucidate the physiological roles of FDP homooligomers. There also appears to be variation between different cyanobacterial species regarding formation of FDP oligomers and their physiological significance. Our recent study with the filamentous cyanobacterial model species *Anabaena* sp. PCC 7120 revealed that while vegetative cell specific Flv1A and Flv3A likely form heterooligomers and function similarly to their *Synechocystis* orthologues, Flv3A on its own is able to contribute to the Mehler-like reaction and photoprotection of PSI via an unknown mechanism that requires the presence of Flv2 and Flv4 (Santana-Sánchez et al. 2023). This suggests the existence of functional oligomers consisting of Flv3 and Flv2/4 or possibly interaction of Flv3 homooligomers with Flv2/4 heterooligomers in vegetative *Anabaena* cells. In line with a previous biochemical analysis, we detected no interaction between Flv3 and Flv2 in *Synechocystis* (Fig. S24C), but it remains to be confirmed whether such interactions or heterooligomerisation occur in *Anabaena* sp. PCC 7120 or other cyanobacterial species.

Importantly, the addition of the *pmf*-uncoupling ionophore CCCP or the K^+^/H^+^ exchanging uncoupler nigericin to the medium resulted in increased thylakoid localisation of Flv1/3 and, to lesser extent, Flv2/4 interactions (Fig. 5, Fig. S14), and increased the maximal light-induced O_2_ uptake rate relative to O_2_ evolution (Fig. 6, Fig. S18). Under air-level [CO_2_], the strong O_2_ photoreduction catalysed by Flv1/3 heterooligomers occurs transiently at the dark-light transition or upon sudden increases in light intensity, providing an outlet for excessive electrons in the PETC (Santana-Sanchez et al. 2019). The transience of this reaction suggests reversible post-translational regulation of Flv1/3 activity, and several putative mechanisms, such as a redox regulation by the thioredoxin system or phosphorylation and de-phosphorylation have been proposed (Alboresi et al. 2019; Nikkanen et al. 2021b; Beraldo et al. 2024). Our findings in the current study suggest that FDP heterooligomers are amphitropic: the magnitude of the *pmf* or cytosolic pH regulates their activity by determining whether they associate with the thylakoid membrane. As we have previously shown that up to 75% of *pmf* generation during dark-to-light transitions is dependent on FDP-catalysed O_2_ photoreduction in *Synechocystis* (Nikkanen et al. 2020), we propose a self-regulatory feedback loop. In darkness or at low light *pmf* is low and cytosolic pH is close to neutral. This condition allows FDP heterooligomers to associate with the thylakoid membrane catalysing a rapid and strong reduction of O_2_ upon illumination (Fig. 8A). This occurs when CO_2_ fixation has not been yet fully activated, and, due to *pmf* being low, photosynthetic control at Cytochrome b_6_f is not limiting electron transport to PSI. As a result, the presence of a strong FDP electron sink becomes central for photoprotection. The high rate of O_2_ photoreduction elevates the *pmf* directly by consuming protons on the cytosolic side of the membrane and indirectly by keeping the Fed1 pool oxidised, thereby allowing a higher electron transport rate in the PETC. The higher *pmf* then causes Flv1/3 heterooligomers to disassociate from the thylakoids, decreasing their O_2_ reduction activity (Fig. 8B). In this manner, high Flv1/3 activity during the first minute of illumination would result in downregulation of Flv1/3 activity after the initial peak. This is exactly what we have previously reported: O_2_ photoreduction peaks after c.a. 20-30 seconds of strong illumination, which is also when a steady-state level of *pmf* is reached in WT cells (Nikkanen et al. 2020). After 30 seconds, the rate of O_2_ photoreduction declines, especially in low [CO_2_] grown cells or under alkaline pH (when *Flv2* and *Flv4* are not expressed) (Santana-Sanchez et al. 2019).

**Fig. 8.**
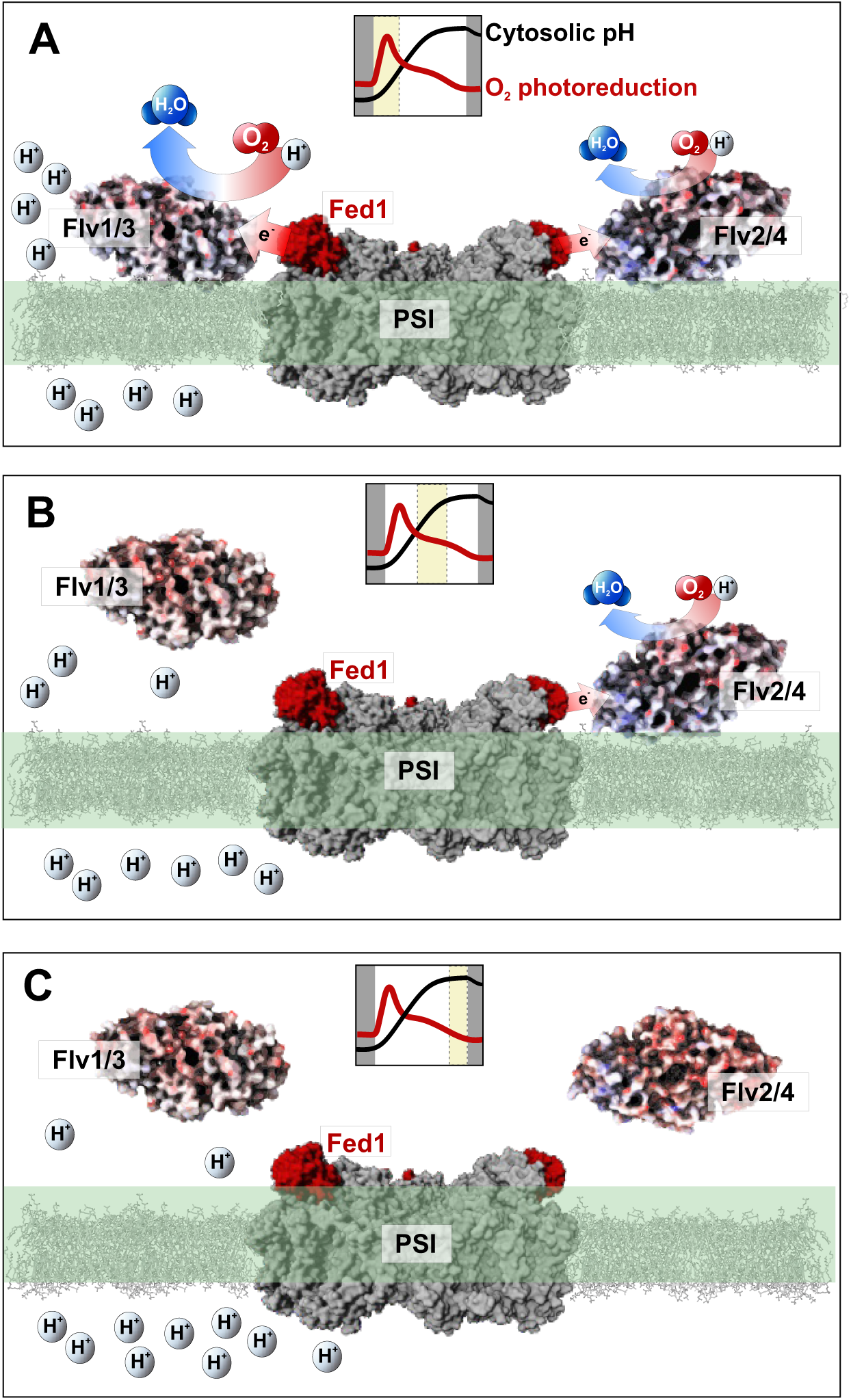
Schematic model for pH-dependent association of FDP heterooligomers with the thylakoid membrane. **(A)** In dark or low light when *pmf* is low and cytosolic pH close to neutral, a more positive surface charge of FDP heterooligomers allows them to bind to the thylakoid membrane. This enables the rapid activation of FDPs upon illumination, catalysing a strong Mehler-like reaction using electrons from reduced Fed1. The process triggers swift *pmf* generation during dark-to-light or low-to high light transitions in *Synechocystis* (Nikkanen et al. 2020). **(B)** As *pmf* rises and the cytosol becomes increasingly alkaline, a self-regulatory feedback mechanism of FDPs is initiated. Flv1/3 heterooligomers are repelled from the membrane due to their surface charges becoming more negative. This inhibits Flv1/3 activity and results in the decline of the O_2_ photoreduction rate. Due to the positive charge of Flv4, however, Flv2/4 heterooligomers still maintain affinity to the thylakoid membrane, catalysing residual O_2_ photoreduction. **(C)** When the cytosol becomes highly alkaline upon full activation of CO_2_ fixation, both FDP heterooligomers disassociate from the thylakoids, strongly inhibiting (now unnecessary) O_2_ photoreduction activity. Trimeric PSI structure with bound Fed is based on (Li et al. 2022). The small inlets within each panel represent typical kinetics of O_2_ photoreduction in red, and alkalisation of the cytosol (see Fig. S13B) in black during a dark-to-light transition. The yellow rectangles indicate the phase depicted in the panel.

It has been reported previously that Flv2 and Flv4 are in the thylakoid fraction when *Synechocystis* proteins are extracted in a buffer containing high concentrations of Mg^2+^ and Ca^2+^ cations, and in the soluble fraction when extracted in a cation-depleted buffer (Zhang et al. 2012). *Pmf* generation was delayed during dark-to-light transitions in low cation in comparison to high cation medium (Fig. S12), but likely due to impairment of the function of unknown cation channels or antiporters on the thylakoid membrane (Carraretto et al. 2016), resulting in diminishment of initial Δψ component of *pmf*. However, we detected no difference in the subcellular localisation of Flv2/4 interactions between low and high cation media (Fig. S23C–D). Moreover, increased thylakoid localization of FDP heterooligomer interactions was induced by the addition of nigericin, which acts as H^+^/K^+^ exchanger, specifically dissipating the proton gradient while maintaining membrane potential (Fig. 5, Fig. S14). A slight further increase in Flv2/Flv4 but not in Flv1/Flv3 thylakoid localisation was induced by the addition of the potassium-specific ionophore valinomycin (Fig. 5, Fig. S14). These observations suggest that the loss of the ΔpH component of *pmf,* specifically, promotes the thylakoid association of at least Flv1/Flv3 heterooligomers, while the localisation of Flv2/Flv4 heterooligomers may additionally be affected by the ionic environment across the thylakoid membrane.

In plant chloroplasts low stromal pH promotes the binding of FNR to its membrane anchor Tic62 (Benz et al. 2009). It is necessary to investigate in future studies whether the FDP:Fed1 interactions observed herein are themselves pH-dependent, and to elucidate the molecular mechanism of electron transfer from Fed1 to FDP heterooligomers.

*In silico* modelling revealed that the molecular mechanism underlying the reversible association of FDP heterooligomers with the thylakoid membrane depends on changes in net surface charges within a physiological pH range (Fig. 7). In neutral cytosolic pH such as in dark or low light conditions, Flv1/3 and Flv2/4 heterooligomers maintain low negative net charge with positive patches that enable interaction with membrane lipids or positively charged membrane proteins. In contrast, at higher pH mimicking the alkalisation of the cytosol upon strong illumination, the net charges of FDP heterooligomers become highly negative, repelling them from the membrane. As the cytosol gradually gets more alkaline under illumination (Fig. S13), the positive net charge of Flv4 (Fig. S26D) may allow Flv2/4 heterooligomers to remain bound to the membrane longer than Flv1/3 heterooligomers (Fig. 8B- C). This could explain the distinct kinetics of Flv1/3 and Flv2/4-mediated O_2_ photoreduction, where Flv1/3 catalyses a strong but transient reaction, and Flv2/4 a more sustained O_2_ photoreduction further into dark-light transitions (Santana-Sanchez et al. 2019).

The proposed pH-dependent regulatory mechanism of FDP activity may also indirectly control the induction of the carbon concentrating mechanism (CCM), as at least in algae, CCM induction was recently shown to depend on *pmf* generation by FDPs and CET (Burlacot et al. 2022). Our recent results with *Anabaena* sp. PCC 7120 suggest that a similar dependency may also exist in cyanobacteria (Santana-Sánchez et al. 2023).

In conclusion, the present study elucidates the regulation of the Mehler-like reaction and advances our understanding of how photoprotective mechanisms are orchestrated in dynamic, natural light environments. Our findings support Fed1 as the primary electron donor for the reaction catalysed by FDP heterooligomers. Furthermore, we demonstrated that the activity of the Mehler-like reaction is regulated by reversible association with the thylakoid membrane via pH-dependent changes in electrostatic surface charges of FDP heterooligomers. We propose a model of a self-regulatory feedback mechanism that controls the activity of the Mehler-like reaction. Our model posits that the Mehler-like reaction, by being crucial for generation of *pmf* upon increases in light intensity, promotes its own inactivation by pH-dependent disassociation from the thylakoid membrane. This ensures that the Mehler-like reaction is downregulated in order not to waste energy when a release valve for excessive electrons in the PETC is not needed. By shedding light on the intricate mechanisms that optimise photosynthetic efficiency, our study provides a foundation for rationally directing electron flux toward desired reactions in photosynthesis-based biotechnological applications.

## Methods

### Creation of *Synechocystis* BiFC strains

BiFC is prone to false positive results, especially when examining interactions of overexpressed proteins in a small compartment such as a cyanobacterial cell (Kudla and Bock 2016). To ameliorate this disadvantage, we used a Venus YFP variant where a I152L mutation in the N-terminal fragment decreases the propensity of the two fragments to self-reassemble, substantially decreasing the number of false positives (Kodama and Hu 2010). Proteins of interest were fused to the N or C terminal Venus-fragment via a short linker sequence to allow increased flexibility. We co-expressed the fusion proteins under IPTG-inducible lac2 promoters and S3 ribosome binding sites (RBS) of the *cpcB* gene (encoding the phycocyanin subunit β) from a single pDF-lac2 plasmid (Thiel et al. 2019). RBS S3 has been shown to enable strong expression of YFP (Thiel et al. 2019).

To create the BiFC strains, the *Synechocystis* sp. PCC 6803 coding sequences of *flv1* (sll1521), *flv2* (sll0219), *flv3* (sll0550), *flv4* (sll0217), *petF* (sll0020), *petH* (slr1643) full sequence coding for FNR_L_, and *petH* nucleotides 337–1242 coding for FNR_S_, as well as N-terminal (VN; nucleotides 1–468) and C-terminal (VC; nucleotides 469–724) fragments of the Venus I152L variant(Kodama and Hu 2010) were synthetized by GenScript (Piscataway, NJ, USA). In all cases a CAT codon coding for an extra Histidine residue was added to the sequences after the first ATG start codon in order to generate NsiI restriction sites (ATGCAT) at 5’ ends of the constructs. The 3’ stop codons were removed and chloramphenicol resistance (Cm^R^) cassettes with an NheI site at the 5’end and an NheI as well as a XhoI site at the 3’end were added. For *flv3*, a synonymous A1579T mutation was introduced to remove an endogenous NheI restriction site. SpeI restriction sites (ACTAGT) as well as flexible short linker sequences (CTGCCGGGCCCGGAGCTGCCG) were added at the 5’ends of the Venus fragments, while CmR cassettes flanked by NheI sites and followed by a SalI site (GTCGAC) were added at the 3’ ends. All *Synechocystis* gene sequences were retrieved from the CyanoBase database (Nakamura 1998).

A modular cloning strategy as described in (Thiel et al. 2019) was then used to assemble the gene-VN/VC expression vectors in the pDF-lac2 plasmid. First, the synthesised gene-Cm^R^ fragments were subcloned as NsiI-XhoI fragments downstream of a S3 RBS in pNiv carrier plasmids. The plasmid was then cleaved with NheI and XhoI, removing the Cm^R^. A VN- or VC-Cm^R^ fragment was inserted in frame to the 3’ of the first gene by SpeI/SalI cloning. The resultant pNiv-S3-gene 1-VN/VC-Cm^R^ plasmid was then cut with NheI/SalI and a S3-gene 2-VN/VC-Cm^R^ construct was inserted as a SpeI/SalI fragment downstream of the gene 1-VN/VC construct in the pNiv-S3 plasmid. Finally, constructs containing the two genes fused to VC or VN were transferred as SpeI/SalI fragments into pDF-lac2 expression plasmids. Wild-type *Synechocystis* sp. PCC 6803 cells were then transformed with the generated pDF-lac2 constructs (Fig. S27). Transformants were selected on BG-11 plates containing increasing concentrations of chloramphenicol and spectinomycin, up to 35 and 50 µg/ml, respectively. The presence of the transgenes in the strains was verified by PCR using primers against on the 5’ side (TGAGCGGATAACAATTTCACACAGAATT) and the 3’ side of the cloning site in pDF-lac2 (GACCGCTTCTGCGTTCTGATTTAAT).

### Creation of the ΔFed10 and ΔFed11 mutants

Three fragments of the *Fed10* (sll1584) and *Fed11* (ssl3044) (Artz et al. 2020) genes were amplified by PCR. These were the respective up- (primer 1 and 2) and downstream (primer 5 and 6) regions as well as the kanamycin antibiotic resistance cassette (primer 3 and 4) (Supplemental Table 1). Subsequently, constructs were generated by Gibson cloning (Gibson et al. 2009), assembling the three fragments into the pBluescript SK(+) vector. After sequencing the plasmids were transformed into *Synechocystis* cells as described (Williams 1988). Resulting transformants were checked by PCR after several rounds of segregation (Fig. S6A).

### Other strains and culture conditions

We used the glucose-tolerant *Synechocystis* sp. PCC 6803 WT strain (Williams 1988), mutant strains ΔFed2/Fed2, ΔFed3, ΔFed4, ΔFed5/Fed5, ΔFed6, ΔFed7Fed8Fed9, ΔFed9 (Wang et al. 2022), the ΔFlv3 strain (Helman et al. 2003), the ΔNdhD1NdhhD2 (ΔD1D2) strain (Ohkawa et al. 2000), the ΔFlv3 D1D2 strain (Nikkanen et al. 2020) as well as the ΔFNR_L_ (FSI) and ΔFNR_S_ (MI6) strains, deficient in the large and small isoform of FNR, respectively (Thomas et al. 2006).

Pre-experimental cultures and the *Synechocystis* BiFC strains were grown in 30 ml of BG-11 medium pH 7.5 with agitation under 3% [CO_2_] at 30°C under continuous illumination of white light of 50 µmol photons m^−2^s^−1^. Mutant pre-cultures were supplemented with appropriate antibiotics. Cells were harvested at the logarithmic growth phase and re-suspended in fresh BG-11 without antibiotics at OD_750_=0.2. Cultures were then moved to air [CO_2_] / 30°C and illuminated continuously with white light at 50 µmol photons m^−2^s^−1^.

### Membrane-inlet mass spectrometry

Exchange of ^16^O_2_ (*m/z*=32) and ^18^O_2_ (*m/z*=36) was measured *in vivo* with membrane-inlet mass spectrometry (MIMS) as described in (Mustila et al. 2016). After harvesting, cells were resuspended in fresh BG-11 pH 7.5, adjusted to 10 µg Chlorophyll (Chl) *a* ml^−1^, and kept for one hour in air [CO_2_] under 50 µmol photons m^−2^s^−1^. Cells were supplemented with ^18^O_2_ at an equivalent concentration to ^16^O_2_ and with 1.5 mM NaHCO_3_, and dark-adapted for 15 min, after which O_2_ exchange was monitored during a 5 min period of illumination with 500 μmol photons m^-2^s^-1^ of white actinic light. O_2_ exchange rates were calculated as described by (Beckmann et al. 2009).

### Bimolecular fluorescence complementation (BiFC) tests

The BiFC strains were grown in BG-11 pH 7.5 supplemented with 35 µg/ml of chloramphenicol and 50 µg/ml of spectinomycin under 3% [CO_2_] at 30°C under continuous illumination of 50 µmol photons m^−2^s^−1^ of white light until OD_750_ was c.a. 0.5. OD_750_ was then adjusted to 0.25 with standard fresh BG- 11 pH 7.5 without antibiotics, BG-11 without added sources of Mg^2+^ or Ca^2+^ cations, or BG-11 supplemented with 25 mM MgCl_2_ and 30mM CaCl_2_. Cultures were supplemented with 1 mM IPTG to induce expression of the BiFC fusion proteins and grown, harvested, and resuspended in 2 ml of fresh BG-11. Finally, Venus fluorescence and chlorophyll autofluorescence were imaged with a Zeiss LSM880 laser-scanning confocal microscope using a 63x or 100x objective with excitation at 488 and 543 nm, and detection at 518–621 nm. 30 µM CCCP, 20 µM Nigericin, 20 µM DCMU, or 40 µM Valinomycin was added to the cell suspensions before imaging when appropriate and as described in the figure legends. Identification of all cells from micrographs and mapping of the Venus and chlorophyll fluorescence signals within the was performed by a custom script for the Fiji software. The script is available at GitHub: https://github.com/gekoneCAC/CellHorizontalProfile. Quantification of Venus and Chl fluorescence co-localisation was performed using the EzColocalization plugin for the Fiji software (Stauffer et al. 2018). Costes’ threshold was used to calculate the Pearson correlation coefficient. An independent experiment constitutes a cyanobacterial culture started at a separate time and grown independently in a flask. As additional negative controls, we also performed BiFC tests with “empty” vectors, expressing a protein of interest fused to one Venus fragment together with a non-fused corresponding Venus fragment (e.g. Flv2-VC+empty-VN). Such constructs have been reported to be suboptimal controls in plants (Kudla and Bock 2016), but in our cyanobacterial BiFC system, only low-intensity Venus fluorescence was emitted by these strains (Fig. S28).

### NADPH fluorescence changes

NADPH fluorescence changes between 420 and 580 nm were monitored with a Dual-PAM 100 spectrophotometer and its 9-AA/NADPH accessory module (Walz) (Schreiber and Klughammer 2009). Cells were harvested, resuspended in fresh BG-11 and adjusted to 5 µg Chl *a* ml^−1^. Cells were dark-adapted for 15 min prior to measurement of NADPH fluorescence changes under 500 μmol photons m^- 2^s^-1^ illumination for 40 s, followed by 40 s in darkness.

### Near-infrared spectrophotometry

*In vivo* redox changes of P700, PC, and Fed were deconvoluted from absorbance differences at four near-infrared (NIR) wavelength pairs at 780–820, 820–870, 840–965 and 870–965 nm measured with a DUAL-KLAS-NIR spectrophotometer (Walz) differential model plot (DMP) used to deconvolute the Fed signal was determined from ΔFlv3 D1D2 triple mutant cells as described in (Nikkanen et al. 2020), while the P700 and PC DMPs were determined as described by (Theune et al. 2021). Cells were grown as for the BiFC tests, harvested, resuspended in fresh BG-11 and adjusted to 20 µg Chl *a* ml^−1^. Cells were dark-adapted for 15 min prior to measurement. We used the NIRMAX script (Klughammer and Schreiber 2016) consisting of a 3 s illumination with strong AL (3000 μmol photons m^−2^ sec^−1^), with a saturating pulse after 200 ms of illumination to fully reduce the Fed pool. After 4 s of darkness, cells were illuminated under far red (FR) light for 10 s with a saturating pulse at the end to fully oxidise P700 and PC. The maximum reduction values of Fed and maximum oxidation values of P700 and PC thus obtained were used to normalise the traces.

### Electrochromic shift (ECS) measurements

Light-induced changes in the *pmf*, thylakoid conductivity (gH+) and thylakoid proton flux (vH+) were determined by monitoring the dark interval relaxation kinetics (DIRK) of the absorbance difference between 500 and 480 nm, which constitutes the ECS in *Synechocystis* (Viola et al. 2019), as described previously (Nikkanen et al. 2020). Cells were harvested and resuspended in fresh BG-11, low cation BG-11, or high cation BG-11 (see above) pH 7.5 at a Chl concentration of 7.5 µg/ml. Suspensions were illuminated at 500 μmol photons m^-2^s^-1^ with 600 ms dark intervals as indicated in the Figure legends, and 500-480 nm absorbance changes were detected using a JTS-10 spectrophotometer (BioLogic) and appropriate interference filters (Edmund Optics).

### Acridine orange (AO) and acridine yellow (AY) fluorescence measurements

To monitor the light-induced alkalisation of the cytosol, AO and AY fluorescence changes (Teuber et al. 2001) were recorded using a Dual-PAM-100 spectrophotometer and the AO/AY emitter-detector module (Schreiber and Klughammer 2009). Cells were grown in air level CO_2_ in BG-11 pH 7.5 under 50 µmol photons m^-2^s^-1^ for four days, harvested, and suspended in fresh BG-11 with Chl concentration adjusted to 5 µg/ml. 5 µM AO or AY, and 1mM KCN (when appropriate) was added before a 10 min dark-adaptation prior to measurements under 216 or 500 µmol photons m^-2^s^-1^, as specified in the figure legends.

### Immunoblotting

Total proteins were extracted from BiFC cultures used for microscopy as described by (Zhang et al. 2009). For the analysis FDP subcellular localization, WT *Synechocystis* cells were grown for 4 days under 50 µmol photons m^-2^ s^-1^ in air-level [CO_2_] at 30°C. Samples were taken from light, or dark-adapted for 20 min, or taken from light and supplemented with 30 µM CCCP, then quickly frozen in liquid N_2_. Soluble and membrane fractions were isolated as described by (Zhang et al. 2009). Proteins were separated by sodium dodecyl sulphate (SDS) polyacrylamide gel electrophoresis on Mini-Protean TGX 4-15% gels (BioRad) and transferred to polyvinylidene fluoride membranes. Membranes were probed with primary antibodies against the N-terminal fragment of YFP (Origene), Flv1 and Flv4 (Genscript), Flv2 and Flv3 (Antiprot), and PetH (kindly shared by H. Matthijs). Horseradish peroxidase (HRP) -conjugated goat-anti mouse (Bio-Rad) and goat-anti-rabbit (GE Healthcare) secondary antibodies as well as ECL (Amersham) were used for detection.

### In silico modelling

The sequences of *Synechocystis* sp. PCC 6803 Flv1 (sll1521), Flv2 (sll0219), Flv3 (sll0550), and Flv4 (Sll0217) proteins were retrieved from UniProt using the following IDs: P74373, P72723, Q55393, and P72721, respectively. The crystal structures for the multiple structure-based alignments were retrieved by BLAST searches from the Protein Data Bank (Berman 2000) using these FDP sequences as queries. The β-lactamase-like and flavodoxin domains of the Flv1 structure (PDB ID: 6H0D) were separately superimposed on the methanogenic archaea *Methanothermobacter marburgensis* F_420_H_2_ oxidase structure (PDB IDs: 2OHI, 2OHJ Seedorf et al. 2007) to create structure-based sequence alignment and find out the correspondence between the amino acids in the two proteins. Thereafter, Flv1 and Flv3 flavodiiron core sequences were aligned to the prealigned structure-based alignment. The 3D models for the open and closed state of the core composed of β-lactamase-like and flavodoxin domains of Flv1, Flv2, Flv3, and Flv4 monomers, as well as Flv1/Flv1 and Flv3/Flv3 homooligomers, and Flv1/Flv3 and Flv2/Flv4 heterooligomers were built based on the alignment and the crystal structure of F_420_H_2_ oxidase from *M. marburgensis* (Seedorf et al. 2007) (PDB IDs: 2OHI for closed conformation and 2OHJ for open conformation) (Seedorf et al. 2007) using MODELLER v.10.4 (Webb and Sali 2016), generating ten models per each model.

All sequence alignments were done with the MALIGN tool within the Bodil modeling suite (Lehtonen et al. 2004), and visualized using ESPript 3.0R (Robert and Gouet 2014). In all cases, ligands and structural waters have been directly transferred from the templates during modelling. The MODELLER models with the lowest zDOPE and Molpdf scores were selected for further analysis. Model quality was assessed using the Protein Reliability Report feature in Schrodinger Maestro 13.3. Following protein preparation at pH 7.0 and pH 8.5 using the “prepwizard” script, the net charges were calculated in triplicate using the “calc_protein_descriptors” script in Schrodinger Bioluminate (Schrödinger Release 2023-1 2021a). The electrostatic potential surfaces for FDP monomers and oligomers were visualised with the Poisson-Boltzmann ESP module in Schrodinger Maestro (Schrödinger Release 2023-1 2021b).

The amino acid sequences of YFP-labeled *Synechocystis* Flv1 and Flv3 used in the experimental part of the study were used to prepare the full-size model of the Flv1-VN/Flv3-VC complex. The YFP-labeled Flv1/Flv3 heterodimers were modeled in two variants – with VN and VC fragments of YFP attached to either Flv1 or Flv3. The structure of Venus Yellow Fluorescent Protein (YFP) was modelled using the AlphaFold2 Multimer Google Collab notebook (Mirdita et al. 2022) with the following settings: 48 recycles, pdb100 template mode, and Amber relaxation of the best model. For modeling the C-terminal flavin reductase domain of Flvs, the crystal structure of *Thermus thermophilus* putative flavoprotein (PDB ID: 1WGB) was used as a template. Since *T. thermophilus* putative flavoprotein is a monomer, we overlaid the monomeric models onto the dimeric structure of *T. thermophilus* probable flavoprotein HB8 [TS1] (Imagawa et al., 2005) (PDB ID: 1YOA[SV2]) to create dimers. The relative spatial arrangement of the previously modeled flavodiiron core and the C-terminal flavin oxidase domain was predicted using the ClusPro webserver (Kozakov et al. 2017). Finally, to reconstruct the full-length YFP-labeled models, the separate models of the flavodiiron core, the docked C-terminal flavin-reductase domains, and AF2-modeled YFP obtained on the first stage were used as templates for multiple-template modelling in MODELLER. The models were subjected to 500-step restrained energy minimisation, followed by 100-ns molecular dynamics simulations in an explicit solvent with Desmond (Schrödinger 2022a) to explore the conformations of long interdomain loops. The models with the lowest free energy were selected for visualisation using Pymol.

### Statistical analysis

Two-sample Student’s t-tests or one-way ANOVA, as indicated in Figure legends, were performed in OriginPro 2024 to determine significant differences in maximal O_2_ photoreduction rates and PCC’s of fluorescence signal co-localisation in BiFC images. One-sample t-tests were used to determine significant difference to control samples in average Western blot band intensity (with control sample band intensity normalised to 1). P<0.05 was interpreted as statistically significant. Results of the statistical tests are reported in more detail in Tables S2 and S3. The meanings of bar charts, line charts, and error bars are indicated in the Figure legends. A culture grown independently in a flask for at least four days was considered a biological replicate.

### Accession numbers

Sequence data from this article can be found in the EMBL/GenBank data libraries under accession numbers *sll1521* (Flv1), *sll0219* (Flv2), sll0550 (Flv3), *sll0217* (Flv4), *sll0020* (Fed1), *sll1382* (Fed2), *slr1828* (Fed3), *slr0150* (Fed4), *slr0148* (Fed5), *ssl2559* (Fed6), *sll0662* (Fed7), *ssr3184* (Fed8), *slr2059* (Fed9), *sll1584* (Fed10), *ssl3044* (Fed11), *slr0331* (NdhD1), *slr1291* (NdhD2) and *slr1643* (FNR).

## Supplementary Material

**Figure S1**: O_2_ evolution, O_2_ uptake, and CO_2_ fixation in ΔFlv3 and ΔFNR mutant strains.

**Figure S2.** NAD(P)H fluorescence changes under 3% [CO_2_] and low irradiance.

**Figure S3.** Redox kinetics of PSI electron carriers in WT, ΔFlv3, ΔFNR_L_ and ΔFNR_S_ cells.

**Figure S4:** BiFC tests between FNR and Flv2

**Figure S5:** Verification of expression of BiFC fusion constructs by immunoblotting

**Figure S6:** Characterisation of Fed mutants

**Figure S7:** Redox changes of PC, P700, and Fed in WT, ΔFed, and ΔFlv3 cells.

**Figure S8**. BiFC test for interaction between Flv3-VN and Fed1-VC.

**Figure S9**. BiFC tests for interactions between Fed9 and Flv1 or Flv3

**Figure S10.** Differently localising subpopulations of Flv1-VN / Flv3-VC and Flv2-VC / Flv4-VN interactions in BiFC.

**Figure S11:** Effect of CCCP on the *pmf* in *Synechocystis*

**Figure S12:** Generation of the *pmf*, thylakoid conductivity (gH+), and proton flux (vH+) during dark-to-light transitions in low vs high cation concentration

**Figure S13.** Effect of Nigericin and DCMU on ΔpH generation during induction of photosynthesis

**Figure S14**. Effect of Nigericin on the subcellular localisation of Flv1/3 and Flv2/4 interactions

**Figure S15.** Effect of DCMU on the subcellular localisation of Flv1/3 and Flv2/4 interactions

**Figure S16.** Immunodetection of the subcellular localization of FDPs

**Figure S17:** Effect of CCCP on O_2_ uptake

**Figure S18.** Effect of Nigericin on O_2_ uptake

**Figure S19:** Cytosolic alkalisation upon illumination

**Figure S20**. Effect of DTT, CuCl_2_, and KCN on photosynthesis

**Figure S21**. Effect of DTT, CuCl_2_, and KCN on the subcellular localisation of Flv2/4 interactions

**Figure S22**. Effect of DTT, CuCl_2_, and KCN on the subcellular localisation of Flv1/3 interactions

**Figure S23:** Effect of cation concentration on the subcellular localisation of Flv1/3 and Flv2/4 interactions

**Figure S24:** BiFC tests for Flv1 and Flv3 self-interactions and between Flv2 and Flv3

**Figure S25:** Effect of cation concentration and *pmf* uncoupling on the subcellular localisation of Flv3 self-interactions

**Figure S26:** *In silico* analysis of electrostatic surface charges of FDP monomers under pH 7.0 and pH 8.5

**Figure S27:** Plasmid map of a BiFC construct to express *Synechocystis flv1* and *petF* (Fed1) as fusion proteins with N- and C-terminal Venus fragments, respectively

**Figure S28.** “Empty” vector control experiments for BiFC.

**Table S1:** Primers used for generating the ΔFed10 and ΔFed11 strains

**Table S2.** Results from statistical tests in the main article

**Table S3**. Results from statistical tests in Supplementary Material

## Acknowledgments

We thank Pauli Kallio for kindly providing plasmids for modular cloning and for helpful discussions, Laura Wey for valuable discussions, and Anniina Lepistö, Janette Vähäsarja, Sofia Westerlund, Ville Käpylä, Mahfuzur Rahman, and Linda Nevala for technical assistance. Biophysical experiments were performed within the PHOTOSYN infrastructure at the University of Turku, and confocal microscopy at the Cell Imaging and Cytometry Core, Turku Bioscience Centre, with the support of Biocenter Finland. We also thank the bioinformatics (J.V. Lehtonen), translational activities, and structural biology FINStruct infrastructure support from Biocenter Finland and CSC IT Center for Science for computational infrastructure support at the Structural Bioinformatics Laboratory, (SBL) Åbo Akademi University.

## Author contributions

L.N. and Y.A. conceived the idea and designed the experiments. L.N. performed most experiments and analysed the data. S.V. and T.A.S performed the computational modelling. A.S.S. and M.H. performed additional cloning, generation of BiFC strains and MIMS. G.K. created tools for microscopy image analysis. Y.W. and M.B. generated the Fed mutants. K.G. and Y.A. provided resources. L.N. wrote the initial draft, L.N. and Y.A. finalised the article with contributions from all authors.

## Data and materials availability

All data are available in the main text or the supplementary materials. Source data and raw microscopy images are available from the corresponding author upon reasonable request.

