## Supplementary Information for "Flavodiiron proteins associate pH-dependently with the thylakoid membrane for ferredoxin-1 powered O_2_ photoreduction"

Lauri Nikkanen *et al.*

**This PDF file includes:**

Figs. S1 to S28
Tables S1 to S3

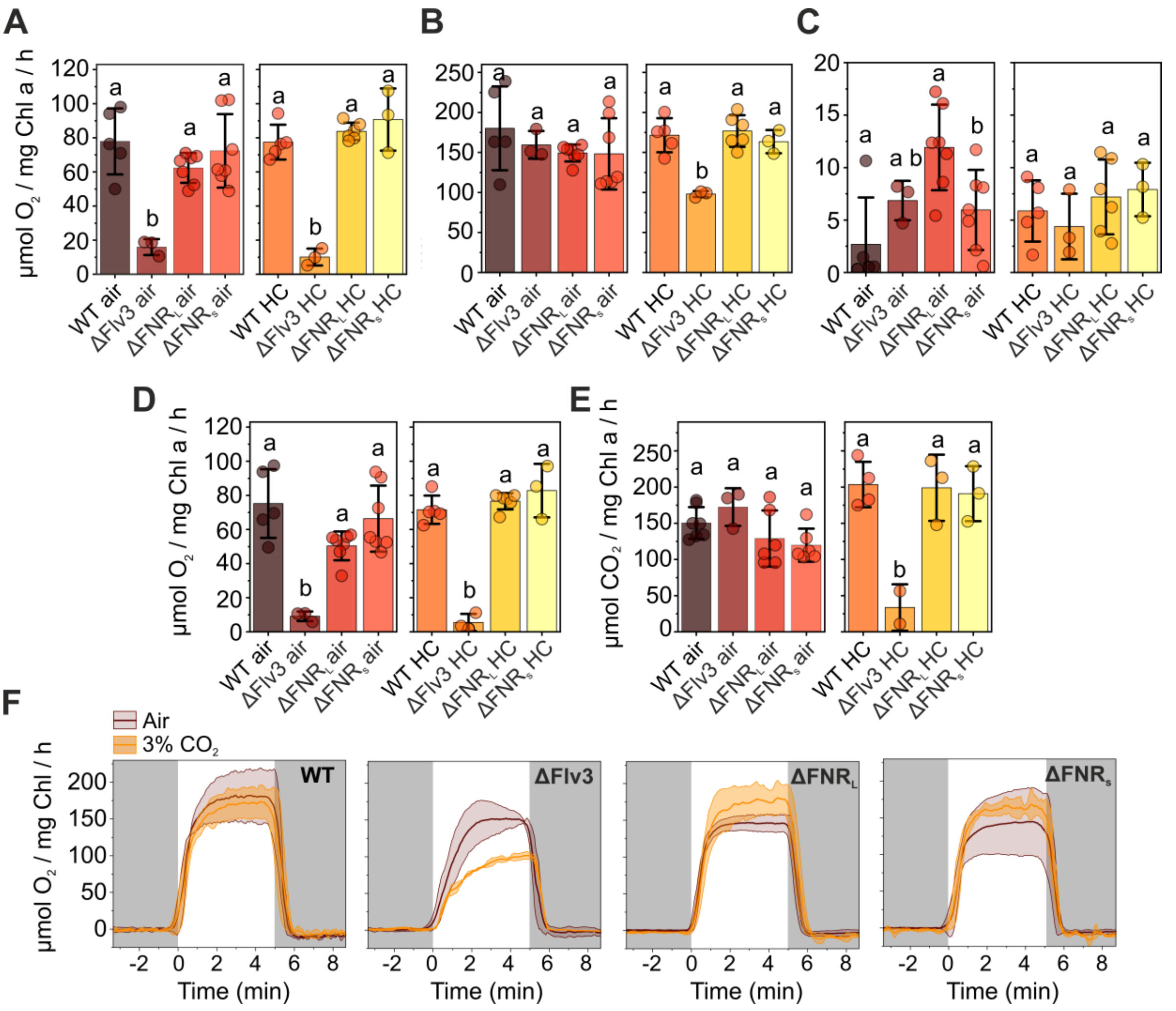


**Figure S1. $O_2$ evolution, $O_2$ uptake, and $CO_2$ fixations in ΔFlv3 and ΔFNR mutant strains.** Supports Figure 1.

(A) Total $O_2$ uptake maximum at the peak during the first minute of illumination as measured by MIMS. (B) Maximum gross $O_2$ evolution during 5 min of illumination. (C) Dark respiration, measured as the average $O_2$ uptake during 2 min in darkness. (D) Maximum $O_2$ photoreduction (D=A-C). (E) Steady state $CO_2$ fixation. Calculated as average $CO_2$ consumption during the last minute of illumination + average $CO_2$ production during first minute of darkness prior to illumination ($CO_2$ release from respiration). Values are from the same MIMS experiments as those in Figure 1C–D. For air-level cultures, the values are averages of 5 (WT), 3 (ΔFlv3), or 7 ($ΔFNR_L$ and $ΔFNR_S$) biological replicates +/- standard deviation in A-D and 7 (WT), 3 (ΔFlv3), or 6 ($ΔFNR_L$ and $ΔFNR_S$) biological replicates in E. For elevated [$CO_2$], values are averages of 5 (WT, 3 (ΔFlv3 and $ΔFNR_S$), and 6 ($ΔFNR_L$) biological replicates in A–D and 4 (WT), 2 (Flv3), or 3 ($ΔFNR_L$ and $ΔFNR_S$) in E. Individual replicate data points are shown as circles. Statistical significance of mean differences was tested by one-way ANOVA and Tukey's post-hoc tests with $p>0.05$ regarded as significant. (F) Gross $O_2$ evolution in air-level and in 3% [$CO_2$]. $O_2$ gas fluxes were measured by MIMS, and traces are derived from the same measurements as A–E and Fig. 1C–D.

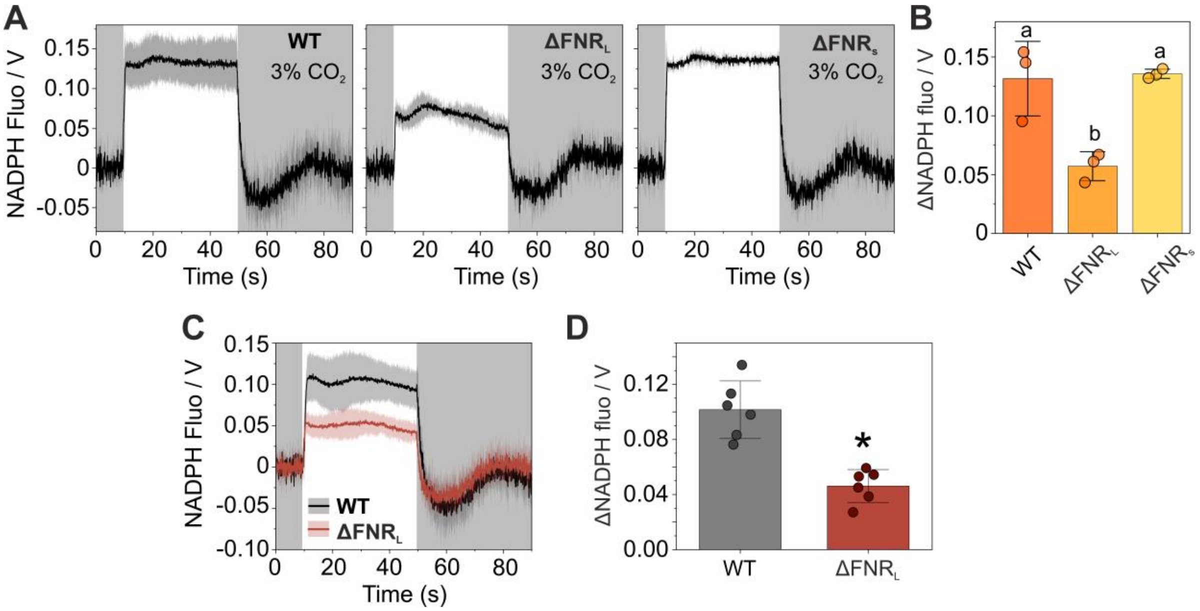


**Figure S2. NAD(P)H fluorescence changes under high [$CO_2$] and low irradiance.** Supports Figure 1.

**(A)** Light-induced NADPH fluorescence changes in WT, $\Delta FNR_L$, and $\Delta FNR_S$ cells grown under 3% [$CO_2$] during 40 s illumination at 500 µmol photons $m^{-2}$ $s^{-1}$ and subsequent darkness. Averaged traces from three biological replicates ± standard deviation (shadowed area) are shown. Dark-adapted NADPH fluorescence levels were set to 0.

**(B)** Quantification of light-induced change in the NADPH fluorescence signal in the experiments for (A), calculated as mean fluorescence level between 30 and 35 sec of illumination – mean fluorescence level in the pre-illumination dark period. Values are averages from three biological replicates ± standard deviation with individual data points shown as circles. Statistical significance was tested by one-way ANOVA and a Tukey's post-hoc test for comparison of means. Means that do not share a grouping letter (a, b) are significantly different ($P<0.05$).

**(C)** Light-induced NADPH fluorescence changes in WT and $\Delta FNR_L$ cells during 40 s illumination at low light (50 µmol photons $m^{-2}$ $s^{-1}$, 3% [$CO_2$]) and subsequent darkness. Averaged traces from six biological replicates ± standard deviation (shadowed area) are shown. Dark-adapted NADPH fluorescence levels were set to 0.

**(D)** Quantification of light-induced change in the NADPH fluorescence signal in the experiments for (C), calculated as mean fluorescence level between 30 and 35 sec of illumination – mean fluorescence level in the pre-illumination dark period. Statistical significance was tested by Student's two-sample t-test, with * denoting significant difference ($P<0.05$).

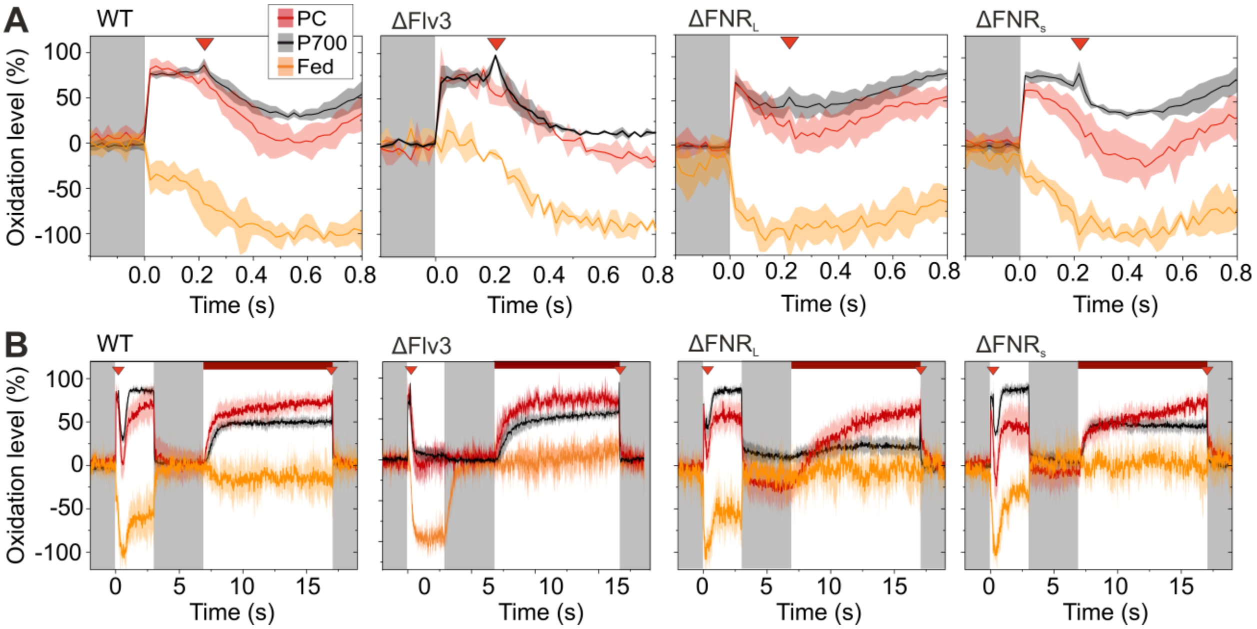


**Figure S3. Redox kinetics of PSI electron carriers in WT, ΔFlv3, $\Delta FNR_L$ and $\Delta FNR_S$ cells.** Supports Figure 1.

PC, P700, and Fed redox changes as measured with a DUAL-KLAS-NIR spectrophotometer. The traces are normalised to the maximal oxidation values of PC and P700 and maximal reduction of Fed, as determined with the NIRMAX protocol, of which first 0.8 of illumination with actinic light is shown in (A), including a multiple turnover pulse after 200 ms (indicated by the red triangles). The traces are averages of 5 biological replicates +/- standard deviation. (B) Full traces from the NIRMAX protocol, consisting of a 3 s illumination with 1750 µmol photons $m^{-2}s^{-1}$ actinic with a multiple turnover pulse after 200 ms (indicated by the red triangles) to fully reduce the Fed pool, 3 s darkness, followed by 10 s of illumination under far red light with another multiple turnover flash at the end of the illumination period to fully oxidise P700 and PC. The traces are normalised to the maximal oxidation values of PC and P700 and maximal reduction of Fed. In some replicates of the $\Delta FNR_L$ strain the far-red light with a multiple turnover flash did not result in full oxidation of P700. In those cases, the maximal P700 value was taken from the maximal oxidation induced by the strong actinic illumination (3–6 seconds). Averaged traces from 5 biological replicates are shown, with standard deviation as the shadowed area.

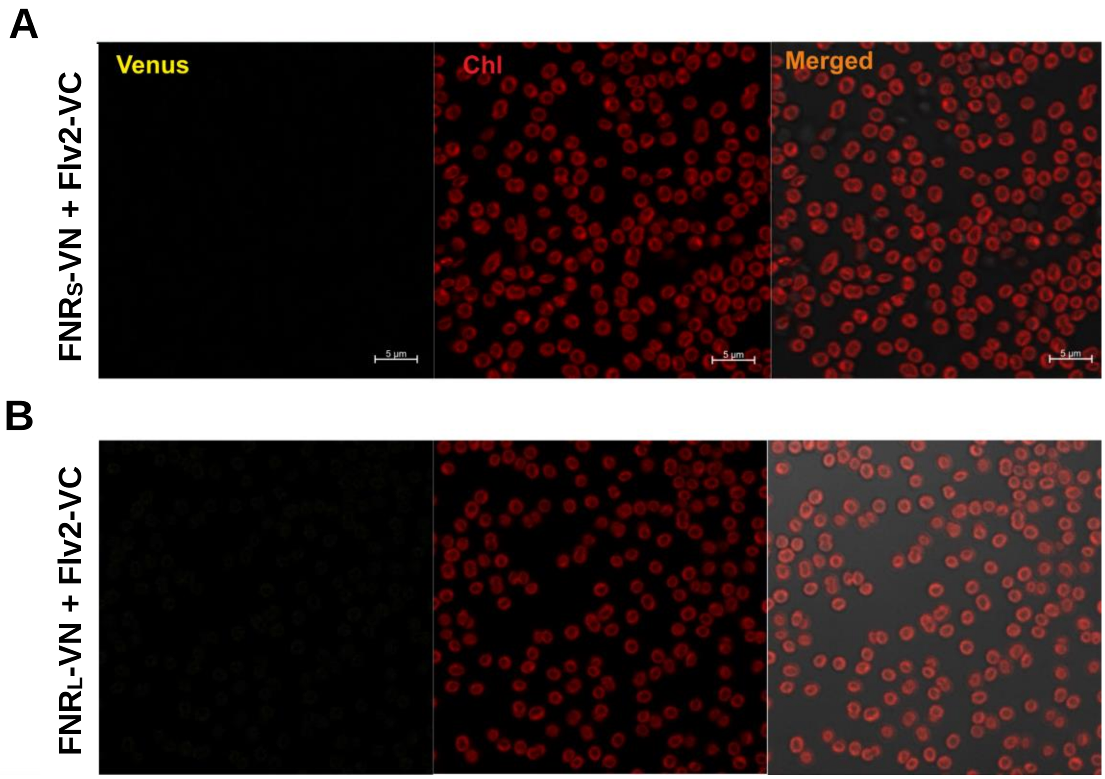


**Figure S4. BiFC tests between FNR and Flv2.** Supports Figure 2.

Representative confocal micrographs from 3 independent experiments are presented, showing fluorescence emanating from re-assembled Venus I152L fluorescent proteins in the left panel, chlorophyll a (Chl) autofluorescence from the thylakoid membranes in the middle panel middle, and overlaid Chl and Venus fluorescence in the right panel.
(A) BiFC test between $FNR_S$-VN and Flv2-VC.
(B) BiFC test between $FNR_L$-VN and Flv2-VC.

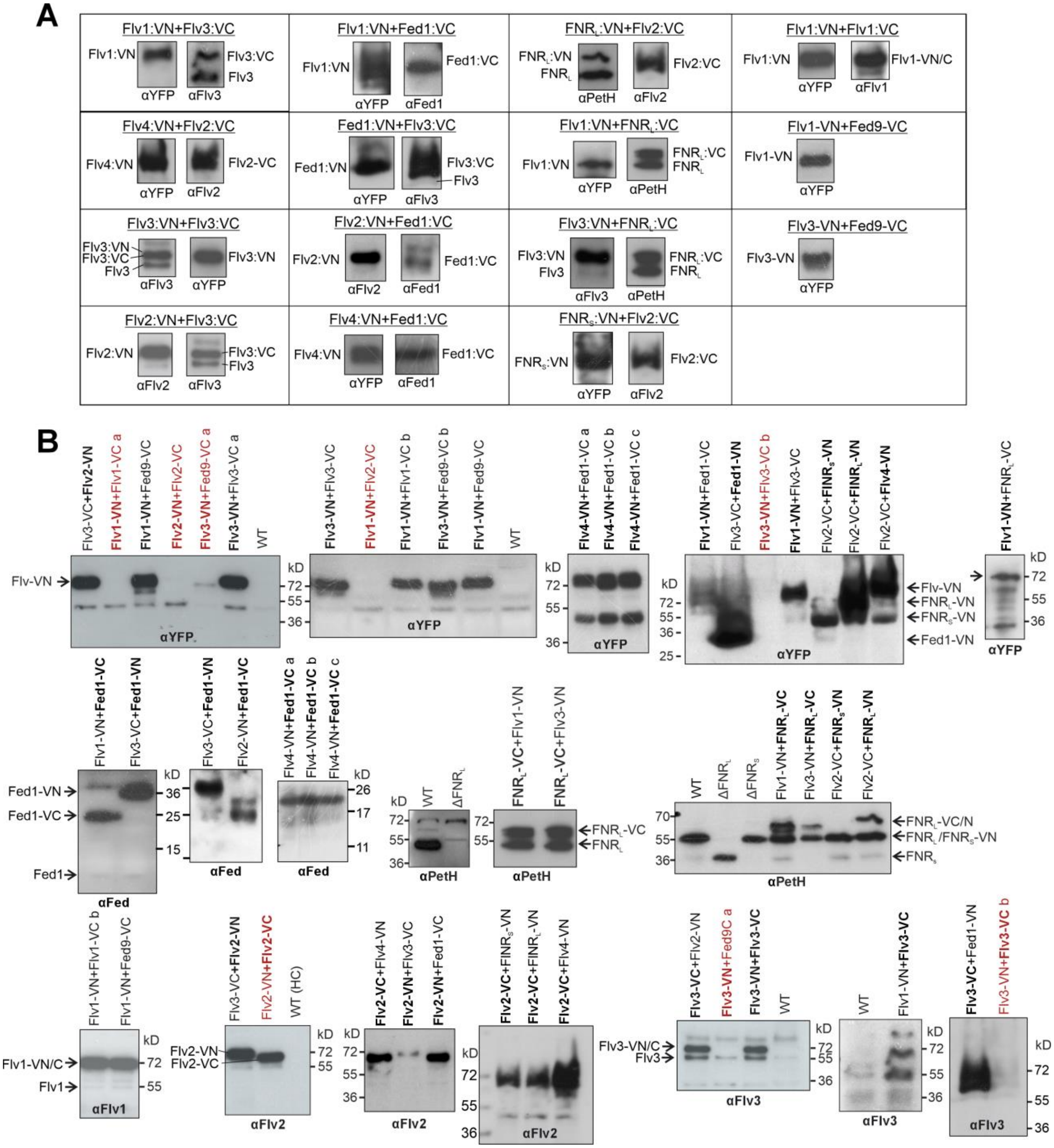


**Figure S5. Verification of expression of BiFC fusion constructs by immunoblotting.** Supports Figures 2, 4, 5, S2, and S15.

Total proteins were extracted from samples used for BiFC experiments, separated by SDS-PAGE and probed with specific antibodies against the N-terminal part of YFP, Fed, PetH (FNR), Flv1, Flv2, and Flv3. **(A)** Shows cropped portions of immunoblots to collect relevant bands for each BiFC strains, while **(B)** shows the original immunoblots with molecular weight markers. The strains marked with red colour represent BiFC clones that did not exhibit proper expression of one or both fusion proteins and where thus excluded from the experiments in this study. The Fed antibody could not recognize the Fed9-VC fusion protein.

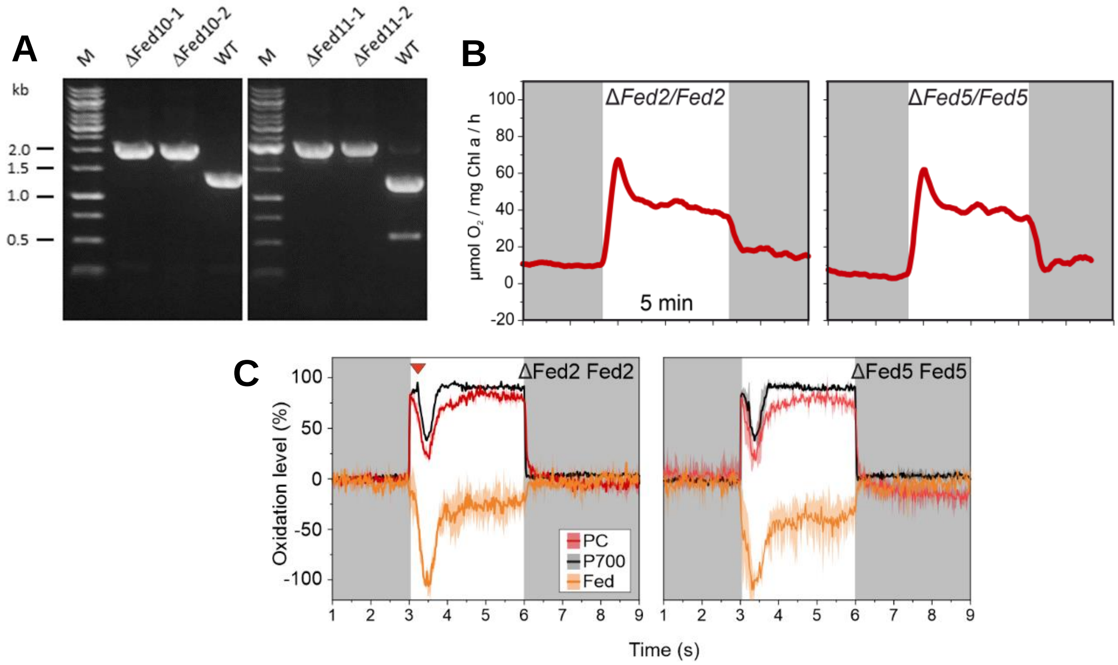


**Figure S6. Characterisation of Fed mutants.** Supports Figure 3.

(A) PCR analysis of the ΔFed10 and ΔFed11 mutant strains and the wild type (WT).
(B) Light-induced $O_2$ uptake in partially segregated Fed2 and Fed5-deficient cells. Experiments were performed as in Figures 1 and 2.
(C) Redox changes of PC, P700, and Fed in in partially segregated Fed2- and Fed5-deficient cells. Averaged traces from two biological replicates with standard deviation are shown. Experiments were performed as in Figures 1 and 3.

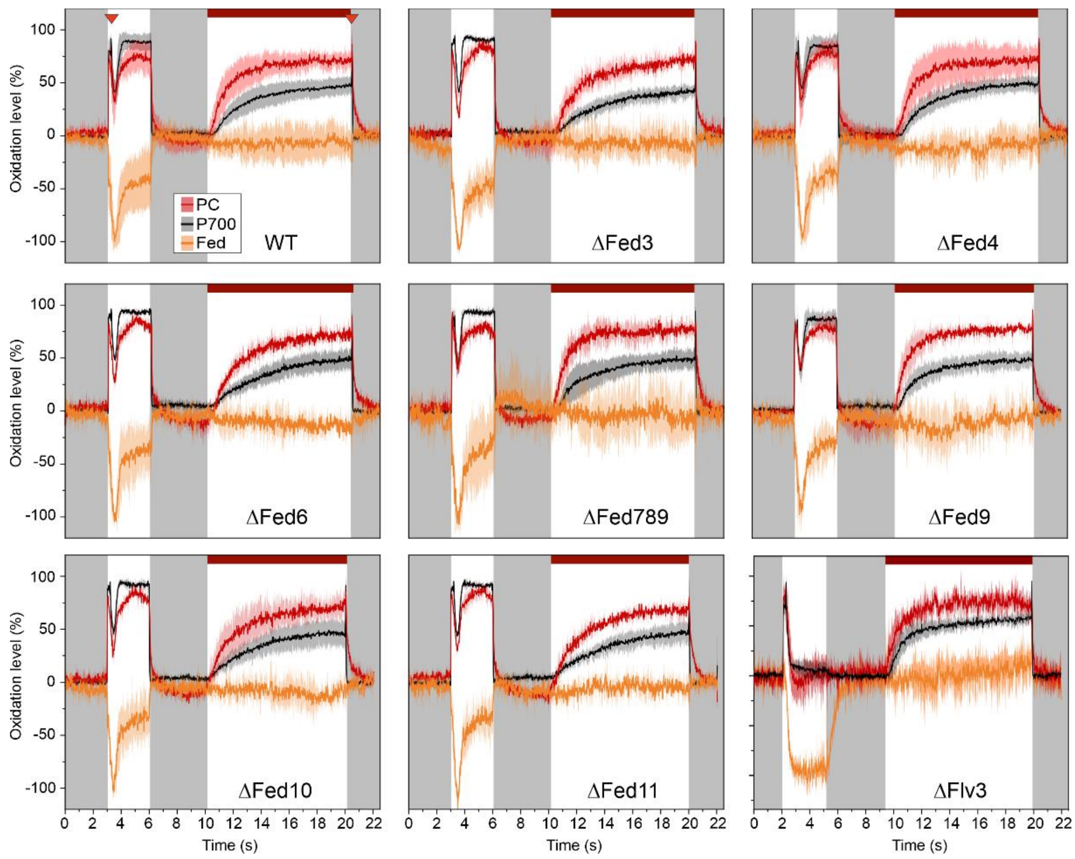


**Figure S7. Redox changes of PC, P700, and Fed in WT, ΔFed, ΔFlv3 cells.** Supports Figure 3.

Cells grown in air level $CO_2$ concentration for 4 days were harvested and resuspended in fresh BG-11 pH 7.5 with Chl concentration adjusted to 15 µg/ml. PC, P700, and Fed redox changes were deconvoluted from near-infrared absorbance differences measured with a DUAL-KLAS-NIR spectrophotometer. A modified NIRMAX protocol was used, consisting of a 3 s illumination with 1750 µmol photons $m^{-2}s^{-1}$ actinic with a multiple turnover pulse after 200 ms (indicated by the red triangle in the WT panel) to fully reduce the Fed pool, 3 s darkness, followed by 10 s of illumination under far red light with another multiple turnover flash at the end of the illumination period to fully oxidise P700 and PC. The traces are normalised to the maximal oxidation values of PC and P700 and maximal reduction of Fed. Averaged traces from 3–6 biological replicates are shown, with standard deviation as the shadowed area.

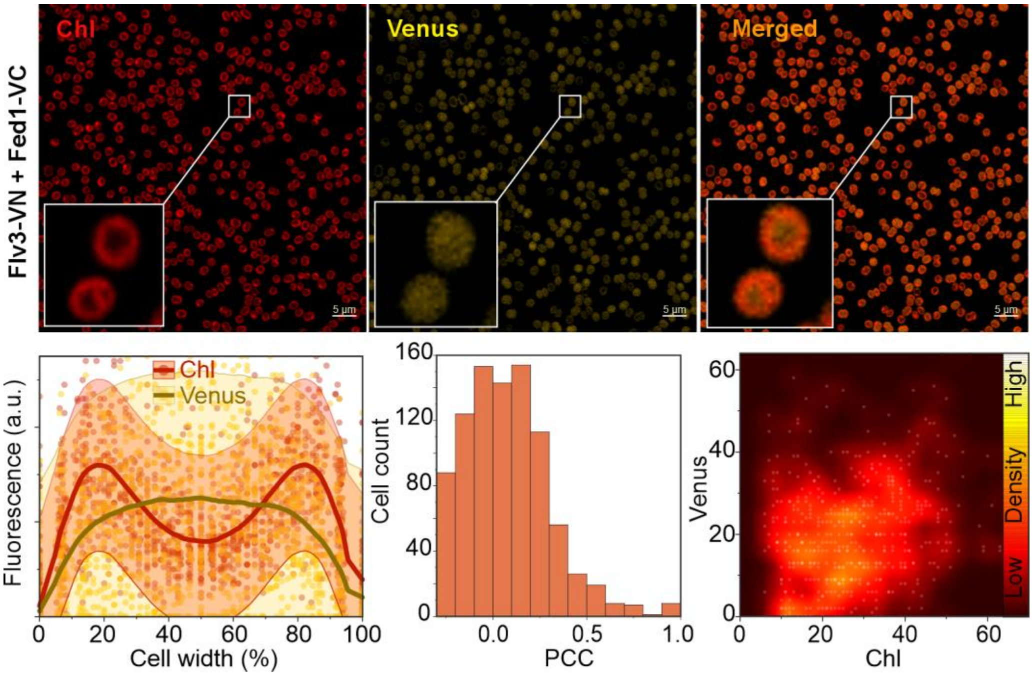


**Figure S8. BiFC test for interaction between Flv3-VN and Fed1-VC.** Supports Figure 4.

BiFC test between Flv3-VN and Fed1-VC. Upper panels in show representative confocal micrographs from the chlorophyll and Venus fluorescence channels as well as a merged image. The lower panels show co-localisation of Chl (red) and Venus (yellow) fluorescence intensities over cross-sections of 436 cells (with width normalised to 100%) shown as a scatterplot. Brighter red and yellow lines show moving regression curves produced by locally weighted scatterplot smoothing (LOWESS). The histograms show the distribution of the Pearson Correlation Coefficient (PCC) for Chl and Venus fluorescence co-localisation in all cells from three micrographs from individual replicates. The cytofluorograms show co-localised Chl and Venus fluorescence intensities as a scatter plot from all cells in four micrographs from independent replicates

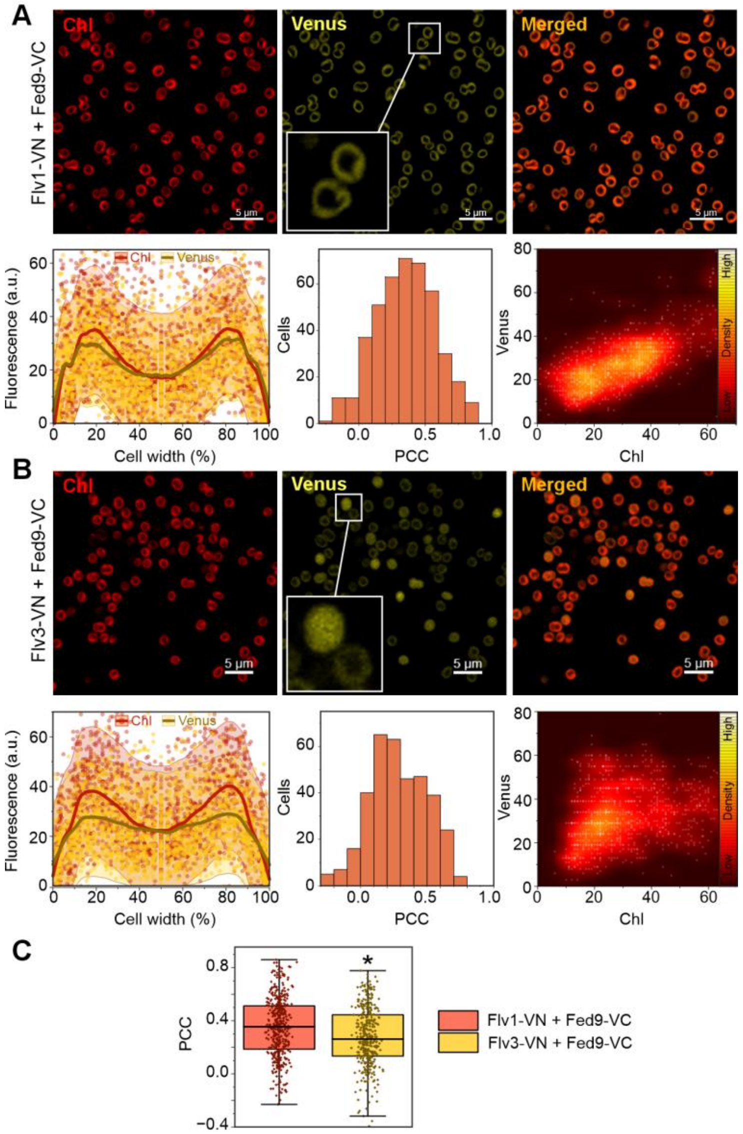


**Figure S9. BiFC tests for interactions between Fed9 and Flv1 or Flv3**. Supports Figure 4.

(A) BiFC tests between Flv1-VN and Fed9-VC. (B) BiFC tests between Flv3-VN and Fed9-VC. Upper panels in A-B show representative confocal micrographs from the chlorophyll and Venus fluorescence channels as well as a merged image. The lower panels show co-localisation of Chl (red) and Venus (yellow) fluorescence intensities over cross-sections of 277 (A) and 239 (B) cells (with width normalised to 100%) shown as a scatterplot. Brighter red and yellow lines show moving regression curves produced by locally weighted scatterplot smoothing (LOWESS). The histograms show the distribution of the Pearson Correlation Coefficient (PCC) for Chl and Venus fluorescence co-localisation in all cells from three micrographs from individual replicates. The cytofluorograms show co-localised Chl and Venus fluorescence intensities as a scatter plot from all cells in four micrographs from independent replicates. (C) PCC values from (A) and (B) plotted as box plots for comparison of means. * indicates statistically significant difference of mean, as determined by one-way ANOVA ($p<0.05$).

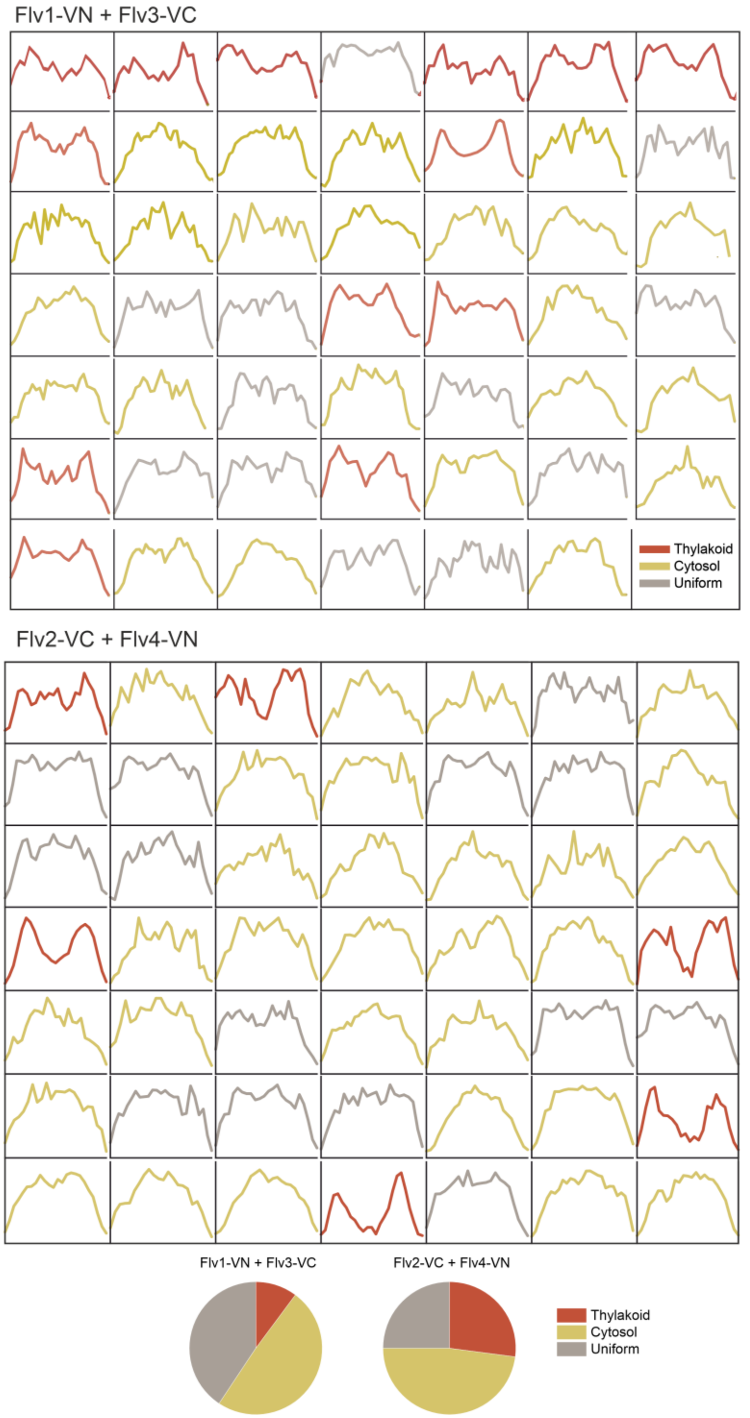


**Figure S10. Differently localising subpopulations of Flv1-VN / Flv3-VC and Flv2-VC / Flv4-VN interactions in BiFC.** Supports Figure 5.

Venus fluorescence cross-section profiles from 48 and 49 cells expressing the Flv1-VN and Flv3-VC or Flv2-VC and Flv4-VN BiFC fusion proteins. The data is taken from Figure 5A-B. The profiles are grouped into either thylakoid (red), cytosolic (gold), or uniform/unclear (grey) localisation, with proportion of cells in each group plotted in the pie charts at the bottom.

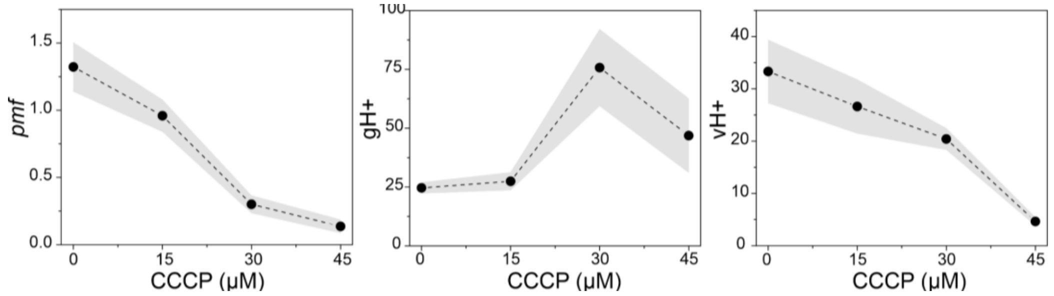


**Figure S11. Effect of CCCP on the *pmf* in *Synechocystis***. Supports Figure 5.

The magnitude of the *pmf* was determined from the dark interval relaxation kinetics (DIRK) of the ECS signal after illuminating WT *Synechocystis* cells f or 27 s at 500 µmol photons $m^{-2}$ $s^{-1}$ in the presence of 0, 15, 30, or 45 µM CCCP. *Pmf* is the extent of ECS signal decay during a DIRK, thylakoid proton conductivity (gH+) is the inverse of the time constant of a first order fit to DIRK kinetics, and thylakoid proton flux (vH+) is calculated as pmf*gH+. The values shown are averages from 3 (15 µM CCCP) or 4 (0, 30, and 45 µM CCCP) biological replicates +/- SEM.

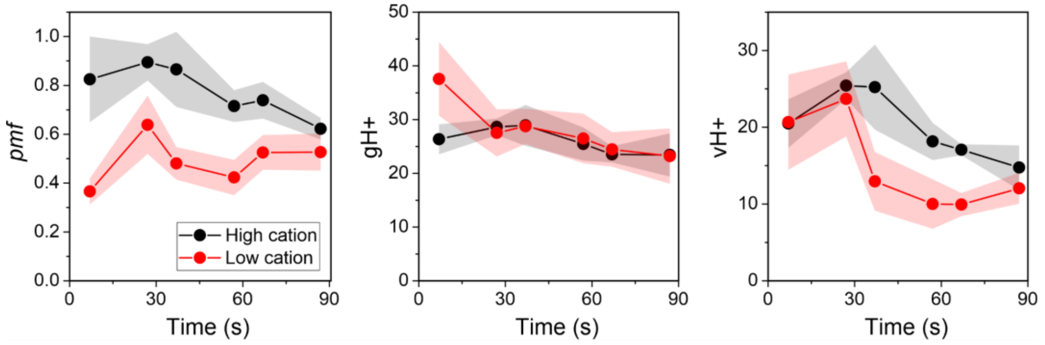


**Figure S12. Generation of the *pmf*, thylakoid conductivity (gH+), and proton flux (vH+) during dark-to-light transitions in low vs high cation concentration.** Supports Figure 5.

Dark-adapted WT *Synechocystis* cells grown for in BG-11 pH 7.5 in 3% $CO_2$ under 50 µmol photons $m^{-2}$ $s^{-1}$ were harvested and resuspended in fresh BG-11 medium containing either now added source of $Mg^{2+}$ and $Ca^{2+}$ (low cation) or in BG-11 supplemented with 25 mM $MgCl_2$ and 30mM $CaCl_2$ (high cation). Suspensions were illuminated for 90s with 500 µmol photons $m^{-2}$ $s^{-1}$ of green light, with 600 ms dark intervals administered after 7, 27, 37, 57, 67, and 87 s of light. *Pmf*, gH+, and vH+ were determined using DIRK analysis of the ECS signal. Values are averages from 4 biological replicates +/- SEM (shadowed area).

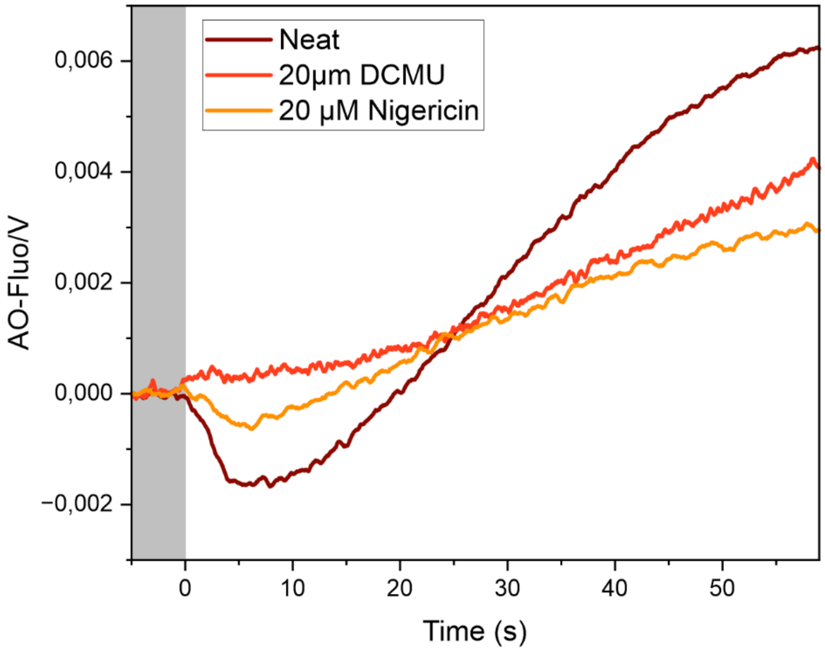


**Figure S13. Effect of Nigericin and DCMU on ΔpH generation during induction of photosynthesis.** Supports Figure 5.

(A) Acridine orange (AO) fluorescence was recorded from dark-adapted WT *Synechocystis* cultures with or without 20 µM Nigericin or 20 µM DCMU under 216 µmol photons $m^{-2}s^{-1}$ of actinic light for 1 min. Cultures were grown in BG-11 pH 7.5 in air-level $CO_2$ under 50 µmol photons $m^{-2}s^{-1}$ for 4 days before measurements. Chl concentration was adjusted to 5 µg/ml, and AO was added to a concentration of 5 µM prior to measurements. Increase and decrease in AO fluorescence indicate alkalization and acidification of the cell, respectively.

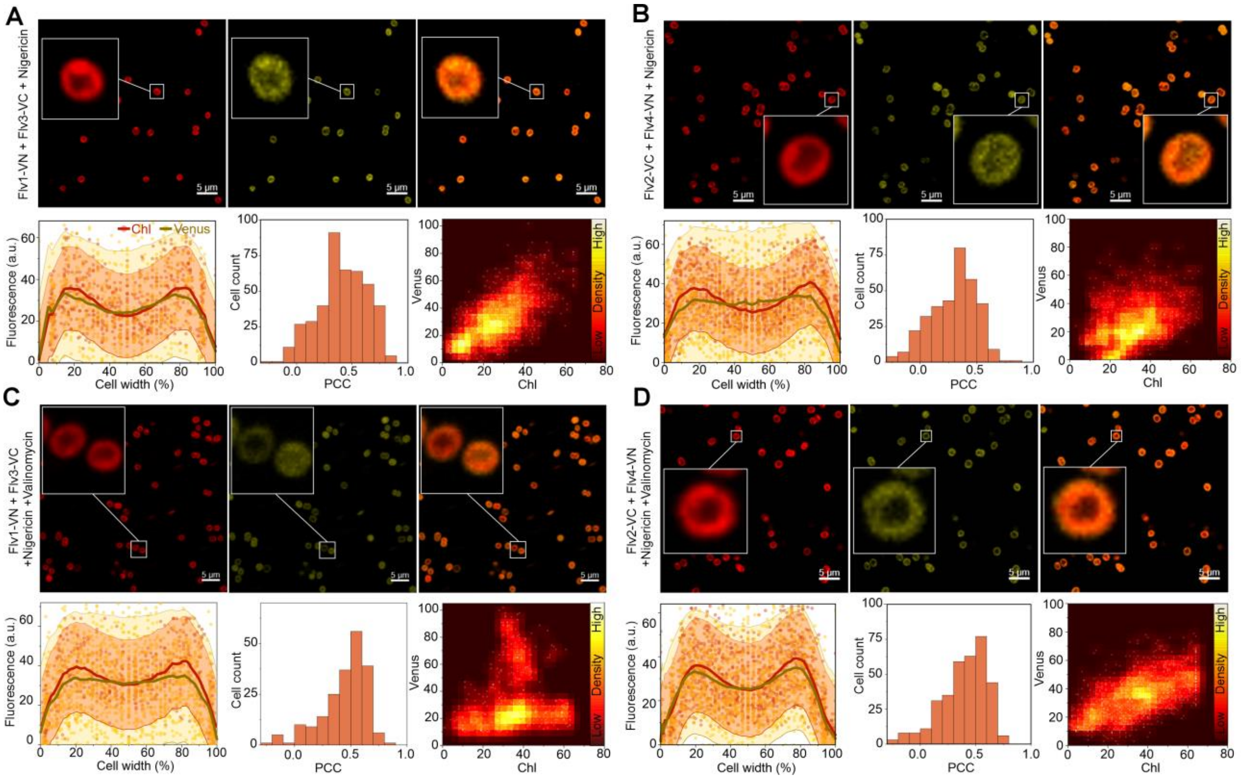


**Figure S14. Effect of Nigericin on the subcellular localisation of Flv1/3 and Flv2/4 interactions**. Supports Figure 5.

(A) BiFC tests between Flv1-VN and Flv3-VC with the culture supplemented with 20 µM Nigericin. (B) BiFC tests between Flv2-VC and Flv4-VN with 20 µM Nigericin. (C) BiFC tests between Flv1-VN and Flv3-VC with 20 µM Nigericin and 40 µM Valinomycin. (D) BiFC tests between Flv2-VC and Flv4-VN with 20 µM Nigericin and 40 µM Valinomycin. Upper panels show representative confocal micrographs from the chlorophyll and Venus fluorescence channels as well as a merged image. The lower panels show co-localisation of Chl (red) and Venus (yellow) fluorescence intensities over cross-sections of 300–350 cells (with width normalised to 100%) shown as a scatterplot. Red and gold lines show moving regression curves produced by locally weighted scatterplot smoothing (LOWESS). 95% confidence intervals are shown as the shadowed areas. The histograms show the distribution of the Pearson Correlation Coefficient (PCC) for Chl and Venus fluorescence co-localisation in all cells from three micrographs from individual replicates. The cytofluorograms show co-localised Chl and Venus fluorescence intensities as a scatter plot from all cells in four micrographs from independent replicates.

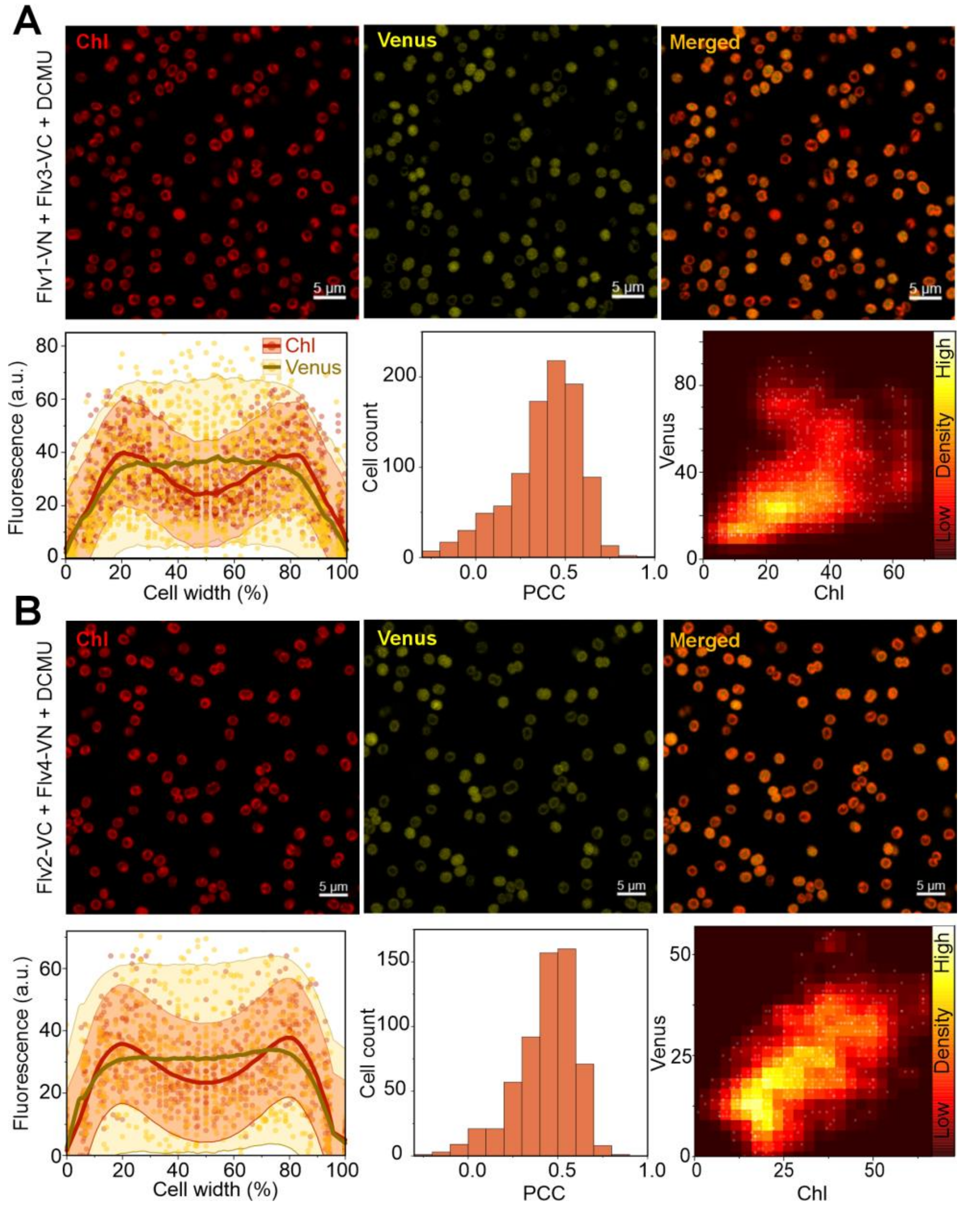


**Figure S15. Effect of DCMU on the subcellular localisation of Flv1/3 and Flv2/4 interactions**. Supports Figure 5.

(A) BiFC tests between Flv1:VN and Flv3:VC with the culture supplemented with 20 µM DCMU. (B) BiFC tests between Flv2:VC and Flv4:VN with 20 µM DCMU. For details, see legend for Fig. S8.

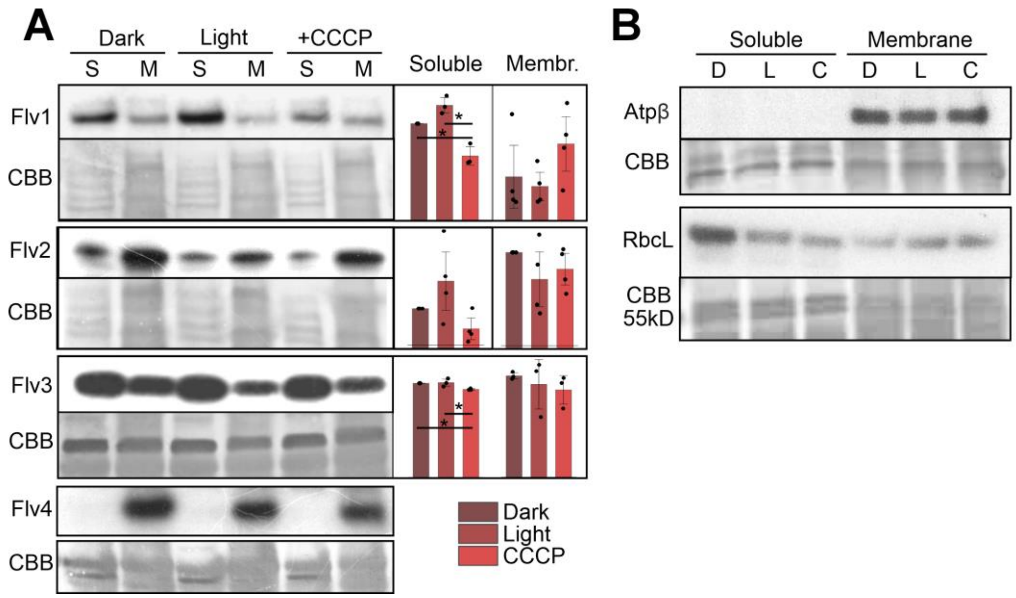


**Figure S16. Immunodetection of the subcellular localization of FDPs.** Supports Figure 5.

(A) Immunoblots showing the protein content of Flv1–4 in the soluble (S) and membrane (M) fractions of WT *Synechocystis* protein extracts. Prior to extraction, cells were either dark-adapted for 20 min, kept illuminated in growth conditions, or kept illuminated and then supplemented with 30 µM CCCP. Coomassie brilliant blue (CBB) -stained membranes are shown as loading controls. Panels on the right show quantifications of band intensities from 3-4 replicates ±SEM, normalised to the intensity of the dark-adapted soluble sample, or for the Flv2 membrane fraction blots, for the dark-adapted membrane sample (due to large variation in overall band intensities between replicates). Individual replicates are shown as black circles. Two-sample T-tests were used to test the difference between light and +CCCP samples.

(B) Control experiments for the purity of the soluble and membrane protein fractions. Exclusive membrane-localisation of the Atpβ of the ATP-synthase in dark (D), light (L), and CCCP-treated (C) cultures indicates no contamination of the soluble fraction with membrane proteins. In contrast, some amount of the large Rubisco subunit (RbcL) was found in the membrane fraction samples.

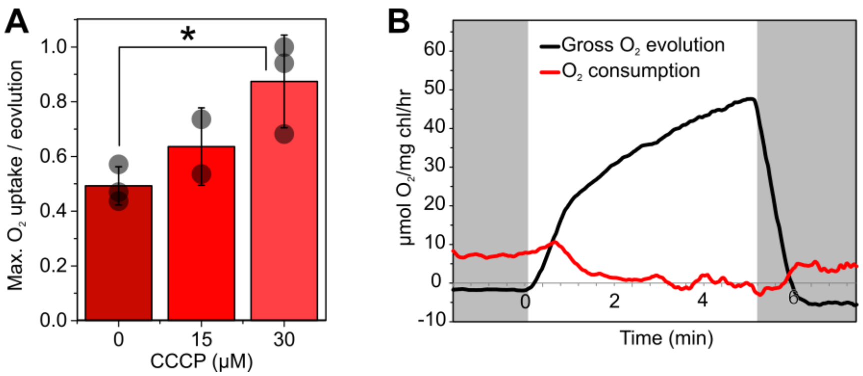


**Figure S17. Effect of CCCP on $O_2$ uptake.** Supports Figure 5.

(A) Maximum light-induced $O_2$ uptake rate during dark-to-light transitions, normalised to maximum gross $O_2$ evolution rate in BG-11 pH 7.5 supplemented with 0, 15, or 30 µM CCCP. WT *Synechocystis* cells were grown under 3% $CO_2$ for 3 days, harvested and resuspended in fresh BG-11 at Chl concentration of 10 µg/ml. Cells were illuminated with 500 µmol photons $m^{-2}s^{-1}$ for 5 min while $^{16}O_2$ and $^{18}O_2$ fluxes were monitored by MIMS. Light-induced uptake was calculated as total uptake - average uptake rate during 1 min in darkness. The values shown are averages from 3 (0 and 30 µM) or 2 (15µM) biological replicates +/- standard deviation, with individual data points shown as circles and statistically significant differences according to one-way ANOVA and Tukey's post-hoc test ($P<0.05$) as *.
(B) $O_2$ fluxes in *ΔFlv3* cells in the presence of 30mM CCCP. $O_2$ evolution and uptake were measured with MIMS as in (A).

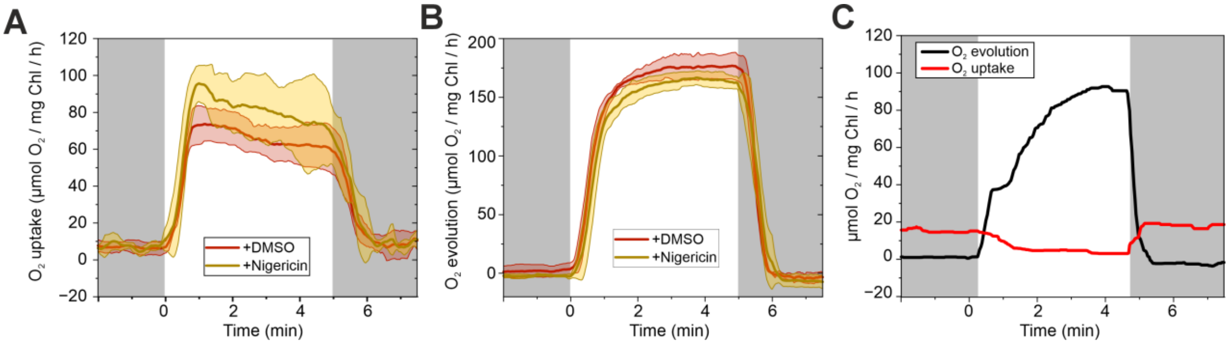


**Figure S18. Effect of Nigericin on $O_2$ uptake.** Supports Figure 6.

(A) $O_2$ uptake and (B) gross $O_2$ evolution in WT *Synechocystis* in the presence of 20 µM Nigericin or 4 µl of the DMSO solvent as measured by MIMS. This is unnormalised data from Figure 6. WT cells grown in 3% [$CO_2$] conditions under 50 µmol photons $m^{-2}$ $s^{-1}$ at pH 7.5 were measured by MIMS during 5 min of illumination at 500 µmol photons $m^{-2}$ $s^{-1}$. The traces are shown as $O_2$ uptake rate / maximal $O_2$ gross evolution rate, and are averaged from four biological replicates ± SD.
(C) Gross $O_2$ evolution (black) and $O_2$ uptake (red) in the ΔFlv3 mutant in the presence of 20 µM Nigericin.

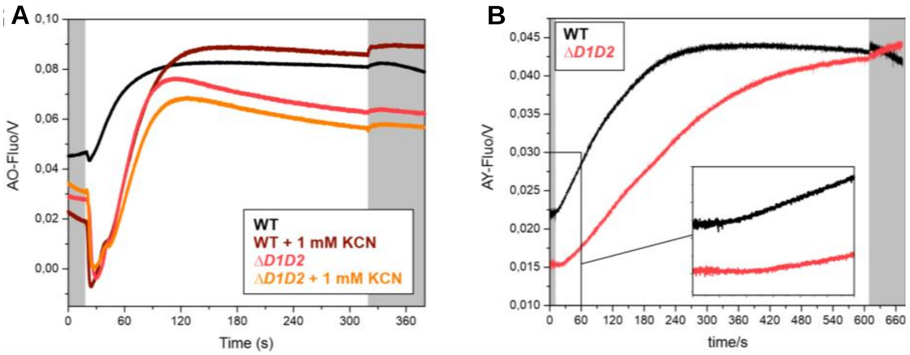


**Figure S19. Cytosolic alkalisation upon illumination.** Supports Figures 5, 6 and 8.

(A) Acridine orange (AO) and (B) acridine yellow (AY) fluorescence was recorded from dark-adapted WT and ΔD1D2 *Synechocystis* cultures with and without 1mM KCN under 216 µmol photons $m^{-2}s^{-1}$ of actinic light for 5 min (A) and 10 min (B). Cultures were grown in BG-11 pH 7.5 in air-level $CO_2$ under 50 µmol photons $m^{-2}s^{-1}$ for 4 days before measurements. Chl concentration was adjusted to 5 µg/ml, and AO and AY were added to a concentration of 5 µM prior to measurements. AY fluorescence derives from cytosolic pH changes only, while AO fluorescence represents an aggregate of cytosolic and lumenal pH changes. Representative traces from 3 (WT +AY), 2 (WT +AO, WT +AO+KCN, ΔD1D2 +AO, ΔD1D2 +AY), or 1 (ΔD1D2 +AO+KCN) biological replicate are shown.

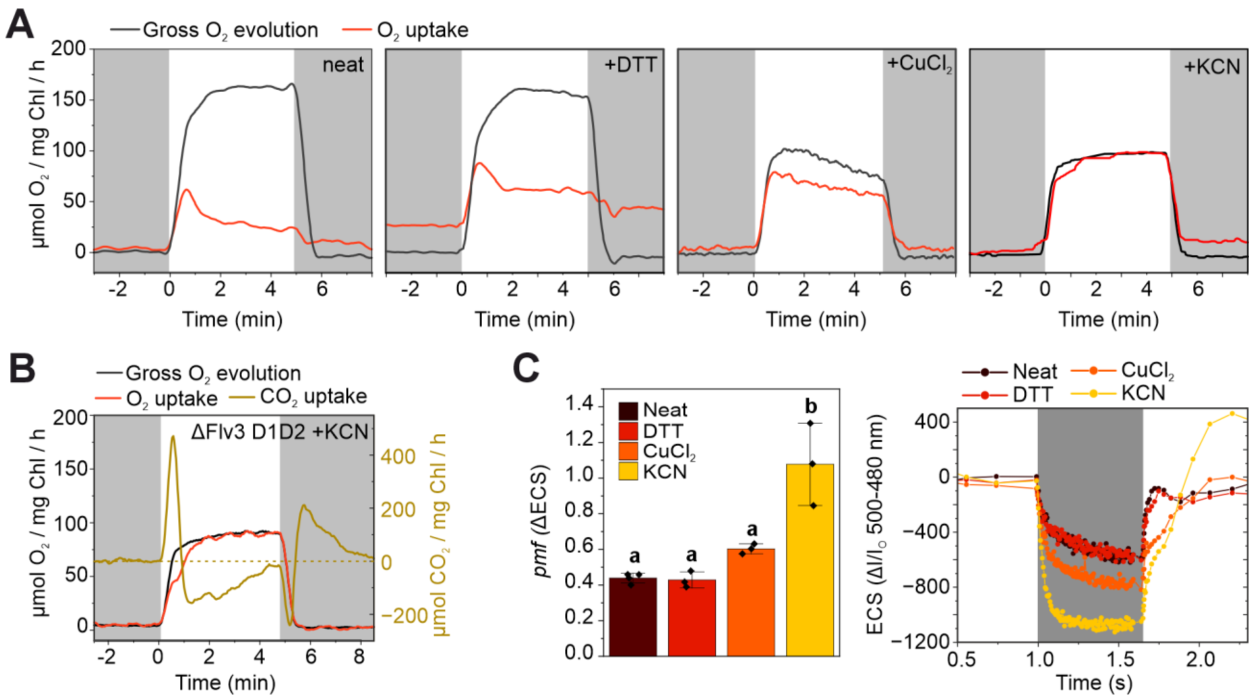


**Figure S20. Effect of DTT, $CuCl_2$, and KCN on photosynthesis.** Supports Figure 5.

(A) Gross $O_2$ evolution and $O_2$ uptake in WT *Synechocystis* in the presence or absence of 2 mM DTT, 100 µM $CuCl_2$, or 2 mM KCN, as measured by MIMS. The cultures were supplemented with the inhibitors, incubated in darkness for 10 min and then illuminated for 5 min at 500 µmol photons $m^{-2}$ $s^{-1}$. Cells were grown under air-level [$CO_2$] and 50 µmol photons $m^{-2}$ $s^{-1}$.
(B) The MIMS experiment with ΔFlv3 D1D2 cells in the presence of 2 mM KCN.
(C) Effect of 2 mM DTT, 100 µM $CuCl_2$, and 2 mM KCN on the magnitude of the *pmf* as measured from the light-induced change in the 500-480 nm ECS signal. Prior to measurement, cells were pre-illuminated for 2 min with 500 µmol µmol photons $m^{-2}$ $s^{-1}$ and then subjected 5 dark intervals of 600 ms with 5 s intervals. Averaged traces from the 5 dark-interval relaxation kinetics (DIRK) were used to determine the steady state *pmf* values. WT *Synechocystis* cells were grown under air-level [$CO_2$] and 50 µmol photons $m^{-2}$ $s^{-1}$, harvested and adjusted to 7.5 µg Chl / ml in fresh BG-11 medium. Means ±SD with individual replicates shown as dots are shown. Statistical significance of differences was tested by one-way ANOVA and Tukey's test ($P<0.05$).

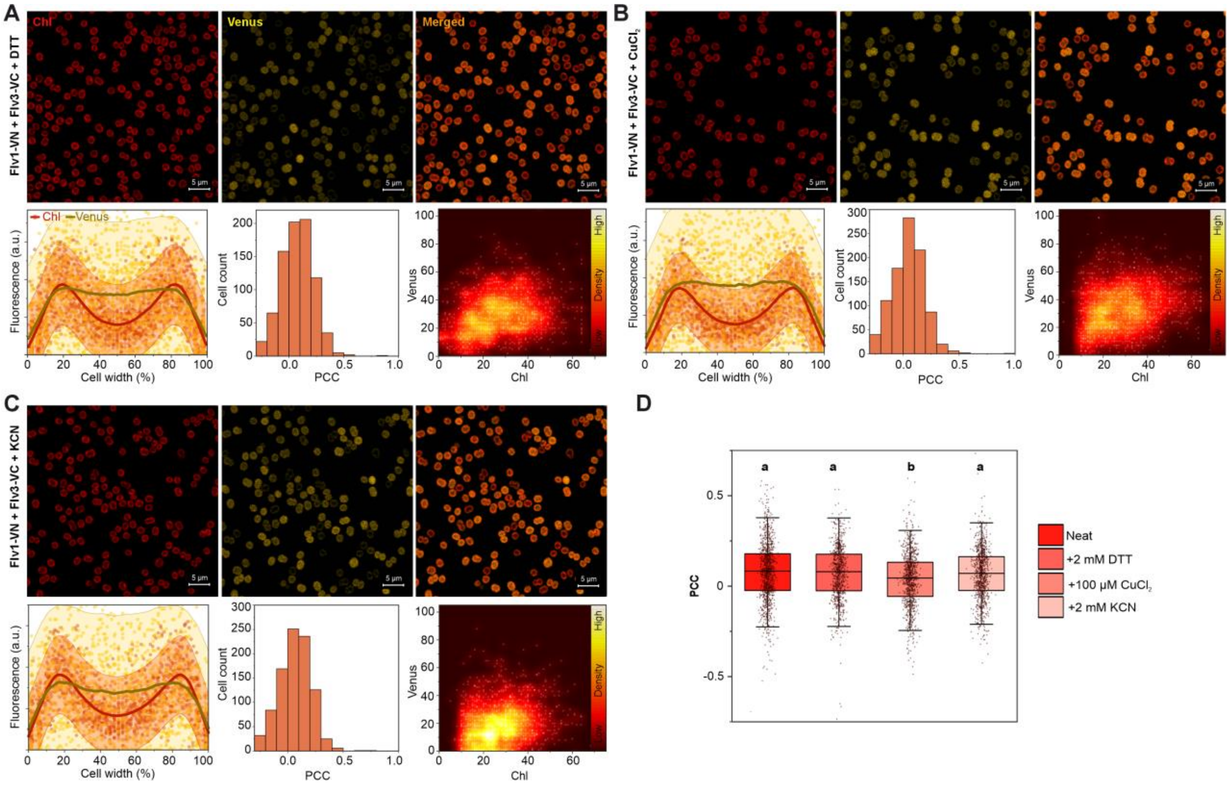


**Figure S21. Effect of DTT, $CuCl_2$, and KCN on the subcellular localisation of Flv1/3 interactions**. Supports Figure 5.

BiFC experiments with cells co-expressing Flv1-VN and Flv3-VC in the presence of (A) 2 mM DTT, (B) 100 µM $CuCl_2$, and (C) 2 mM KCN. For explanations of sub-panels see the legend for Figure 1. Cell cross section fluorescence profiles were analysed from 482 (A), 572 (B), and 338 (C) cells. Every 12$^{th}$ datapoint is shown in the scatterplots. (D) Comparison of PCC values for Chl and Venus fluorescence co-localisation between un-treated (neat) Flv1-VN+Flv3-VC cultures and cultures treated with 2 mM DTT, 100 µM $CuCl_2$, and 2 mM KCN. Box plots with median ± SD and individual data points as dots are shown. Statistical significance is indicated by lower case letters according to one-way ANOVA and Tukey's post-hoc tests for comparisons of means ($P<0.05$).

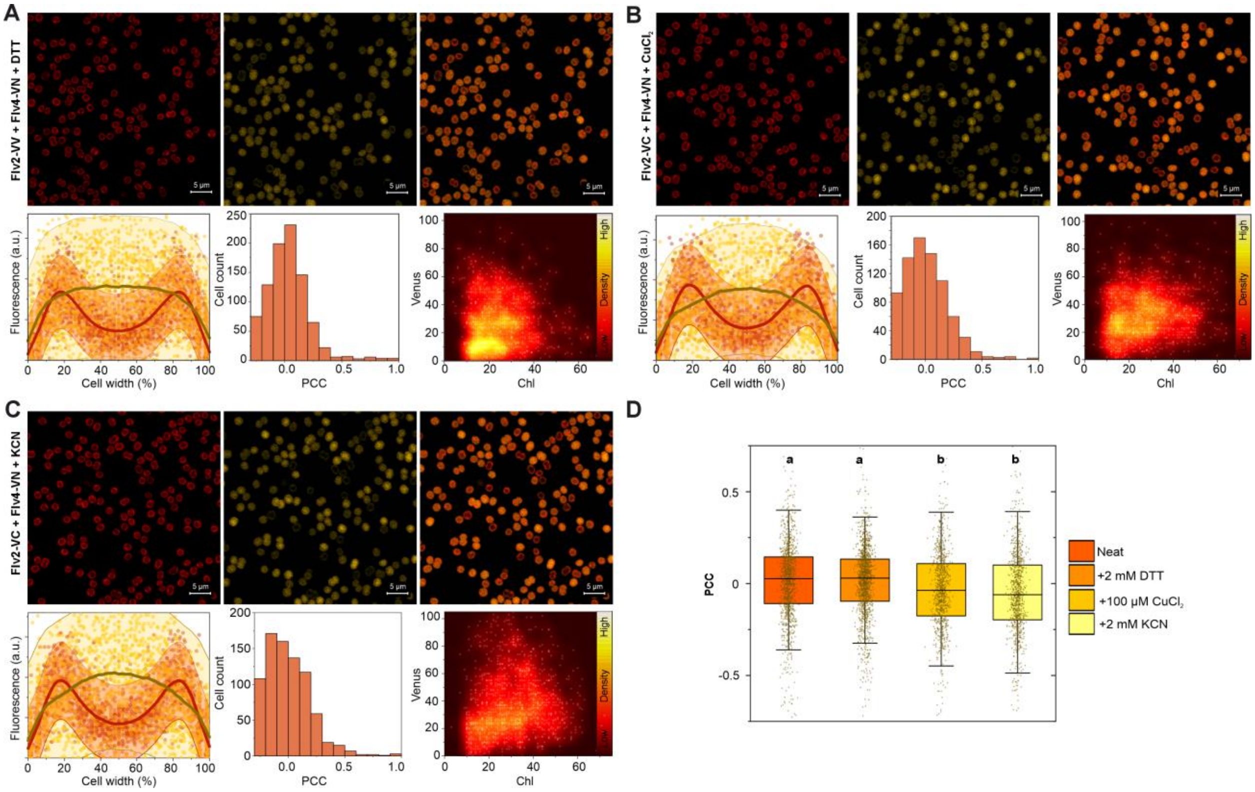


**Figure S22. Effect of DTT, $CuCl_2$, and KCN on the subcellular localisation of Flv2/4 interactions**. Supports Figure 5.

BiFC experiments with cells co-expressing Flv2-VC and Flv4-VN in the presence of (A) 2 mM DTT, (B) 100 µM $CuCl_2$, and (C) 2 mM KCN. For explanations of sub-panels see the legend for Figure 1. Cell cross section fluorescence profiles were analysed from 480 (A), 351 (B), and 376 (C) cells. Every 12$^{th}$ datapoint is shown in the scatterplots. (D) Comparison of PCC values for Chl and Venus fluorescence co-localisation between un-treated (neat) Flv1-VN+Flv3-VC cultures and cultures treated with 2 mM DTT, 100 µM $CuCl_2$, and 2 mM KCN. Box plots with median ± SD and individual data points as dots are shown. Statistical significance is indicated by lower case letters according to one-way ANOVA and Tukey's post-hoc tests for comparisons of means ($P<0.05$).

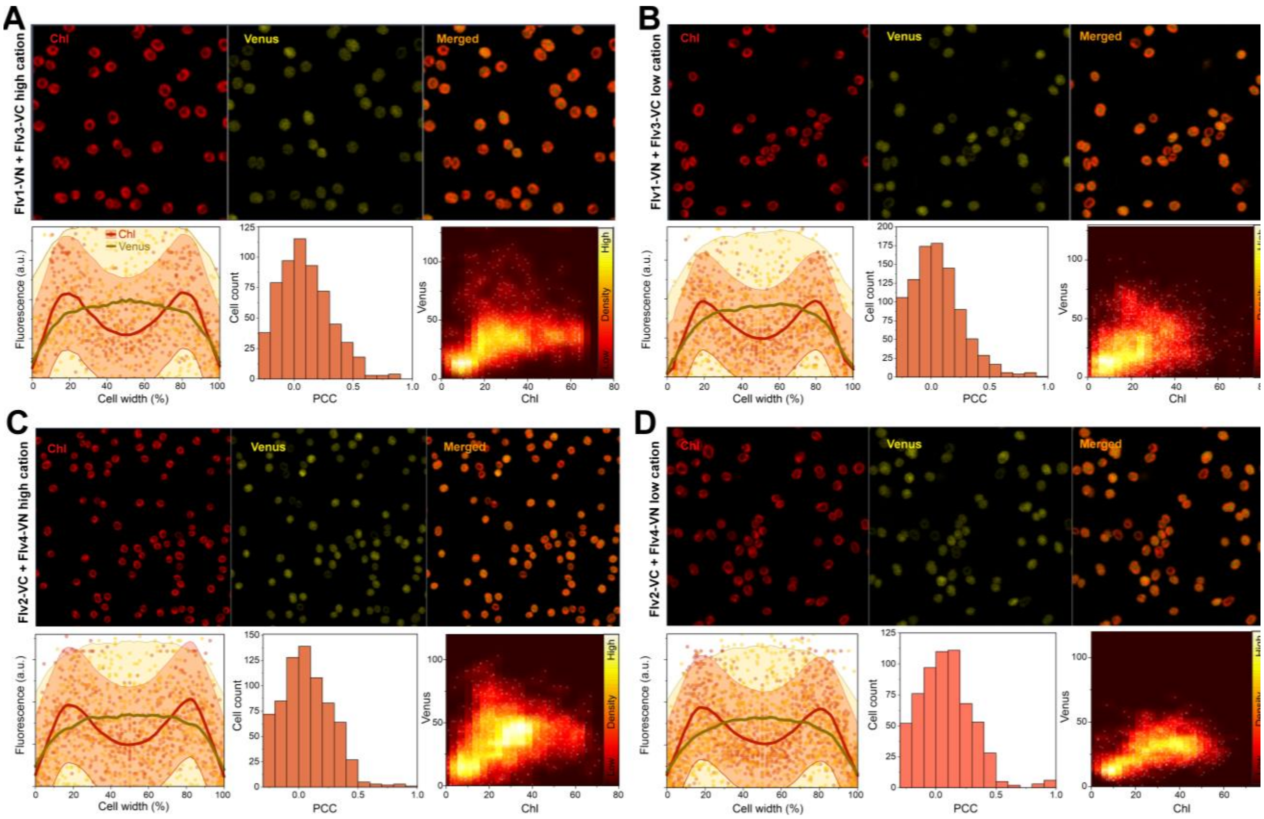


**Figure S23. Effect of cation concentration on the subcellular localisation of Flv1/3 and Flv2/4 interactions**. Supports Figure 5.

BiFC tests between Flv1-VN and Flv3-VC and between Flv2-VC and Flv4-VN in BG-11 medium with no added source of $Mg^{2+}$ or $Ca^{2+}$ (A and C, respectively) or in BG-11 supplemented with 25mM $MgCl_2$ and 30mM $CaCl_2$ (B and D, respectively) are shown. Upper panels show representative confocal micrographs from the chlorophyll and Venus fluorescence channels as well as a merged image. The lower panels show co-localisation of Chl (red) and Venus (yellow) fluorescence intensities over cross-sections of 300-400 cells (with width normalised to 100%) shown as a scatterplot. Every 12$^{th}$ value is shown in the scatterplots. Brighter red and yellow lines show moving regression curves produced by locally weighted scatterplot smoothing (LOWESS). The histograms show the distribution of the Pearson Correlation Coefficient (PCC) for Chl and Venus fluorescence co-localisation in all cells from three micrographs from individual replicates. A PCC of 1 indicates perfect co-localisation and a PCC of 0 no co-localisation. The cytofluorograms show co-localised Chl and Venus fluorescence intensities as a scatter plot from all cells in three micrographs from independent replicates.

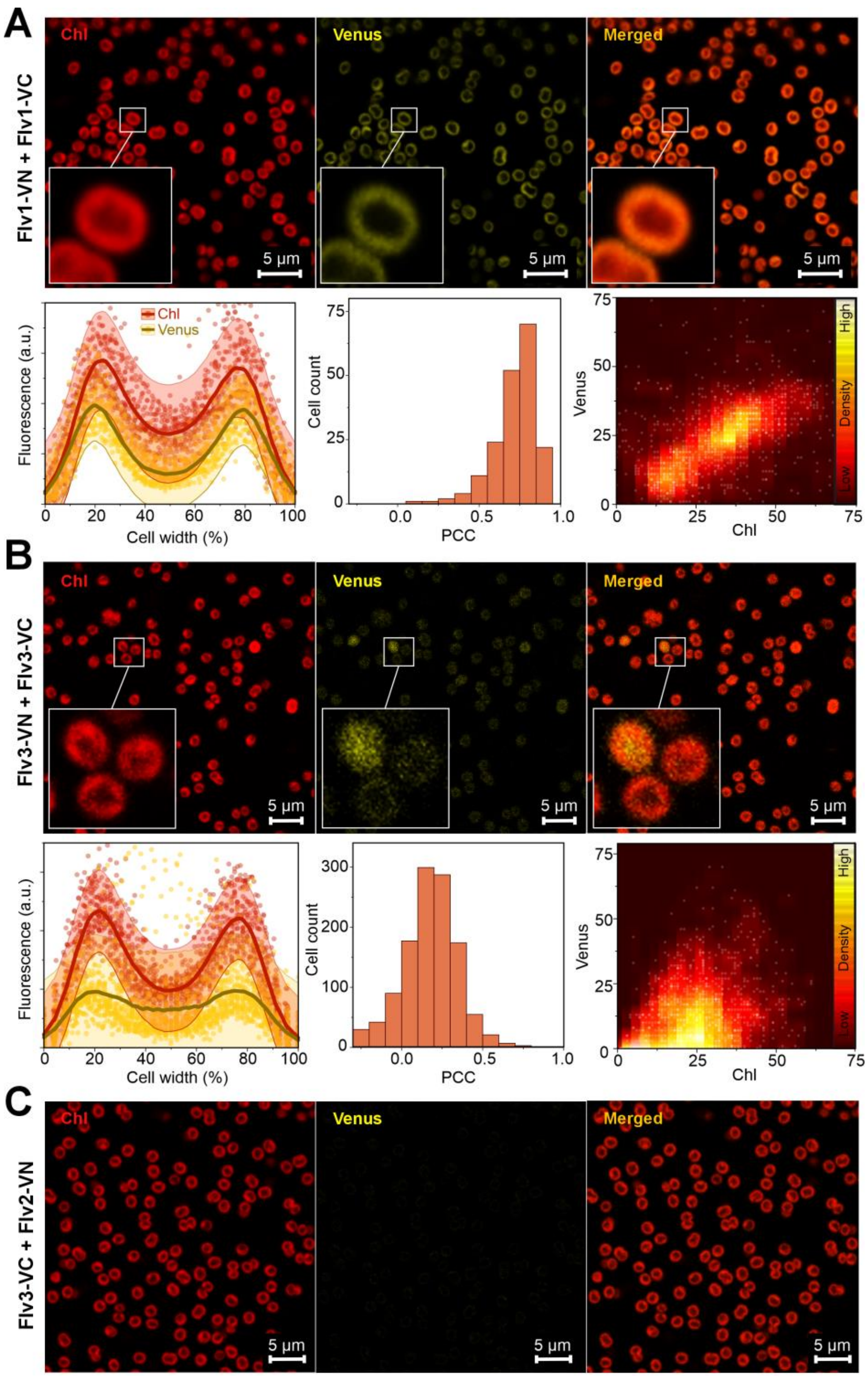


**Figure S24. BiFC tests for Flv1 (A) and Flv3 self-interactions (B) and between Flv2 and Flv3 (C).** Supports Figures 4 and 5.

For details see legends for Figures 4 and 5.

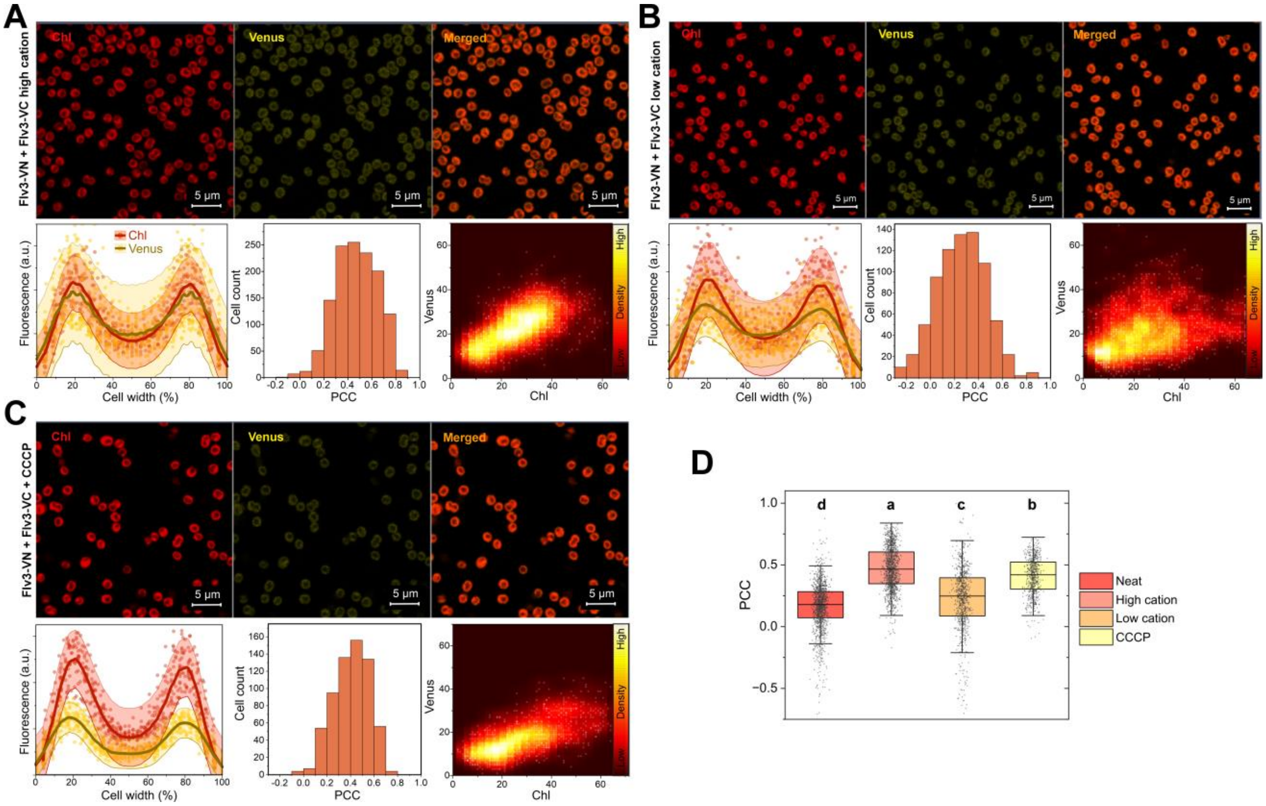


**Figure S25. Effect of cation concentration and *pmf* uncoupling on the subcellular localisation of Flv3 self-interactions**. Supports Figure 5.

See legend for Figure S8 for explanations of subpanels. BiFC tests were performed in medium supplemented with 25 mM $MgCl_2$ and 30mM $CaCl_2$ (high cation, A), in medium without added source of $MgCl_2$ and $CaCl_2$ (low cation, B), and in medium supplemented with 30 µM CCCP (C).

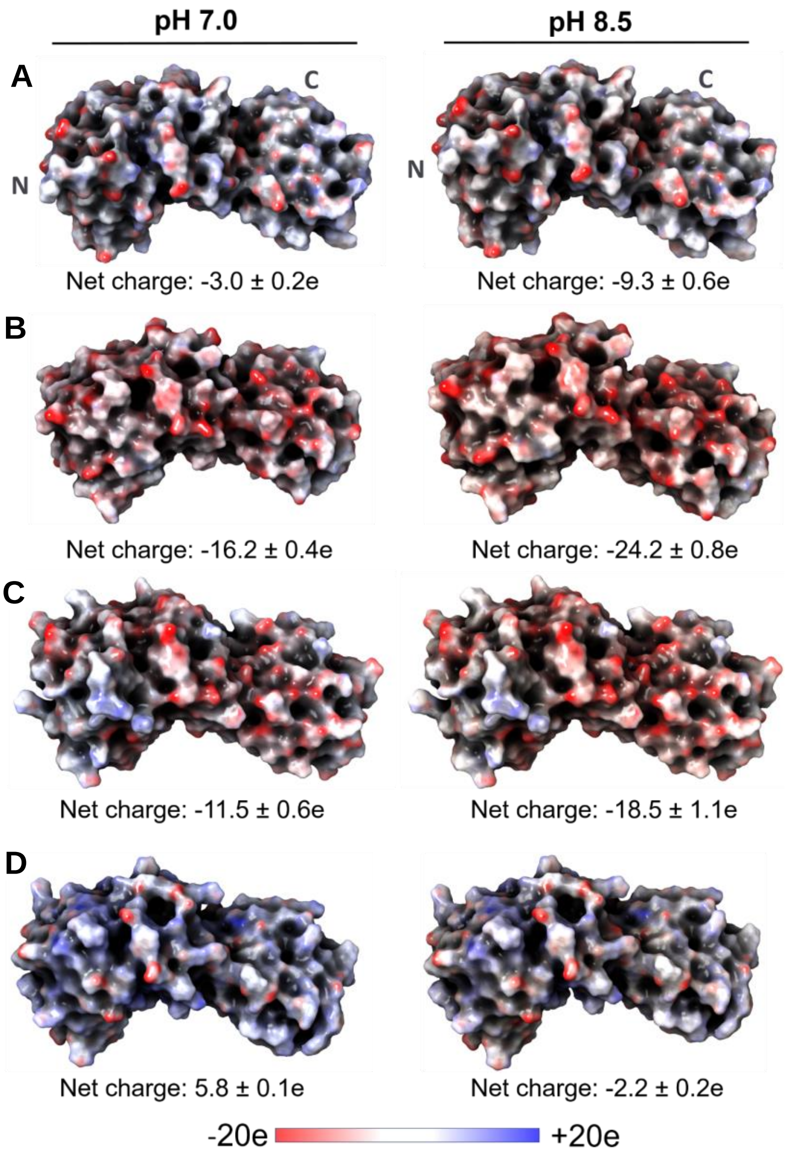


**Figure S26. *In silico* analysis of electrostatic surface charges of FDP monomers under pH 7.0 and pH 8.5.** Supports Figure 6.

(A) Flv1 monomer, (B) Flv2 monomer, (C) Flv3 monomer, and (D) Flv4 monomer. Surface charges are color-coded with red representing negatively charged regions, blue depicting positively charged regions, and white showing neutral regions. Left panels show charge at pH 7.0, while right panels depict charge at pH 8.5. Since relative orientation of C-terminal Flavin reductase domain could not be modelled reliably, only core domains (β-lactamase-like and Flavodoxin-like domains) are shown for visualisation purposes.

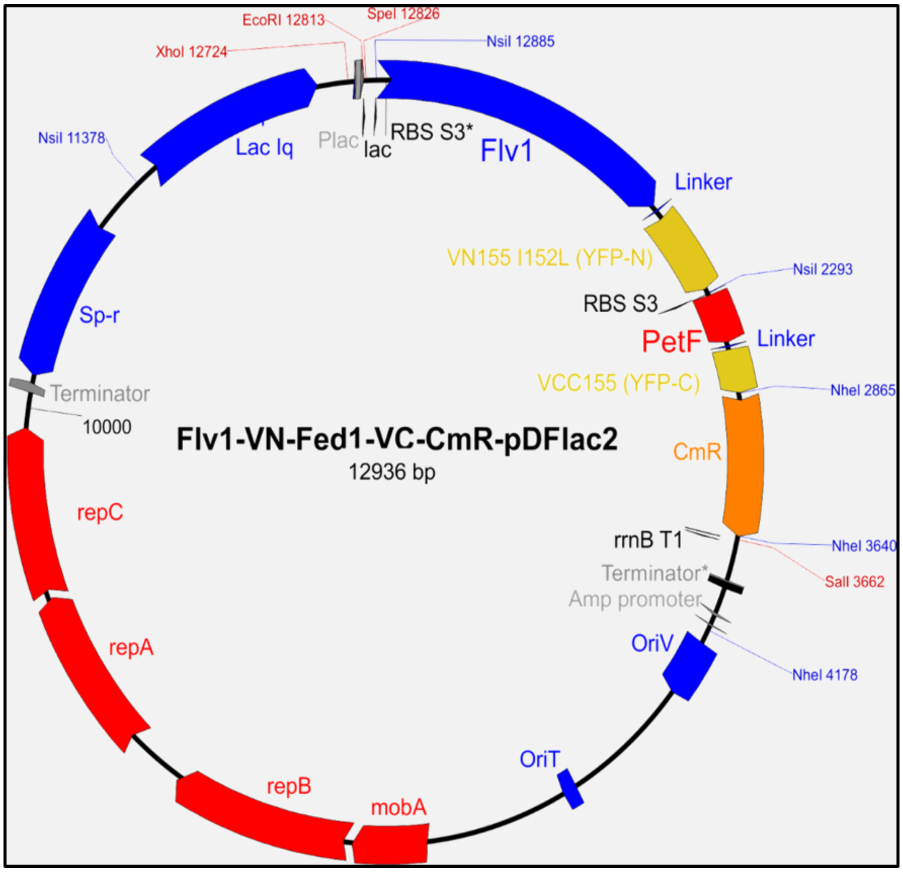


**Figure S27. Plasmid map of a BiFC construct to express *Synechocystis flv1* and *petF* (Fed1) as fusion proteins with N- and C-terminal Venus fragments, respectively.**
Supports Figures 2, 4, and 5.

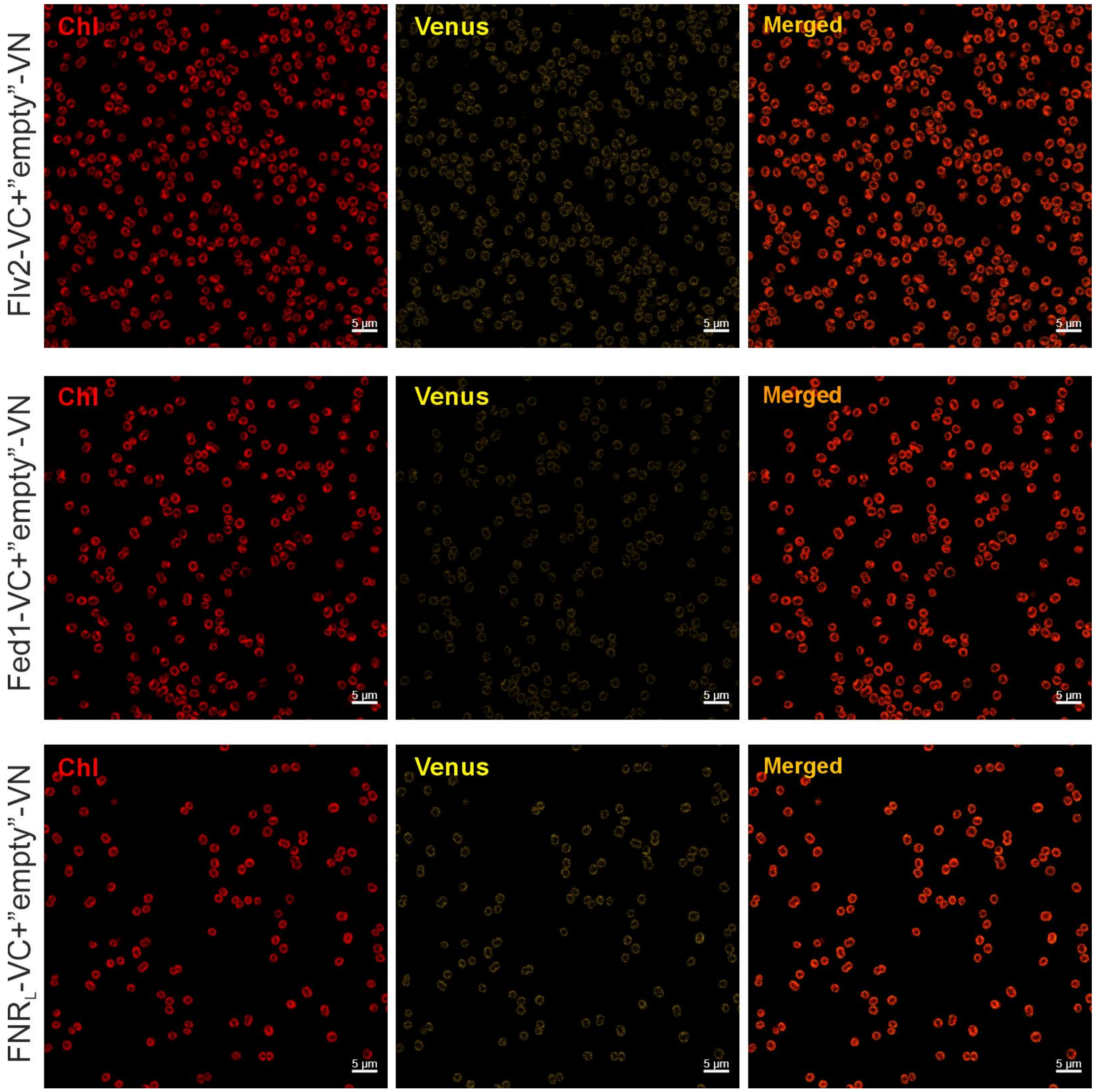


**Figure S28. "Empty" vector control experiments for BiFC.** Supports Figures 2, 4, and 5.

BiFC experiments were performed as for the other figures with strains expressing a protein of interest (Flv2, Fed1, or FNR$_L$) fused to the C-terminal Venus fragment together with a non-fused N-terminal Venus fragment. Representative confocal micrographs are shown.

**Table S1. Primers used for generating the ∆Fed10 and ∆Fed11 strains.**

| Primer Name | Sequence |
|---|---|
| ∆FED10-GIB1 | GACTACTATAGGGCGAATTGGGTACGTAAAACCGATTACATTGTGGTGGGCAA |
| ∆FED10-GIB2 | TTCTGGCTGGATGATGGGGCGATGGAAATGTGCGCGAAAACTGAATAATCG |
| ∆FED10-GIB3 | CGATTATTCAGTTTTCGCGCACATTTCCATCGCCCCATCATCCAGCCAGAA |
| ∆FED10-GIB4 | CCGACTCGTTTCTATAGCGGTTTTGATAATCCACGTTGTGTCTCAAAATCTCTGATGTTAC |
| ∆FED10-GIB5 | GTAACATCAGAGATTTTGAGACACAACGTGGATTATCAAAACCGCTATAGAAACGAGTCGG |
| ∆FED10-GIB6 | ACTAAAGGGAACAAAAGCTGGAGCTCGGCAATATCCTCGTTGCGTTTAATAACTA |
| ∆FED11-GIB1 | GACTACTATAGGGCGAATTGGGTACGTAAAAAGCCCGGCAGAGGGGAT |
| ∆FED11-GIB2 | TTCTGGCTGGATGATGGGGCGATTGCAATTTCCTCCTGTCCTCAGGG |
| ∆FED11-GIB3 | CCCTGAGGACAGGAGGAAATTGCAATCGCCCCATCATCCAGCCAGAA |
| ∆FED11-GIB4 | TTGGCCTCCGGTGATCGCCCCCACGTTGTGTCTCAAAATCTCTGATGTTAC |
| ∆FED11-GIB5 | GTAACATCAGAGATTTTGAGACACAACGTGGGGGCGATCACCGGAGGCCAA |
| ∆FED11-GIB6 | ACTAAAGGGAACAAAAGCTGGAGCTCGGCGGATAACGATGTTGTACTTTATAGT |

**Table S2. Results from statistical tests in the main article. Supports Figures 1, 3, and 5.**

| **Figure 1B** | One-way ANOVA | | | | |
|---|---|---|---|---|---|
| strain | n | Mean | SD | SEM | Tukey's groups |
| WT | 3 | 0.054 | 0.013 | 0.008 | a |
| ΔFlv3 | 4 | 0.052 | 0.020 | 0.010 | a |
| $\Delta FNR_L$ | 3 | 0.004 | 0.004 | 0.002 | b |
| $\Delta FNR_S$ | 3 | 0.052 | 0.010 | 0.006 | a |
| | DF | Sum of Sq | Mean Sq | F Value | Prob>F |
| Model | 3 | 0.006 | 0.002 | 9.117 | **0.004** |
| Error | 9 | 0.002 | 0.000 | | |
| Total | 12 | 0.007 | | | |
| **Figure 1D** | **One-way ANOVA and Tukey's test** | | | | |
| strain | n | Mean | SD | SEM | Tukey's groups |
| WT air | 5 | 0.418 | 0.022 | 0.010 | a |
| ΔFlv3 air | 3 | 0.059 | 0.023 | 0.013 | c |
| $\Delta FNR_L$ air | 7 | 0.338 | 0.052 | 0.019 | b |
| $\Delta FNR_S$ air | 7 | 0.449 | 0.028 | 0.011 | a |
| | DF | Sum of Sq | Mean Sq | F Value | Prob>F |
| Model | 3 | 0.346 | 0.115 | 87.638 | **<0.0001** |
| Error | 18 | 0.024 | 0.001 | | |
| Total | 21 | 0.370 | | | |
| **Figure 1D** | **One-way ANOVA and Tukey's test** | | | | |
| strain | n | Mean | SD | SEM | Tukey's groups |
| WT 3%$CO_2$ | 5 | 0.413 | 0.080 | 0.036 | a |
| ΔFlv3 3% $CO_2$ | 3 | 0.053 | 0.046 | 0.027 | b |
| $\Delta FNR_L$ 3%$CO_2$ | 6 | 0.427 | 0.053 | 0.022 | a |
| $\Delta FNR_S$ 3%$CO_2$ | 3 | 0.484 | 0.139 | 0.080 | a |
| | DF | Sum of Sq | Mean Sq | F Value | Prob>F |
| Model | 3 | 0.369 | 0.123 | 19.437 | <0.0001 |
| Error | 13 | 0.082 | 0.006 | | |
| Total | 16 | 0.451 | | | |
| **Figure 1E** | **One-sample T-tests** | | | null hypothesis: mean=1 | |
| **Flv1** | n | Mean | SD | SEM | P |
| $\Delta FNR_L$ | 4 | 1.104 | 0.104 | 0.052 | **0.137** |
| $\Delta FNR_S$ | 4 | 1.168 | 0.047 | 0.024 | **0,006*** |
| **Flv2** | n | Mean | SD | SEM | P |
| $\Delta FNR_L$ | 4 | 0.515 | 0.169 | 0.084 | **0,0105*** |
| $\Delta FNR_S$ | 4 | 1.189 | 0.288 | 0.144 | **0.280** |
| **Flv3** | n | Mean | SD | SEM | P |
| $\Delta FNR_L$ | 4 | 1.292 | 0.235 | 0.118 | **0.089** |
| $\Delta FNR_S$ | 4 | 1.421 | 0.340 | 0.170 | **0.089** |
| **Flv4** | n | Mean | SD | SEM | P |
| $\Delta FNR_L$ | 3 | 0.837 | 0.274 | 0.158 | **0.410** |
| $\Delta FNR_S$ | 3 | 0.879 | 0.083 | 0.048 | **0.127** |
| **Figure 1F** | **One-sample T-tests** | | | null hypothesis: mean=1 | |
| **$FNR_L$** | n | Mean | SD | SEM | P |
| ΔFlv3 | 4 | 1.055 | 0.130 | 0.065 | **0.459** |

| **Figure 3B** | **One-way ANOVA and Tukey's test** | | | | |
|---|---|---|---|---|---|
| | n | Mean | SD | SEM | Tukey's groups |
| WT | 8 | 0.441 | 0.066 | 0.023 | ab |
| ΔFed3 | 4 | 0.466 | 0.048 | 0.024 | ab |
| ΔFed4 | 4 | 0.514 | 0.076 | 0.038 | a |
| ΔFed6 | 4 | 0.397 | 0.048 | 0.024 | ab |
| ΔFed789 | 4 | 0.340 | 0.036 | 0.018 | b |
| ΔFed9 | 4 | 0.339 | 0.083 | 0.041 | b |
| ΔFed10 | 4 | 0.421 | 0.028 | 0.014 | ab |
| ΔFed11 | 4 | 0.399 | 0.032 | 0.016 | ab |
| | DF | Sum of Sq | Mean Sq | F Value | Prob>F |
| Model | 7 | 0.103 | 0.015 | 4.509 | **0.002** |
| Error | 28 | 0.091 | 0.003 | | |
| Total | 35 | 0.194 | | | |
| **Figure 5E** | **One-way ANOVA and Tukey's test** | | | | |
| | n | Mean | SD | SEM | Tukey's groups |
| Neat | 350 | 0.093 | 0.227 | 0.012 | c |
| CCCP | 2989 | 0.368 | 0.181 | 0.003 | b |
| Nigericin | 430 | 0.418 | 0.224 | 0.011 | a |
| Nig.+ Val. | 206 | 0.436 | 0.225 | 0.016 | a |
| DCMU | 946 | 0.380 | 0.211 | 0.007 | b |
| | DF | Sum of Sq | Mean Sq | F Value | Prob>F |
| Model | 1 | 21.055 | 21.055 | 455.65 | **<0.0001** |
| Error | 1294 | 59.794 | 0.046 | | |
| Total | 1295 | 80.849 | | | |
| **Figure 5F** | **One-way ANOVA and Tukey's test** | | | | |
| | n | Mean | SD | SEM | Tukey's groups |
| Neat | 355 | 0.057 | 0.215 | 0.011 | d |
| CCCP | 893 | 0.291 | 0.179 | 0.006 | c |
| Nigericin | 339 | 0.288 | 0.207 | 0.011 | c |
| Nig.+ Val. | 345 | 0.394 | 0.207 | 0.011 | b |
| DCMU | 601 | 0.433 | 0.168 | 0.007 | a |
| | DF | Sum of Sq | Mean Sq | F Value | Prob>F |
| Model | 4 | 34.642 | 8.660 | 240.088 | **<0.0001** |
| Error | 2528 | 91.190 | 0.036 | | |
| Total | 2532 | 125.832 | | | |

### Table S3. Results from statistical tests in Supplementary Material

| Figure S1A | One-way ANOVA | | | | |
|---|---|---|---|---|---|
| strain | n | Mean | SD | SEM | Tukey's |
| WT air | 5 | 77.89 | 19.34 | 8.65 | a |
| ΔFlv3 air | 3 | 15.99 | 4.67 | 2.70 | b |
| $\Delta FNR_L$ air | 7 | 62.32 | 8.73 | 3.30 | a |
| $\Delta FNR_S$ air | 7 | 72.31 | 21.54 | 8.14 | a |
| | DF | Sum of Sq | Mean Sq | F Value | Prob>F |
| Model | 3 | 8347.9 | 2782.6 | 10.477 | **0.0003** |
| Error | 18 | 4780.7 | 265.6 | | |
| Total | 21 | 13128.6 | | | |

| Figure S1A | One-way ANOVA | | | | |
|---|---|---|---|---|---|
| strain | n | Mean | SD | SEM | Tukey's |
| WT 3%$CO_2$ | 5 | 77.44 | 10.21 | 4.56 | a |
| ΔFlv3 3%$CO_2$ | 3 | 10.12 | 4.94 | 2.85 | b |
| $\Delta FNR_L$ 3%$CO_2$ | 6 | 83.76 | 5.08 | 2.07 | a |
| $\Delta FNR_S$ 3%$CO_2$ | 3 | 90.71 | 18.21 | 10.52 | a |
| | DF | Sum of Sq | Mean Sq | F Value | Prob>F |
| Model | 3 | 13457.3 | 4485.76 | 46.36 | **<0.0001** |
| Error | 13 | 1257.9 | 96.76 | | |
| Total | 16 | 14715.2 | | | |

| Figure S1B | One-way ANOVA | | | | |
|---|---|---|---|---|---|
| strain | n | Mean | SD | SEM | Tukey's |
| WT air | 5 | 180.15 | 52.41 | 23.44 | a |
| ΔFlv3 air | 3 | 159.47 | 17.34 | 10.01 | a |
| $\Delta FNR_L$ air | 7 | 149.30 | 10.60 | 4.01 | a |
| $\Delta FNR_S$ air | 7 | 148.32 | 44.55 | 16.84 | a |
| | DF | Sum of Sq | Mean Sq | F Value | Prob>F |
| Model | 3 | 3636.2 | 1212.079 | 0.903 | **0.459** |
| Error | 18 | 24172.5 | 1342.916 | | |
| Total | 21 | 27808.7 | | | |

| Figure S1B | One-way ANOVA | | | | |
|---|---|---|---|---|---|
| strain | n | Mean | SD | SEM | Tukey's |
| WT 3%$CO_2$ | 5 | 171.53 | 21.35 | 9.55 | a |
| ΔFlv3 3%$CO_2$ | 3 | 98.09 | 3.72 | 2.15 | b |
| $\Delta FNR_L$ 3%$CO_2$ | 6 | 176.76 | 19.79 | 8.08 | a |
| $\Delta FNR_S$ 3%$CO_2$ | 3 | 163.50 | 14.51 | 8.38 | a |
| | DF | Sum of Sq | Mean Sq | F Value | Prob>F |
| Model | 2 | 353.9 | 176.971 | 0.463 | **0.641** |
| Error | 11 | 4201.8 | 381.978 | | |
| Total | 13 | 4555.7 | | | |

| Figure S1C | One-way ANOVA | | | | |
|---|---|---|---|---|---|
| strain | n | Mean | SD | SEM | Tukey's |
| WT air | 5 | 2.69 | 4.46 | 2.00 | a |
| ΔFlv3 air | 3 | 6.86 | 1.88 | 1.09 | ab |
| $\Delta FNR_L$ air | 7 | 11.92 | 4.08 | 1.54 | a |
| $\Delta FNR_S$ air | 7 | 5.96 | 3.82 | 1.44 | b |
| | DF | Sum of Sq | Mean Sq | F Value | Prob>F |
| Model | 3 | 13868.2 | 4622.7 | 14.2 | **0.0002** |
| Error | 13 | 4229.5 | 325.3 | | |
| Total | 16 | 18097.7 | | | |

| Figure S1C | One-way ANOVA | | | | |
|---|---|---|---|---|---|
| strain | n | Mean | SD | SEM | Tukey's |
| WT 3%$CO_2$ | 5 | 5.87 | 2.92 | 1.30 | a |
| ΔFlv3 3%$CO_2$ | 3 | 4.39 | 3.13 | 1.80 | a |
| $\Delta FNR_L$ 3%$CO_2$ | 6 | 7.20 | 3.56 | 1.45 | a |
| $\Delta FNR_S$ 3%$CO_2$ | 3 | 7.91 | 2.55 | 1.47 | a |
| | DF | Sum of Sq | Mean Sq | F Value | Prob>F |
| Model | 3 | 24.2 | 8.071 | 0.807 | **0.512** |
| Error | 13 | 130.1 | 10.006 | | |
| Total | 16 | 154.3 | | | |

| Figure S1D | One-way ANOVA | | | | |
|---|---|---|---|---|---|
| strain | n | Mean | SD | SEM | Tukey's |
| WT air | 5 | 75.20 | 20.14 | 9.01 | a |
| ΔFlv3 air | 3 | 9.13 | 2.81 | 1.62 | b |
| $\Delta FNR_L$ air | 7 | 50.40 | 8.43 | 3.19 | a |
| $\Delta FNR_S$ air | 7 | 66.35 | 19.38 | 7.32 | a |
| | DF | Sum of Sq | Mean Sq | F Value | Prob>F |
| Model | 3 | 9397.1 | 3132.4 | 13.06 | **<0.0001** |
| Error | 18 | 4318.5 | 239.9 | | |
| Total | 21 | 13715.6 | | | |

| Figure S1D | One-way ANOVA | | | | |
|---|---|---|---|---|---|
| strain | n | Mean | SD | SEM | Tukey's |
| WT 3%$CO_2$ | 5 | 71.57 | 8.29 | 3.71 | a |
| ΔFlv3 3%$CO_2$ | 3 | 5.51 | 5.01 | 2.89 | b |
| $\Delta FNR_L$ 3%$CO_2$ | 6 | 76.56 | 4.78 | 1.95 | a |
| $\Delta FNR_S$ 3%$CO_2$ | 3 | 82.80 | 15.66 | 9.04 | a |
| | DF | Sum of Sq | Mean Sq | F Value | Prob>F |
| Model | 3 | 12556.2 | 4185.39 | 58.54 | **<0.0001** |
| Error | 13 | 929.5 | 71.50 | | |
| Total | 16 | 13485.7 | | | |

| Figure S1E | One-way ANOVA | | | | |
|---|---|---|---|---|---|
| strain | n | Mean | SD | SEM | Tukey's |
| WT air | 7 | 150.46 | 21.93 | 8.29 | a |
| ΔFlv3 air | 3 | 172.44 | 26.02 | 15.02 | a |
| $\Delta FNR_L$ air | 6 | 128.97 | 38.82 | 15.85 | a |
| $\Delta FNR_S$ air | 6 | 119.78 | 22.85 | 9.33 | a |
| | DF | Sum of Sq | Mean Sq | F Value | Prob>F |
| Model | 3 | 7091.8 | 2363.93 | 2.96 | **0.060** |
| Error | 18 | 14382.6 | 799.03 | | |
| Total | 21 | 21474.4 | | | |

| Figure S1E | One-way ANOVA | | | | |
|---|---|---|---|---|---|
| strain | n | Mean | SD | SEM | Tukey's |
| WT 3%$CO_2$ | 4 | 203.56 | 31.37 | 15.69 | a |
| ΔFlv3 3%$CO_2$ | 2 | 33.62 | 32.16 | 22.74 | b |
| $\Delta FNR_L$ 3%$CO_2$ | 3 | 199.01 | 45.78 | 26.43 | a |
| $\Delta FNR_S$ 3%$CO_2$ | 3 | 190.87 | 37.95 | 21.91 | a |
| | DF | Sum of Sq | Mean Sq | F Value | Prob>F |
| Model | 3 | 45526.8 | ####### | 10.98 | **0.003** |
| Error | 8 | 11059.7 | 1382.47 | | |
| Total | 11 | 56586.6 | | | |

| Figure S2B | One-way ANOVA | | | | |
|---|---|---|---|---|---|
| strain | n | Mean | SD | SEM | Tukey's |
| WT 3%$CO_2$ | 3 | 0.132 | 0.032 | 0.018 | a |
| $\Delta FNR_L$ 3%$CO_2$ | 3 | 0.057 | 0.012 | 0.007 | b |
| $\Delta FNR_S$ 3%$CO_2$ | 3 | 0.136 | 0.004 | 0.002 | a |
| | DF | Sum of Sq | Mean Sq | F Value | Prob>F |
| Model | 2 | 0.012 | 0.0059 | 15.00 | **0.005** |
| Error | 6 | 0.002 | 0.0004 | | |
| Total | 8 | 0.014 | | | |

| Figure S2D | Two-Sample T-test | | | | |
|---|---|---|---|---|---|
| null hypothesis: difference in mean=0 | | | | | |
| **low light** | n | Mean | SD | SEM | P |
| WT | 6 | 0.102 | 0.021 | 0.01 | |
| ΔFNRL | 6 | 0.046 | 0.012 | 0 | |
| difference | 6 | 0.055 | 0.026 | 0.01 | **0.0035** |

| Figure S9 | One-way ANOVA | | | | |
|---|---|---|---|---|---|
| strain | n | Mean | SD | SEM | Tukey's |
| Flv1+Fed9 | 429 | 0.350 | 0.229 | 0.011 | a |
| Flv3+Fed9 | 361 | 0.275 | 0.231 | 0.012 | b |
| | DF | Sum of Sq | Mean Sq | F Value | Prob>F |
| Model | 1 | 1.08 | 1.08 | 20.61 | **<0.0001** |
| Error | 788 | 41.48 | 0.05 | | |
| Total | 789 | 42.57 | | | |

| Figure S16A | One-sample T-te | null hypothesis: mean=1 | | | |
|---|---|---|---|---|---|
| **Flv1 soluble** | n | Mean | SD | SEM | P |
| light | 3 | 1.190 | 0.092 | 0.053 | **0.070** |
| CCCP | 3 | 0.666 | 0.112 | 0.065 | **0,035** |
| Two-Sample T-test, null hypothesis: difference in mean=0 | | | | | |
| light-CCCP | | 0.524 | | 0.084 | **0,003** |
| **Flv1 membran** | n | Mean | SD | SEM | P |
| dark | 4 | 0.222 | 0.213 | 0.107 | |
| light | 4 | 0.173 | 0.096 | 0.048 | |
| CCCP | 4 | 0.389 | 0.184 | 0.092 | |
| dark-light | | 0.049 | | 0.117 | **0.689** |
| dark-CCCP | | -0.167 | | 0.141 | **0.282** |
| light-CCCP | | -0.216 | | 0.104 | **0.083** |
| **Figure S16A** | **One-sample T-te** | null hypothesis: mean=1 | | | |
| **Flv2 soluble** | n | Mean | SD | SEM | P |
| light | 4 | 1.694 | 0.983 | 0.492 | **0.253** |
| CCCP | 4 | 0.489 | 0.366 | 0.183 | **0.068** |
| Two-Sample T-test, null hypothesis: difference in mean=0 | | | | | |
| light-CCCP | | 1.205 | | 0.524 | **0.061** |
| | **One-sample T-te** | null hypothesis: mean=1 | | | |
| **Flv2 membran** | n | Mean | SD | SEM | P |
| light | 4 | 0.721 | 0.379 | 0.190 | **0.237** |
| CCCP | 4 | 0.826 | 0.219 | 0.110 | **0.211** |
| Two-Sample T-test, null hypothesis: difference in mean=0 | | | | | |
| light-CCCP | | -0.105 | | 0.219 | **0.648** |
| **Figure S16A** | **One-sample T-te** | null hypothesis: mean=1 | | | |
| **Flv3 soluble** | n | Mean | SD | SEM | P |
| light | 3 | 1.006 | 0.043 | 0.025 | **0.83** |
| CCCP | 3 | 0.935 | 0.008 | 0.005 | **0,005** |
| Two-Sample T-test, null hypothesis: difference in mean=0 | | | | | |
| light-CCCP | | 0.071 | | 0.025 | **0,047** |
| **Flv3 membran** | n | Mean | SD | SEM | P |
| dark | 3 | 0.840 | 0.030 | 0.017 | |
| light | 3 | 0.769 | 0.236 | 0.136 | |
| CCCP | 3 | 0.724 | 0.136 | 0.078 | |
| dark-light | | 0.049 | | 0.117 | **0.635** |
| dark-CCCP | | -0.167 | | 0.141 | **0.222** |
| light-CCCP | | 0.046 | | 0.157 | **0.786** |

| **Figure S20D** | | **One-way ANOVA** | | | |
|---|---|---|---|---|---|
| strain | n | Mean | SD | SEM | Tukey's |
| neat | 4 | 0.439 | 0.027 | 0.014 | a |
| DTT | 3 | 0.429 | 0.046 | 0.026 | a |
| $CuCl_2$ | 3 | 0.603 | 0.028 | 0.016 | a |
| KCN | 3 | 1.078 | 0.231 | 0.133 | b |
| | DF | Sum of Sq | Mean Sq | F Value | Prob>F |
| Model | 3 | 0.9 | 0.290 | 22.75 | **0.0002** |
| Error | 9 | 0.1 | 0.013 | | |
| Total | 12 | 1.0 | | | |

| **Figure S17A** | | **One-way ANOVA** | | | |
|---|---|---|---|---|---|
| μM CCCP | n | Mean | SD | SEM | Tukey's |
| 0 | 3 | 0.493 | 0.070 | 0.040 | b |
| 15 | 2 | 0.636 | 0.141 | 0.100 | ab |
| 30 | 3 | 0.874 | 0.169 | 0.098 | a |
| | DF | Sum of Sq | Mean Sq | F Value | Prob>F |
| Model | 2 | 0.222 | 0.111 | 6.378 | **0.042** |
| Error | 5 | 0.087 | 0.017 | | |
| Total | 7 | 0.309 | | | |

| **Figure S21D** | | **One-way ANOVA** | | | |
|---|---|---|---|---|---|
| strain | n | Mean | SD | SEM | Tukey's |
| neat | 1099 | 0.077 | 0.177 | 0.005 | a |
| DTT | 820 | 0.072 | 0.156 | 0.005 | a |
| $CuCl_2$ | 961 | 0.038 | 0.150 | 0.005 | b |
| KCN | 936 | 0.065 | 0.143 | 0.005 | a |
| | DF | Sum of Sq | Mean Sq | F Value | Prob>F |
| Model | 3 | 0.9 | 0.289 | 11.57 | **<0.0001** |
| Error | 3812 | 95.3 | 0.025 | | |
| Total | 3815 | 96.2 | | | |

| **Figure S22D** | | **One-way ANOVA** | | | |
|---|---|---|---|---|---|
| strain | n | Mean | SD | SEM | Tukey's |
| neat | 1147 | 0.010 | 0.238 | 0.007 | a |
| DTT | 1358 | 0.004 | 0.233 | 0.006 | a |
| $CuCl_2$ | 873 | -0.032 | 0.254 | 0.009 | b |
| KCN | 924 | -0.055 | 0.281 | 0.009 | b |
| | DF | Sum of Sq | Mean Sq | F Value | Prob>F |
| Model | 3 | 2.9 | 0.972 | 15.61 | **<0.0001** |
| Error | 4298 | 267.6 | 0.062 | | |
| Total | 4301 | 270.5 | | | |

| **Figure S25D** | | **One-way ANOVA** | | | |
|---|---|---|---|---|---|
| strain | n | Mean | SD | SEM | Tukey's |
| neat | 1227 | 0.159 | 0.216 | 0.006 | d |
| high cation | 1290 | 0.470 | 0.174 | 0.005 | a |
| low cation | 808 | 0.219 | 0.278 | 0.010 | c |
| CCCP | 646 | 0.408 | 0.148 | 0.006 | b |
| | DF | Sum of Sq | Mean Sq | F Value | Prob>F |
| Model | 3 | 74.1 | 24.695 | 568.0 | **<0.0001** |
| Error | 3967 | 172.5 | 0.043 | | |
| Total | 3970 | 246.5 | | | |
